# Structure of Bifunctional Variediene Synthase Yields Unique Insight on Biosynthetic Diterpene Assembly and Cyclization

**DOI:** 10.1101/2024.12.03.626647

**Authors:** Eliott S. Wenger, David W. Christianson

## Abstract

An unusual family of bifunctional terpene synthases has been discovered in which both catalytic domains – a prenyltransferase and a cyclase – are connected by a long, flexible linker. These enzymes are unique to fungi and catalyze the first committed steps in the biosynthesis of complex terpenoid natural products: the prenyltransferase assembles 5-carbon precursors to form C_20_ geranylgeranyl diphosphate (GGPP), and the cyclase converts GGPP into a polycyclic hydrocarbon product. Weak domain-domain interactions as well as linker flexibility render these enzymes refractory to crystallization and challenge their visualization by cryo-EM. Despite these challenges, we now present the first experimentally-determined structure of a massive, 495-kD bifunctional terpene synthase revealing the assembly of all catalytic domains. The cryo-EM structure of variediene synthase from *Emericella variecolor* (EvVS) exhibits a bollard-like architecture, consisting of a hexameric prenyltransferase core sandwiched between two triads of cyclase domains. Although prenyltransferase and cyclase active sites are relatively close together, enzymological measurements indicate that GGPP is not channeled from one to the other. Surprisingly, however, the individual cyclase domain from another bifunctional diterpene synthase, fusicoccadiene synthase from *Phomopsis amygdali*, preferentially receives GGPP from the EvVS prenyltransferase in substrate competition experiments. Our previous studies of fusicoccadiene synthase suggest that GGPP channeling occurs through transient binding of cyclase domains to the sides of the prenyltransferase oligomer. The bollard-like architecture of EvVS leaves the sides of the prenyltransferase oligomer open and accessible, suggesting that a non-native cyclase could bind to the sides of the prenyltransferase oligomer to achieve GGPP channeling.

## Introduction

Terpenes represent the largest class of natural products and play many roles in nature, such as the deterrence of herbivores^1–2^ and the attraction of pollinators.^3^ Terpenes also have myriad industrial and medicinal uses, e.g., in pharmaceuticals,^4^ in flavors and fragrances,^5^ and in biofuels.^6^ The astonishing diversity of hydrocarbon skeletons in the terpenome is rooted in a single 5-carbon metabolite of primary metabolism, isopentenyl diphosphate (IPP).^7^ IPP can be isomerized to form dimethylallyl diphosphate (DMAPP),^8^ and then DMAPP can be combined with additional equivalents of IPP in iterative, head-to-tail condensation reactions to yield linear isoprenoids containing 5*n* carbons (*n* = 2, 3, 4…).^9^ These reactions are catalyzed in all domains of life by prenyltransferases that share a common α fold with conserved metal-binding motifs that coordinate to a catalytic Mg^2+^_3_ cluster.^10,11^

Linear isoprenoids serve as substrates for terpene cyclases that generate complex hydrocarbon scaffolds in a single enzymatic reaction with structural and stereochemical precision.^11,12^ A class I terpene cyclase shares the α fold of a prenyltransferase with certain variations in conserved metal-binding motifs. Coordination of the substrate diphosphate group to the Mg^2+^_3_ cluster triggers ionization to form inorganic pyrophosphate (PP_i_) and an allylic carbocation. The initially-formed carbocation then propagates through several intermediates in a mechanistic sequence typically resulting in a product containing multiple rings and stereocenters.^13–15^

While individual prenyltransferases and terpene cyclases are found in animals,^16–20^ plants,^3^ fungi,^21^ and bacteria,^22^ several fungal terpene synthases are quite unusual in that they contain both a prenyltransferase and a cyclase in a single polypeptide chain.^23–26^ Given that these enzymes catalyze sequential biosynthetic steps, they have been referred to as assembly-line terpene synthases.^27^ Bifunctional terpene synthases are refractory to crystallization due to the flexibility of the polypeptide linker connecting catalytic domains, but individual domains have yielded X-ray crystal structures.^28,29^ Cryo-EM studies of full-length bifunctional terpene synthases clearly reveal oligomeric prenyltransferase cores, but not all cyclase domains are visualized; indeed, most are characterized by weak, uninterpretable, or nonexistent density.^30–33^

For example, negative-stain EM and cryo-EM studies of fusicoccadiene synthase from *Phomopsis amygdala* (PaFS) reveal that cyclase domains are randomly splayed out around prenyltransferase oligomers; however, some particles are observed with one or two cyclase domains associated with the side of the prenyltransferase oligomer.^30,31^ Of note, the low resolution cryo-EM structure of a PaFS variant with a spliced-out linker shows all cyclase domains locked in place on the sides of the prenyltransferase oligomer;^34^ moreover, substrate channeling is definitively established in this variant and in wild-type PaFS.^34,35^ In another example, cryo-EM studies of glutaraldehyde-crosslinked macrophomene synthase from *Macrophomina phaseolina* reveal partial densities at low resolution corresponding to some cyclase domains above and below the prenyltransferase oligomer.^32^

Here, we report the cryo-EM structure of variediene synthase from *Emericella variecolor* (EvVS; **Figure 1**).^36^ Strikingly, all catalytic domains of the 495-kD bifunctional synthase are visualized: the hexameric prenyltransferase core is sandwiched between triads of cyclase domains above and below the prenyltransferase core, yielding a bollard-like assembly. While the proximity of prenyltransferase and cyclase domains in this assembly might suggest the possibility of geranylgeranyl diphosphate (GGPP) channeling between active sites, we show that no intramolecular substrate channeling occurs. Thus, GGPP is released from the prenyltransferase to solution before binding to the cyclase. Intriguingly, however, the individual cyclase domain of fusicoccadiene synthase (PaFS_CY_), *does* exhibit channeling with this system when added to EvVS reaction mixtures. In other words, GGPP is preferentially channeled from the EvVS prenyltransferase to the non-native cyclase PaFS_CY_ rather than being released to solution. Structural comparisons of EvVS and PaFS suggest a molecular rationale for intermolecular substrate channeling.

**Figure 1.**
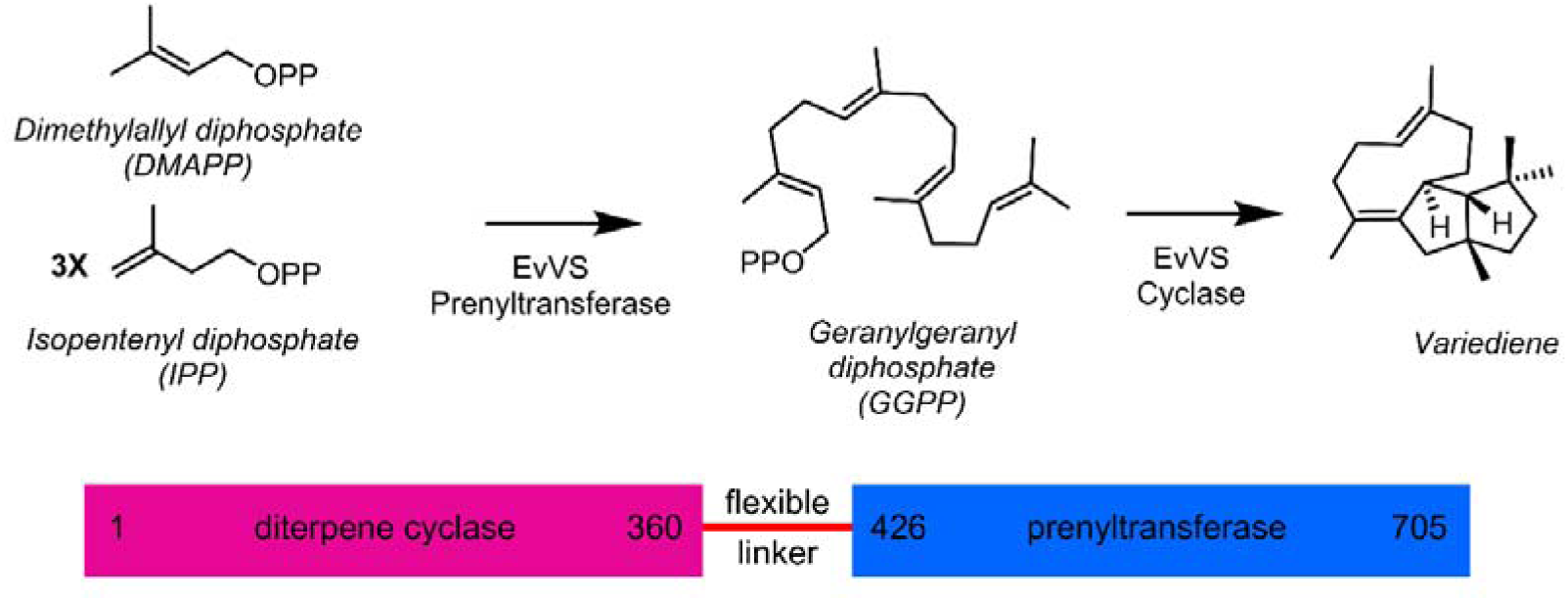
Reaction scheme and primary structure of EvVS.

## Results

### Catalytic Activity and Steady-State Kinetics

The catalytic activity of EvVS was first confirmed with substrate GGPP, which yields a single product previously established to be variediene.^36^ When incubated with the 5-carbon precursors DMAPP and IPP, variediene is similarly observed, establishing that both domains are catalytically active: the prenyltransferase domain synthesizes GGPP from DMAPP and 3 equivalents of IPP, and the cyclase domain converts GGPP into variediene (**Figure S1A**). No other cyclization products are observed. We also purified a separate construct comprising only the EvVS cyclase domain (EvVS_CY_). When incubated with GGPP, this construct exhibits similar high-fidelity variediene synthase activity, with no substantial generation of alternative diterpene products or any other product from any other isoprenoid substrate (**Figure S1B**). When EvVS_CY_ is incubated with GGPP and a range of divalent cations, maximal variediene generation is observed only with Mg^2+^ (**Figure S1C**). Similar results were reported with the initial discovery and characterization of full-length EvVS.^36^ Additionally, steady-state kinetics of full-length EvVS and EvVS_CY_ reported herein show that the turnover number (*k*_cat_) of GGPP cyclization by EvVS_CY_ is only reduced by 50% compared with full-length EvVS (**Figure S2**). Steady-state kinetic parameters measured for both constructs are comparable to those previously measured for PaFS.^34^ Cooperativity was not observed for either construct.

To ascertain the possibility of substrate channeling between the prenyltransferase and cyclase domains of EvVS, substrate competition experiments were performed in similar fashion to those described in our recent study of PaFS.^34^ When an equimolar mixture of EvVS and cyclooctatenol synthase (CotB2) is incubated with GGPP, the variediene:cyclooctatenol product ratio is 1:1.1 (**Figure 2A**, **Table 1**). When the same enzyme mixture is incubated with DMAPP and IPP, such that the only GGPP available for cyclization is that generated by the EvVS prenyltransferase, the variediene:cyclooctatenol product ratio changes only slightly to 1:0.7. When an equimolar mixture of EvVS and spatadiene synthase is incubated with GGPP, and then in a second experiment with DMAPP and IPP, the variediene:spatadiene product ratio is essentially unchanged at 1:0.6 and 1:0.5, respectively (**Figure 2B**, **Table 1**). If there were GGPP channeling in full-length EvVS, more variediene would be generated in these experiments when incubated with DMAPP and IPP compared with GGPP, i.e., more of the GGPP generated by the EvVS prenyltransferase would remain on the enzyme for cyclization. Therefore, GGPP channeling does not occur in EvVS. This contrasts with the results of substrate competition experiments with PaFS, where substantial fusicoccadiene enrichment is observed when incubated with DMAPP and IPP compared with GGPP.^34^

**Figure 2.**
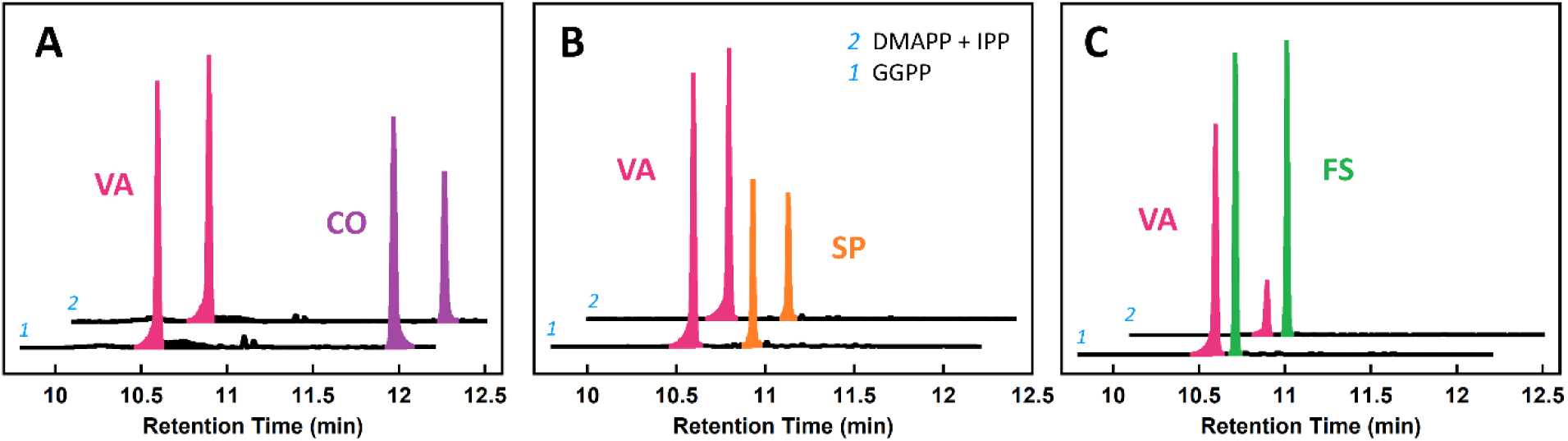
Interrogation of substrate channeling. (**A**) When an equimolar mixture of EvVS and CotB2 is incubated with GGPP, variediene (VA) and cyclooctatenol (CO) are generated at a 1:1.1 ratio (trace 1). When incubated with DMAPP and IPP instead, only a small change is observed in the product ratio (trace 2). (**B**) When an equimolar mixture of EvVS and spatadiene synthase is incubated with GGPP, variediene (VA) and spatadiene (SP) are generated at a 1:0.6 ratio (trace 1). When incubated with DMAPP and IPP instead, no substantial change is observed in the product ratio (trace 2). (**C**) When an equimolar mixture of EvVS and PaFS_CY_ is incubated with GGPP, variediene and fusicoccadiene (FS) are generated at a 1:1.1 ratio. When incubated with DMAPP and IPP instead, the variediene:fusicoccadiene product ratio is 1:5. Substantial enrichment of fusicoccadiene indicates that the GGPP generated by the EvVS prenyltransferase is preferentially delivered to PaFS_CY_.

**Table 1.** Substrate Competition Assay.

| Equimolar Enzyme Mixture | Products Formed<sup>a</sup> | Product Ratio from GGPP | Product Ratio from DMAPP and IPP |
| --- | --- | --- | --- |
| EvVS, CotB2 | VA:CO | 1 : 1.1 ± 0.2 | 1 : 0.7 ± 0.1 |
| EvVS, SpS | VA:SP | 1 : 0.6 ± 0.1 | 1 : 0.5 ± 0.1 |
| EvVS, PaFS<sub>CY</sub> | VA:FS | 1 : 1.1 ± 0.1 | 1 : 5.0 ± 0.4 |

<sup>a</sup>VA, variediene; CO, cyclooctatenol; SP, spatadiene; FS, fusicoccadiene.

When an equimolar mixture of EvVS and the cyclase domain of PaFS (PaFS_CY_) is incubated with GGPP, a variediene:fusicoccadiene product ratio of 1:1.1 is observed; when incubated with DMAPP and IPP, the variediene:fusicoccadiene ratio is 1:5 (**Figure 2C**). Surprisingly, the substantial enrichment of fusicoccadiene indicates that GGPP generated by the EvVS prenyltransferase preferentially transits to the non-native PaFS_CY_ cyclase domain rather than the native cyclase domain of EvVS. Thus, the three-dimensional (3D) structure of EvVS structure must be able to accommodate transient association of PaFS_CY_ to enable preferential GGPP transfer. Our recent studies of PaFS indicate that transient cyclase-prenyltransferase association supporting GGPP channeling occurs at the side of the prenyltransferase oligomer.^30,31,34^ Accordingly, we hypothesize that PaFS_CY_ similarly associates with the side of the EvVS prenyltransferase oligomer for preferential GGPP transfer.

### Cryo-EM Structure Determination

To ascertain the mode of prenyltranferase-cyclase association in EvVS, we determined the cryo-EM structure of the full-length enzyme. Protein samples were incubated with Mg^2+^ and the unreactive substrate analogue geranylgeranyl thiolodiphosphate (GGSPP). Following cryo-EM data collection, initial two-dimensional (2D) classification revealed several orientations of a well-defined hexameric assembly with all catalytic domains visualized (**Figure S3**). Initial particle sorting and *ab initio* 3D reconstructions confirmed that cyclase domains associate with the top and bottom of the hexameric prenyltransferase core (**Figure S4**). Heterogenous refinement was used for initial sorting of 637,378 particles. While the majority of particles (398,696; 63%) exhibit densities corresponding to all six cyclase domains, 146,937 particles (23%) are missing density for one cyclase domain and 17,383 particles (3%) are missing densities for two cyclase domains. Weak or absent densities for one or two associated cyclase domains suggest that these domains are randomly splayed-out and hence averaged out in 3D reconstructions. Of note, particles with two missing cyclase densities align to only 9.9 Å resolution, indicating that high-quality particles have at least five cyclase domains bound.

Overall, cyclase-prenyltransferase association appears to be much more stable in full-length EvVS compared with all other full-length bifunctional terpene synthases previously studied by cryo-EM.^30–33^ Even so, the observation of EvVS particles containing only 4 or 5 ordered cyclase domains suggests that prenyltransferase-associated cyclase positions are in dynamic equilibrium with splayed-out cyclase positions.

For 3D refinement, all available particles were used first to determine the structure of the hexameric prenyltransferase core with densities for cyclase domains subjected to particle subtraction. Density is visible beginning at L426, and ends at G710 (portions of the final C-terminal helix (P712-V725) were masked out during particle subtraction). After local refinement, prenyltransferase particles aligned to 2.77 Å resolution (**Figure 3A, B**). Only one polypeptide loop (E620-D630) lacks density and is judged to be disordered, and therefore not modeled, in each monomer. The hexameric prenyltransferase can be described as a trimer of dimers and aligns closely with the structures of hexameric prenyltransferases from other bifunctional fungal diterpene synthases (**Figure 3C**).^28,29,33,34^ The complete workflow is summarized in **Figure S5** with relevant statistics recorded in **Table S1**. Data coverage and Gold-Standard Fourier Shell Correlation (GSFSC) plots are presented in **Figures S6** and **S7**.

**Figure 3.**
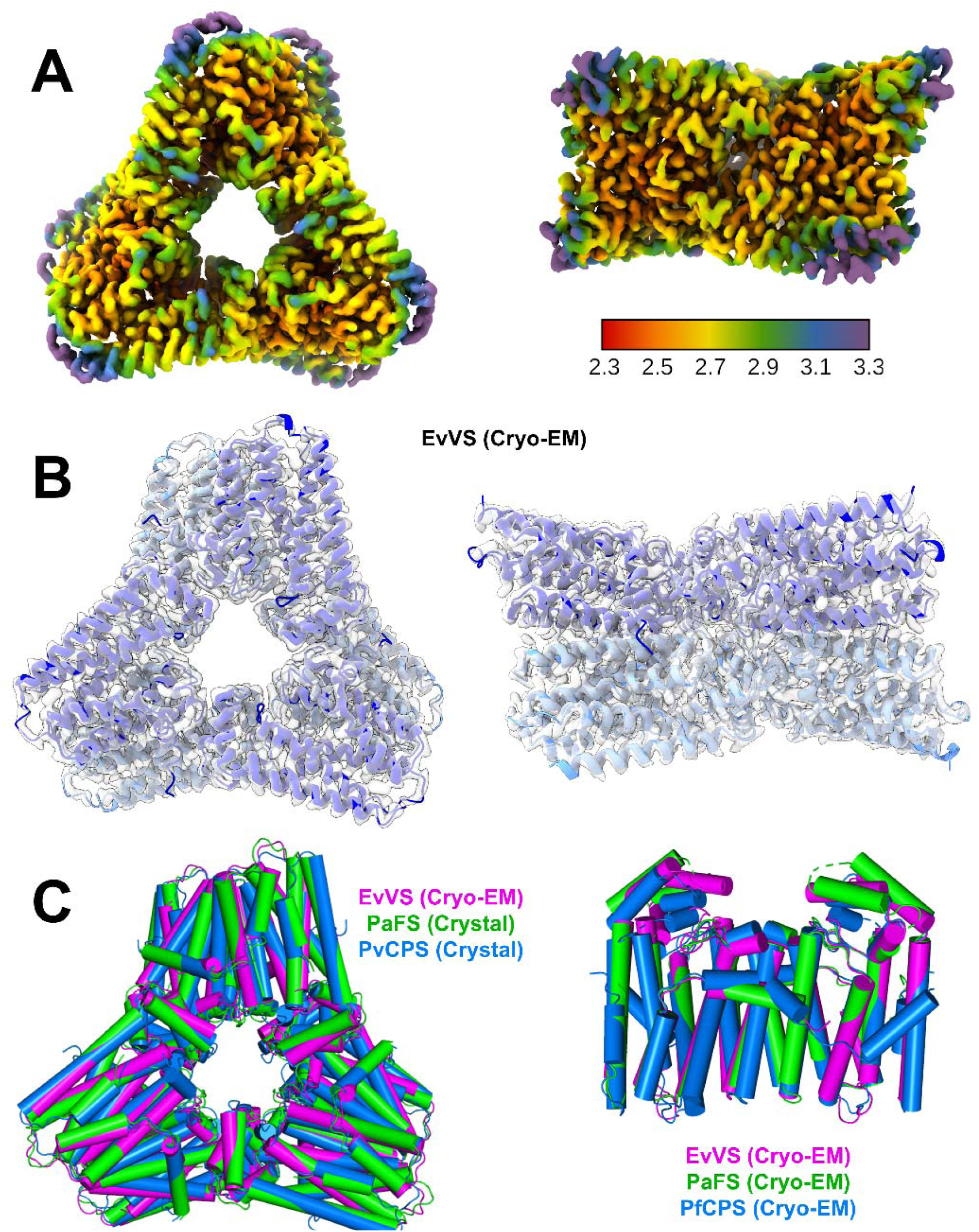
Structure of the EvVS hexameric core and comparison with similar structures. (**A**) The overall resolution of the EvVS hexamer is 2.77 Å; interior regions have higher resolutions and external loops have lower resolutions. (**B**) The model of the EvVS prenyltransferase is fit into the map with *C*3 symmetry. (**C**) The EvVS prenyltransferase hexamer as well as a single dimer overlay closely with structures of other fungal prenyltransferases determined by X-ray crystallography (left; PDB 5ERO and 6V0K, respectively) and cryo-EM (right; PDB 8EAX and 8V0F, respectively). Copalyl diphosphate synthase from *Penicillium verruculosum*, PvCPS; copalyl diphosphate synthase from *Penicillium fellutanum*, PfCPS.

Structures of hexameric prenyltransferase-cyclase assemblies containing six and five cyclases were then determined. Using the particle stacks identified above, 3D classification was used to find the particles in each subset aligning with the highest resolution. The six-cyclase structure is based on 60,587 particles that aligned to 3.52 Å resolution. Phenix confirmed the presence of *C*3 symmetry and the overall resolution after symmetry-averaging improved to 3.18 Å (**Figure 4A**). While density for the prenyltransferase core is sufficiently strong to define orientations for most side chains, densities for cyclase domains are sufficient only to define α-helices. Even so, the orientation of each cyclase domain is unambiguously determined, allowing for the construction of a model of the entire assembly using cyclase coordinates generated with AlphaFold^37^ (**Figure 4B**). In similar fashion, the five-cyclase structure is based on 51,215 particles that aligned to 3.59 Å resolution (**Figure 4C**). Here, too, density for the prenyltransferase core is sufficiently strong to define orientations for most side chains, whereas densities for cyclase domains are sufficient only to define α-helices and the orientation of each domain (**Figures 4D**). The complete workflow is summarized in **Figure S5**, and relevant statistics are recorded in **Table S1**, **Figure S6**, and **Figure S7**.

**Figure 4.**
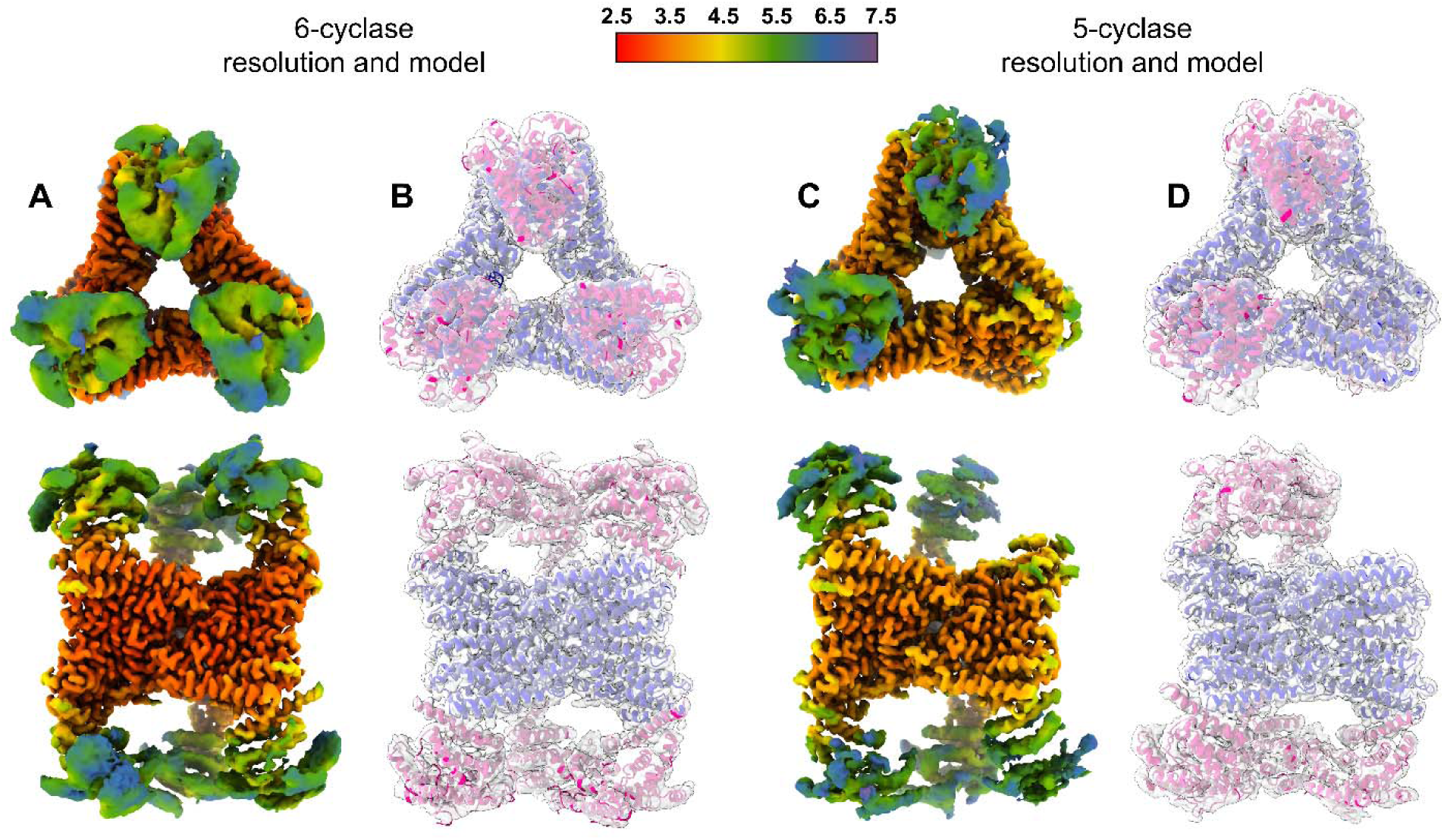
Six-cyclase and five-cyclase structures of EvVS. (**A**) Map showing the six-cyclase structure of EvVS. (**B**) In the corresponding model of this structure, the prenyltransferase domains are blue and the cyclase domains are magenta. (**C**) Map showing the five-cyclase structure of EvVS. (**D**) Model of the structure with the same color scheme outlined in (**B**).

The 66-residue flexible linker connecting the prenyltransferase and cyclase domains of EvVS is not visible in cryo-EM maps, but the C-terminus of the cyclase and the N-terminus of the prenyltransferase are sufficiently well defined in the six-cyclase structure to suggest how they are most likely linked together (**Figure 5A**). The maximum length of an extended 66-residue linker would be approximately 230 Å; two different prenyltransferase domain N-termini are within this distance from the C-terminus of each cyclase. One of these connections is more direct and the other would require the linker to partially wrap around the cyclase or the interior of the assembly. Accordingly, the more direct connection is judged to be more likely, yielding a domain-swapped architecture (**Figure 5B**).

**Figure 5.**
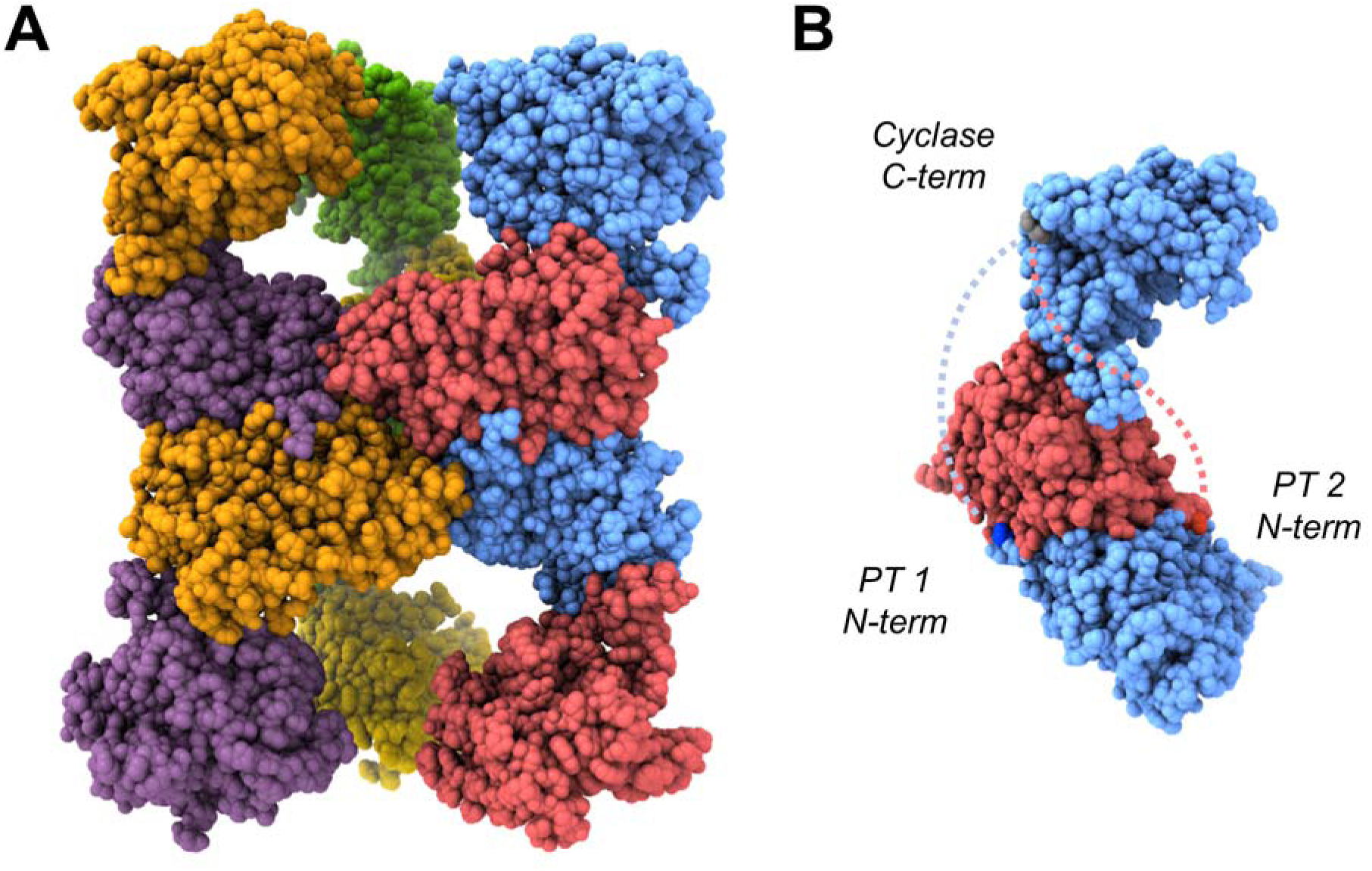
Domain swapped architecture of EvVS. (**A**) The more direct prenyltransferase-cyclase linker connection yields a domain-swapped architecture; each connected domain pair appears as a single color. (**B**) Support for domain-swapping derives from the locations of domain termini: the C-terminus of the blue cyclase is grey, the N-terminus of prenyltransferase 1 (PT 1, blue) is dark blue, and the N-terminus of prenyltransferase 2 (PT 2, red) is dark red. The blue dotted line connects the cyclase with prenyltransferase 1, which is a more direct connection. The red dotted line connects the cyclase with prenyltransferase 2, which is a less direct connection that would require the linker to partially wrap around the assembly. The chains shown in **B** are rotated approximately 60° clockwise around a vertical axis from their position in **A** so that both N-termini are visible.

Notably, cyclase resolution drops with increasing distance from the core in both the six-cyclase and five-cyclase structures. However, a single helix of the cyclase always appears with well-defined density as it interacts with the prenyltransferase. Moreover, the densities of more distant helices are distorted, suggesting that the cyclase domain wobbles back and forth with the prenyltransferase-associated helix serving as a pivot point. The application of 3D conformational variability analysis reveals that cyclase wobbling is captured in the dataset (**Movies 1** and **2**).

To resolve these conformations and to determine the structure of the cyclase domain with the highest possible resolution, the entire set of 491,079 particles was subjected to symmetry expansion and particle subtraction to remove all cyclases except the one defined by the strongest density in each particle. After local refinement, we used 3D classification with a focus mask over the cyclase domain. Particle subtraction and refinement worked well only if the entire prenyltransferase core was retained along with the single cyclase domain, which yielded a particle of sufficient size to achieve effective alignment. Ultimately, three classes emerged at 3.00 Å, 3.08 Å, and 2.98 Å overall resolution showing the cyclase in slightly different orientations illustrating the extent of cyclase wobble (**Figure 6**). The complete workflow is summarized in **Figure S5**; relevant statistics are found in **Table S1**, **Figure S6**, and **Figure S7**.

**Figure 6.**
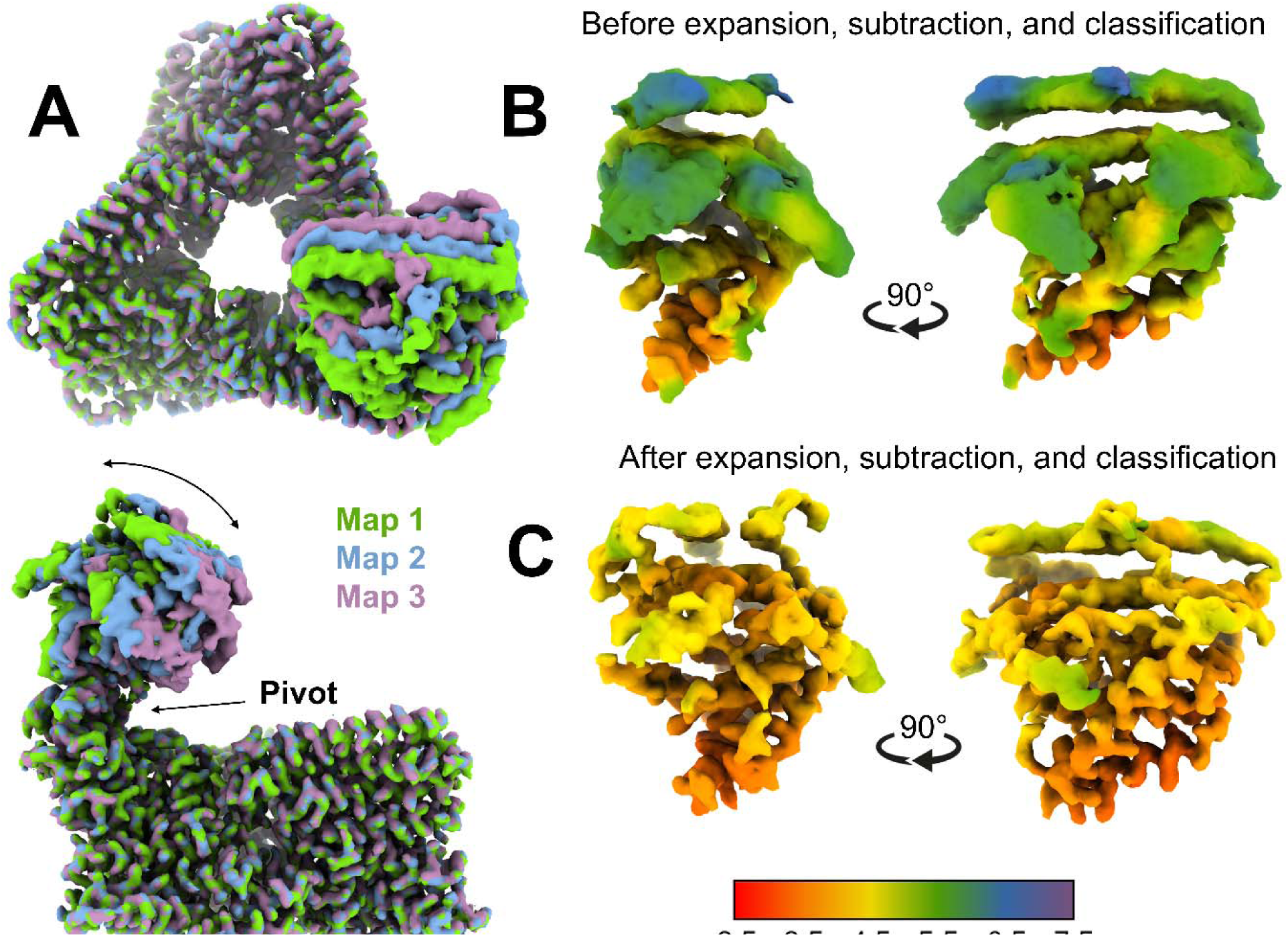
Highest-resolution view of the EvVS cyclase domain. (**A**) 3D classification led to three different cyclases conformations illustrated in Maps 1, 2, and 3, the average of which yields the distorted densities observed in the six- and five-cyclase structures. These maps illustrate the extent of cyclase wobble. (**B**) Representative cyclase domain map with distorted density. (**C**) Cyclase domain map showing substantial resolution improvement resulting from symmetry-expansion and particle subtraction.

In the best map of the EvVS cyclase domain (Map 3, 2.98 Å resolution), the model begins at residue D9 and ends with Q343. While several loops are disordered and not modeled for the cyclase domains in the six- and five-cyclase structures, overall densities in these three models are sufficiently improved such that only one disordered loop is not modeled (V122– R146). This segment contains two helix termini and a loop connecting them adjacent to the active site (**Figure 7A**). The overall structure overlays nearly perfectly with the AlphaFold-predicted structure except for the disordered loop (AlphaFold designates this loop as a segment of low prediction confidence (**Figure S8**)). The cryo-EM structure of the cyclase domain also displays strong agreement with the structure of GJ1012 synthase from *Fusarium graminearum* (FgGS), a fungal cyclase exhibiting the highest sequence identity with EvVS (40%) for which a crystal structure is available.^38,39^

**Figure 7.**
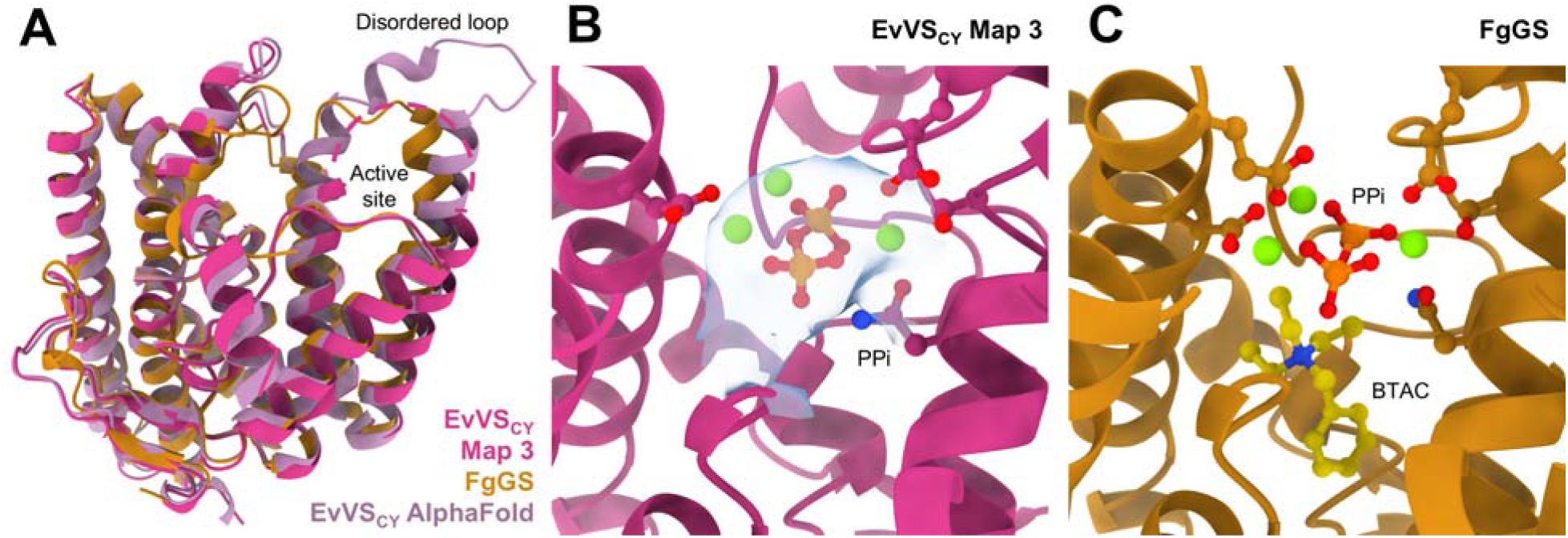
Structure of the EvVS cyclase domain. (**A**) Overlay of the cryo-EM structure of the cyclase domain from Map 3 (magenta), the AlphaFold structure prediction (purple), and the crystal structure of FgGS (yellow). (**B**) Close-up view of the active site of the EvVS cyclase domain. Density is shown by a light blue surface for cyclase Map 3 (EMD-47457, contour level 0.090). (**C**) The Mg^2+^_3_ -PP_i_ cluster in the active site of FgGS (PDB 6VYD) binds similarly to that modeled in the EvVS cyclase active site in (**A**). Also bound in the FgGS active site is the benzyl triethylammonium cation (BTAC).

Additional densities are observed in the active sites of all three cyclase conformations, located between conserved metal binding motifs characteristic of terpene cyclases: the DXXXD/E motif (D120 and E124) and the NXXXSXXXE motif (N250, S254, and E258). The strongest density is observed in Map 3 and is consistent with the binding of a Mg^2+^_3_-PP_i_ cluster in similar fashion to that observed in the active site of FgGS (**Figure 7B,C**). Maps 1-3 contain uninterpretable patches of density extending into the hydrophobic active site cavity that may correspond to the 20-carbon isoprenoid tail of GGSPP. However, if GGSPP were hydrolyzed during the time course of sample preparation, these patches of density would be spurious, leaving only the binding of the Mg^2+^_3_-PP_i_ cluster.

The structure of the cyclase-prenyltransferase interface is visualized at approximately 3.8 Å resolution, which is sufficiently high to model side chains involved in domain-domain interactions. A single α-helix from each domain mediates these interactions – P712-V725 in the prenyltransferase and R129-S150 in the cyclase (**Figure 8A**). The total buried surface area at the interface is 1,228 Å^2^, consistent with a stable protein-protein interaction. No obvious hydrogen-bonding interactions are evident to confer stability; instead, helix-helix association appears to be driven mainly by “knobs-into-holes” packing^40^ and the hydrophobic effect. The three most important residues on the prenyltransferase that form the binding surface, W714, L718, and H721 (**Figure 8B**), are only loosely conserved in other bifunctional terpene synthases (**Figure S9**), so it is not clear if a similar, stable domain-domain interaction can form in these other systems.

**Figure 8.**
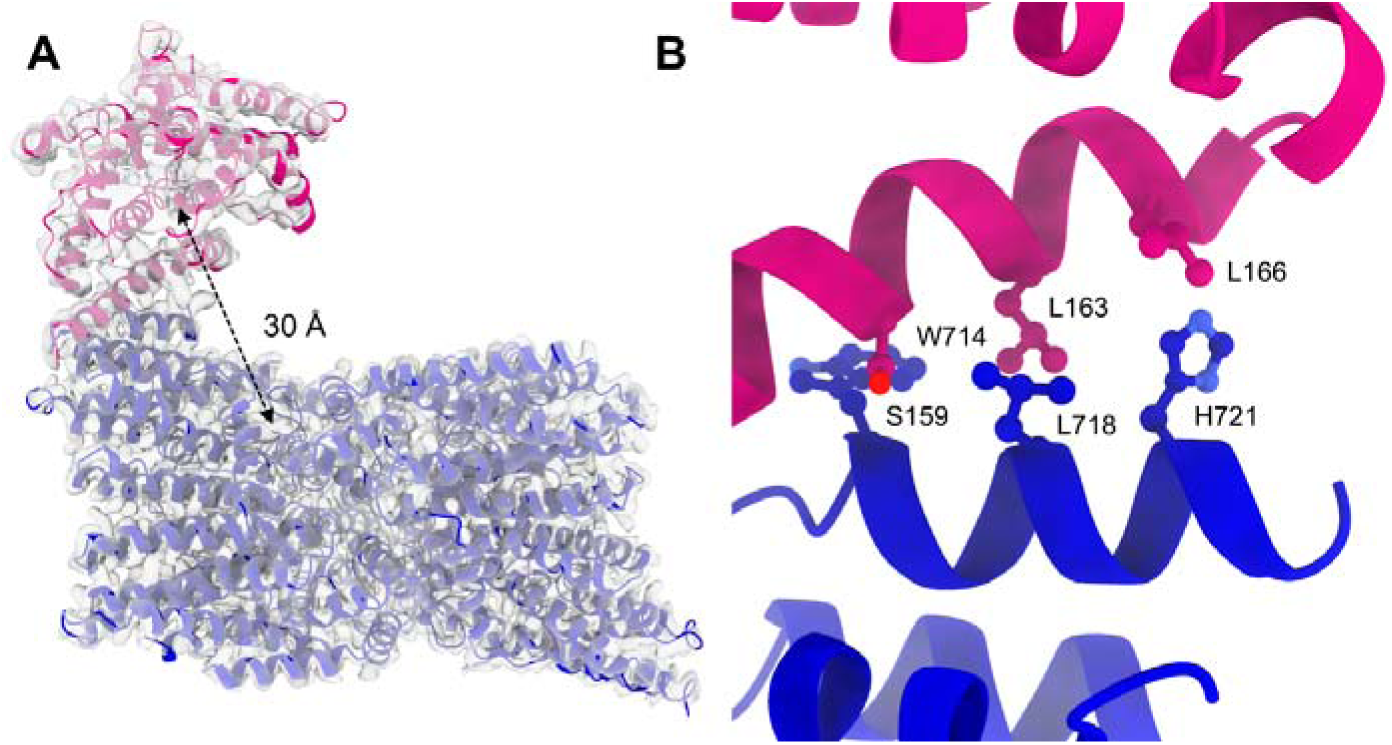
Active site separation and domain-domain interface. (**A**) The structure of the cyclase (magenta) interacting with the prenyltransferase core (blue) reveals an active site separation of approximately 30 Å. (**B**) Three residues from each domain define the domain-domain interface.

As observed in all terpene cyclases, active site residues are predominantly hydrophobic. The proposed cyclization mechanism of EvVS (**Figure 9A**)^36^ requires an active site base to quench the final carbocation intermediate by deprotonation, and the PP_i_ co-product likely serves this function. A plausible model of the precatalytic enzyme-substrate complex shows that GGPP can bind in the active site with a conformation supporting the two ring closure reactions in the first steps of the cyclization cascade (**Figure 9B**). A computational study of variediene formation by Hong and Tantillo shows that after substrate ionization, the C1-C11 and C10-C14 coupling reactions are likely concerted.^41^ The model of GGPP binding in the cyclase active site (**Figure 9b**) roughly corresponds to the calculated conformation of the initially-formed geranylgeranyl cation described in this study.

**Figure 9.**
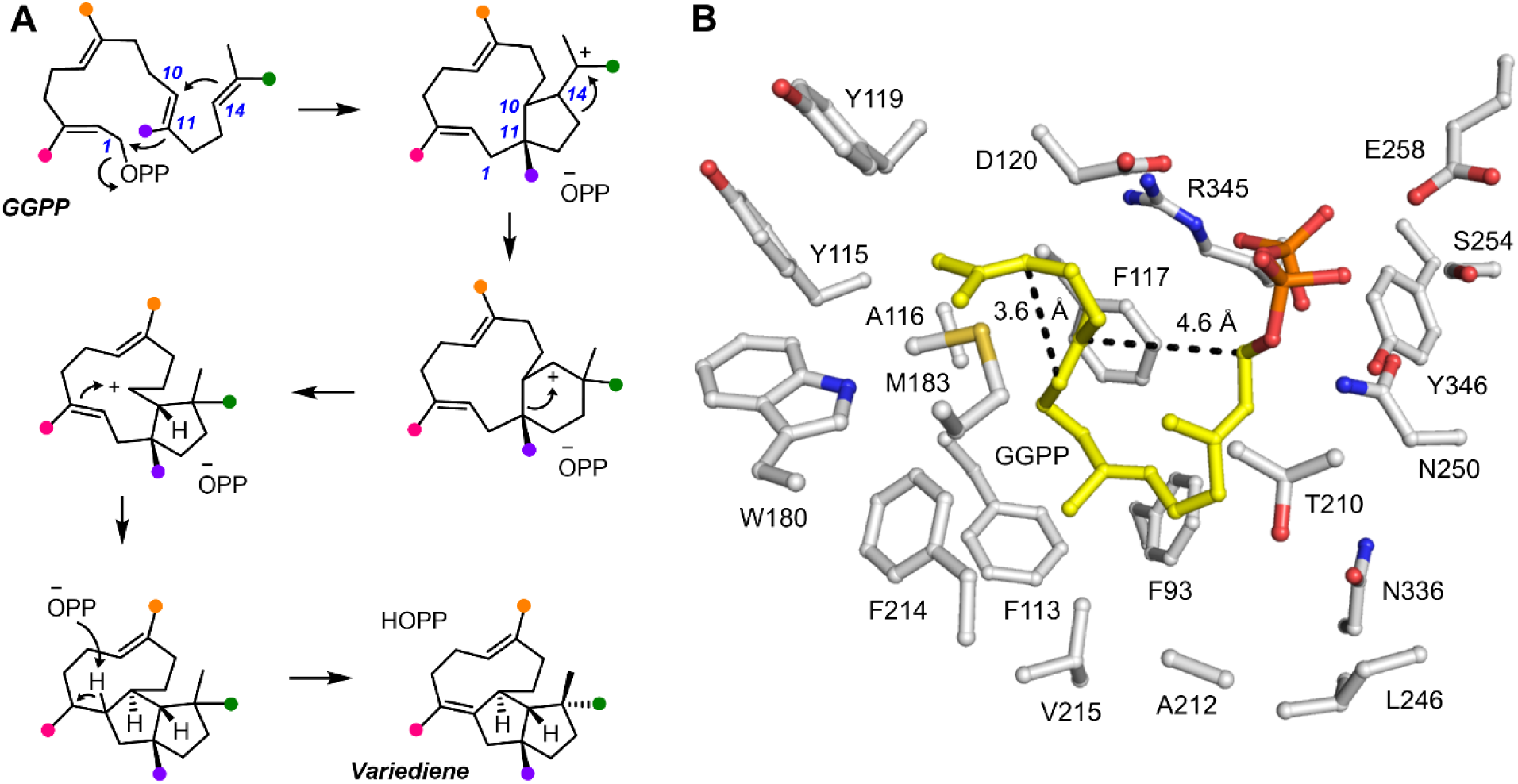
Cyclization cascade. (**A**) Mechanism of GGPP cyclization proposed by Abe and colleagues.^36^ Isoprenoid methyl groups are color-coded to better follow the mechanistic sequence. (**B**) Model of the precatalytic binding conformation of GGPP in the active site of the EvVS cyclase (atomic coordinates from Map 3) consistent with the ring closure reactions outlined in (**A**).

## Discussion

The cryo-EM structure of EvVS is the most complete structure of a bifunctional terpene synthase reported to date, providing the highest-resolution view ever achieved of a prenyltransferase oligomer with all associated cyclase domains. Catalytic domains are positioned in the oligomer so as to make a domain-swapped architecture most likely. Given that the prenyltransferase-cyclase active site separation is 30 Å (**Figure 8A**), it is surprising that intramolecular GGPP channeling is not observed, but intermolecular GGPP channeling is observed with PaFS_CY_. Since all six cyclase domains are locked in place at the top and bottom of the EvVS oligomer, we speculate that PaFS_CY_ associates with the side of the oligomer to enable preferential GGPP transfer (**Figure S10**).

To complement the newly-determined structure of EvVS, we identified sequences for other class I bifunctional fungal terpene synthases and performed systematic structural predictions using AlphaFold (**Table S2** and **Figures S11-S21**). Each prediction yields atomic coordinates for five different models with a predicted template modeling (PTM) score and interface predicted template modeling (iPTM) score based on discrepancies in the five models. A high confidence prediction generates five very similar models and yields a high iPTM score. We performed an AlphaFold prediction for each sequence five times, with six copies of the sequence entered to allow hexamer formation, to generate an average iPTM score from five distinct predictive trials. Scores ranged between 0.69 and 0.43; sequences with high scores generally yield five predictions very similar to each other, whereas sequences with low scores yield multiple predicted domain-domain binding modes. EvVS yields the highest iPTM score of 0.69, consistent with the stable interdomain interaction observed in the cryo-EM structure of this system. PaFS yields a score of only 0.43, consistent with weaker and transient domain-domain interactions as observed in cryo-EM studies.^30,31^ Thus, AlphaFold appears to have useful predictive power in determining whether a bifunctional terpene synthase features stable prenyltransferase-cyclase interactions or not.

Side-on interactions of the cyclase domain with the hexameric prenyltransferase core are very rarely predicted by AlphaFold: only nine examples are observed in 110 total predictions. Three examples are observed for PaFS, the only system with experimental evidence for a side-on interaction;^30,31^ two examples are observed with deoxyconidiogenol synthase, and one example is observed with talarodiene synthase. In these predicted structures, all cyclases interact with the sides of the prenyltransferase core. The remaining three examples are observed with ophiobolin F synthase, where some but not all cyclase domains are directly or diagonally associated with the sides of the prenyltransferase core. The remaining 101 predictions show cyclases interacting with the top and bottom of the respective hexameric prenyltransferase cores, in a manner similar to that observed in the cryo-EM structure of EvVS. Notably, these systems exhibit higher sequence identity across the 22 enzymes studied (**Figures S22** and **S23**). AlphaFold may thus have utility in prospecting for systems that are capable of achieving transient, side-on cyclase-prenyltransferase binding as well as systems with stable top-and-bottom cyclase-prenyltransferase interactions. Our results with EvVS as well as our previously reported results with PaFS^34^ suggest that substrate channeling is more likely to occur in systems capable of side-on cyclase-prenyltransferase binding. These results will guide our continuing explorations of substrate management in bifunctional terpene synthases, as well as protein engineering approaches aimed at optimizing biosynthetic efficiency in such systems.

## Methods

### Protein Expression and Purification

All proteins described herein were prepared in similar fashion. Full sequences for the EvVS (Uniprot identifier A0A0P0ZD79) and EvVS_CY_ construct are listed in the Supporting Information. For each protein, a plasmid containing the target gene in the pet28A vector fused to an N-terminal His_6_ tag (twenty additional residues added) was acquired from Genscript and transformed into BL21-DE3 competent cells (New England Biolabs). Following overnight incubation at 37°C on agar, a single colony was used to inoculate a culture supplemented with 50 µg/mL kanamycin. The culture was then grown overnight at 37°C shaking at 150 RPM. The starter culture was then transferred into six 1-L flasks with Luria-Bertani media, each containing 50 µg/mL kanamycin. When the OD_600_ reached 1.0, flasks were cooled on ice for 45 min prior to induction of protein expression by the addition of 0.5 mM isopropyl-β-D-1-thiogalactopyranoside. Cultures were then shaken at 16°C for 18 h shaking at 150 RPM.

The next morning, cells were harvested by centrifugation and resuspended in lysis buffer [50 mM HEPES (pH 7.5), 200 mM NaCl, 2 mM TCEP, and 10% glycerol]. The suspension was stirred for 2 h at 4°C. Cell lysis was achieved by sonication (1 s pulse on and 3 s off at 30% power for 40 minutes) and the lysate was clarified via centrifugation. Clarified lysate was applied to a pre-equilibrated Ni-NTA column (Cytiva) and washed with five column volumes of lysis buffer. Protein was then eluted using the same buffer supplemented with 300 mM imidazole. Fractions containing the target protein were pooled and immediately loaded onto a size-exclusion column (GE Healthcare) equilibrated with lysis buffer, skipping any concentration or dialysis steps. Fractions containing the target protein were combined and diluted to a total volume of 100 mL with zero-salt lysis buffer (lysis buffer without NaCl). This solution was then applied to an anion-exchange column (Cytiva), washed with five column volumes of zero-salt buffer, and eluted using a 100-mL gradient between zero-salt and high-salt lysis buffer (lysis buffer containing 1 M NaCl). Protein fractions were combined and the glycerol concentration was adjusted to 30% by addition of a solution of 75% glycerol in water (the amount of 75% glycerol solution spiked in was equal to one quarter the volume of the pooled fractions). The solution was then concentrated using an Amicon centrifugal filter (MWCO 30,000 kDa) until the protein concentration reached approximately 120 µM. The protein was flash-frozen in liquid nitrogen and stored at -80° C.

### Steady-State Kinetics

Steady-state kinetic parameters were determined using the EnzChek^TM^ Pyrophosphate Assay Kit (ThermoFisher Scientific) as previously described for PaFS and other terpene synthases.^29,30^ Assays were performed on a 100-µL scale with the protein concentration at 0.5 µM in lysis buffer. Purine nucleoside phosphorylase, inorganic pyrophosphatase, MgCl_2_, and 2-amino-6-mercapto-7-methylpurine ribonucleoside were added to final concentrations of 0.1 U, 0.003 U, 2.5 mM, and 200 µM, respectively. Each reaction was individually assembled in a cuvette to a final volume of 90 µL. The desired concentration of GGPP (ranging from 2.5 to 50 µM) was added from the 0.5 mM working dilution (a 1:10 dilution in lysis buffer of the 5 mM stock prepared by reconstitution of solid in 70% methanol, 30% 10 mM NH_4_HCO_3_ in water). Addition of 10 µL of enzyme from a 5 µM stock initiated the reaction and the cuvette was immediately placed into a Cary 60 UV/vis spectrometer. The absorbance at 360 nm was measured every 2 s for 2 min. In total, two identical trials for each substrate concentration were performed for each enzyme.

The initial velocity of each reaction was determined by fitting the linear portion of each trace (40–80 s) using linear regression in Prism software. Slopes were converted into rates (rise in AU/s^−1^ to nM phosphate/s^−1^) using a conversion factor (0.0258) determined in a separate control experiment wherein pyrophosphate was added directly to the coupled-enzyme assay. Plotting of the rates over GGPP concentration allowed for analysis in Prism software. The data were fit using the Michaelis-Menten equation:

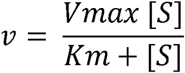

### Substrate Competition Assay

In a typical substrate competition experiment, both proteins were diluted to 3.0 µM in freshly-prepared lysis buffer with a total volume of 200 µL in a 2-mL glass vial with no insert. MgCl_2_ was added to a final concentration of 3 mM. To initiate the reaction, 20 µL of substrate was added – either 20 µL GGPP from a 5 mM stock, or 10 µL of DMAPP from a 10 mM stock and 10 µL of IPP from a 30 mM stock. All substrates were resuspended in a mixture of 70% methanol, 30% 10 mM NH_4_HCO_3_ in water, and adding substrate in the specified amounts from the specified stock concentrations ensured that each sample contained the same final volume of methanol regardless of which substrates were added. The capped vial was incubated for 4 h on the benchtop at room temperature. Ethyl acetate (200 µL) was then added and the mixture was vortexed for 5 s. The entire mixture was then transferred to a 1.7 mL Eppendorf tube and centrifuged at 6,000 RPM for 5 s. The organic layer (200 µL) was then removed using a micro-pipette and transferred to a fresh GC vial with an insert.

Gas chromatography-mass spectrometry (GC-MS) was used for the identification and quantification of diterpene cyclization products. Samples were run on an Agilent 8890 GC system coupled to a 5977C GC/MSD mass spectrometer with a J&W HP-5MS GC column capillary column in EI+ mode. The following temperature program was used: hold at 60°C for 2 min, ramp to 320°C at 10°C/min, and hold at 320°C for 2 min. A solvent delay of 4 min was used. Variediene eluted at 10.5 min, fusicoccadiene eluted at 10.7 min, spatadiene at 10.9 min, and cyclooctatenol at 11.9 min.

### Cryo-EM Structure Determination

Prior to grid preparation, EvVS was dialyzed into cryo-EM buffer [25 mM HEPES (pH 7.5), 100 mM NaCl, 60 mM sodium phosphate monobasic, and 140 mM potassium phosphate dibasic] at 4°C for 1 h. MgCl_2_, geranylgeranyl thiodiphosphate (GGSPP), and 3-([3-cholamidopropyl]dimethylammonio)-2-hydroxy-1-propanesulfonate (CHAPSO) protein detergent were then added to final concentrations of 2.5 mM, 20 mM, and 8 mM, respectively. The final protein concentration was approximately 100 µM. The sample was filtered by centrifugation in a 0.22-µM filtration unit (Millipore). R1.2/1.3 200-mesh copper grids (Quantifoil) were glow-discharged for 2 min (easiGlow, Pelco). 3 µL protein sample was applied to the grid, the grid was blotted for 3 s with blot force of 3, and another 3 µL of protein sample was applied and blotted (multiple protein application and blotting steps can increase particle density dramatically^42^). The grid was then flash-frozen in liquid ethane using a Mark IV Vitrobot (ThermoFisher Scientific). Grids were prepared at 100% humidity. Frozen grids were clipped and transferred to a Titan Krios G3i cryogenic transmission electron microscope (Thermo Fischer Scientific) operating at 300 keV at Brookhaven National Laboratory. Images were recorded with a K3 Summit electron detector at 81,000 magnification (1.07 Å/pixel) with a defocus of -1.0 to -3.0 µM. The beam intensity was 15 e^−^/px/s (13 e^−^/Å^2^/s) and the exposure time was 3.84 s. The total dose was 50 e^−^/Å^2^/s. 40 frames were taken. A total of 8,727 movies was collected from a single grid.

The EvVS dataset was processed using cryoSPARC (version 4.4.1)^43^ and the workflow of all 3D reconstructions is summarized in **Figure S5**. After motion and CTF correction, 106 exposures were rejected during curation to yield 8,621 micrographs. Topaz Extract (box size of 384 Å) was used to generate a pool of 844,262 particles that were subjected to 2D classification (150 classes). Selection of all protein-resembling classes yielded a pool of 421,245 particles. After *ab initio* 3D reconstruction and heterogenous refinement, the best model (115,628 particles) was used with 983 of the micrographs to train a Topaz model. The model was then used for re-extraction of the entire dataset, yielding a pool of 1,228,533 particles. 2D classification of these particles allowed for selection of 807,870 particles for further processing.

With these particles, we generated *ab initio* 3D reconstructions of four different classes and subjected them to four rounds of heterogenous refinement. After the refinement, one class (83,058 particles) contained only junk. The remaining 724,812 were reference motion corrected. Heterogenous refinement was used to exclude an additional 87,134 junk particles and to sort the remaining 637,678 particles into three stacks: a stack with all six cyclases present (398,696 particles), with only five cyclases present (164,320 particles), and with only four cyclases present (17,383). The four-cyclase class was subjected to homogenous refinement and then non-uniform refinement to generate the map shown in **Figure S4**.

To obtain the structure of the hexameric core, the five-cyclase and six-cyclase particle stacks were combined by homogenous refinement (491,079 particles) and then subjected to non-uniform refinement (wherein *C*3 symmetry was applied). The cyclase domains were subjected to particle subtraction and the resulting map was locally refined. The map shown in **Figure 3** was polished with DeepEMhancer^44^ and has an overall resolution of 2.77 Å.

To determine the six-cyclase structure, the stack of 398,696 particles was subjected to 3D classification with a focus mask covering all six cyclase domains. The best class that emerged had 60,587 particles and a resolution of 3.63 Å after homogenous refinement. Non-uniform refinement was used to apply *C*3 symmetry and the resolution improved to 3.18 Å. The map of the six-cyclase structure shown in **Figure 4A** was polished with DeepEMhancer.^44^ To determine the five-cyclase structure, the stack of 164,320 particles was subjected to 3D classification with a focus mask covering five cyclase domains. The best class had 51,215 particles and an overall resolution of 3.67 Å. After non-uniform refinement, the resolution improved to 3.59 Å. The map of the five-cyclase structure shown in **Figure 4B** was polished with DeepEMhancer.^44^ 3D variability analysis on the same pool of 398,696 particles produced **Movie 1**. We used UCSF ChimeraX^45^ to generate movie (.mov) files showing the transition across the set of 20 maps from different orientations, and converted the .mov files to GIFs using a freely available online tool (ezgif.com).

To determine the highest-resolution structure of the cyclase domain, the combined pool of five- and six-cyclase particles with applied *C*3 symmetry described previously was symmetry expanded and locally refined to give a pool of 1,473,237 particles. Five cyclases were subtracted, leaving the entire hexameric prenyltransferase core and a single cyclase domain. 3D classification into five classes with a focus mask over the cyclase domain produced two lower-resolution classes and a series of three classes with 298,050, 294,185, and 289,027 particles with 3.0, 3.08, and 2.98 Å resolution overall (after local refinement) revealing clear density for the cyclase domain in three slightly different positions relative to the core (designated Map 1, Map 2, and Map 3, respectively). The maps shown in **Figure 6** were polished with DeepEMhancer.^44^

### Model Building and Refinement

Atomic coordinates generated by AlphaFold^37^ were used as starting points for all six models. UCSF ChimeraX 1.3 was initially used to dock the models into the maps.^45^ Iterative rounds of refinement and model building were then performed with Phenix Real Space Refine^46^ and WinCoot.^47^ Model quality was assessed with MolProbity.^48^

The final model of the EvVS prenyltransferase hexamer (PDB 9E2H, EMD-47452) contains 6 polypeptide chains and no ligands or water molecules. Chain A contains residues 426-710. Chain B contains residues 425-710. Chain C contains residues 425-709. Chain D contains residues 425-710. Chain E contains residues 426-709. Chain F contains residues 425-710. One loop is disordered and not modeled due to insufficient density, comprising residues 620-630, in all six chains.

The final model of EvVS with six cyclases (PDB 9E2I, EMD-47453) contains 6 polypeptide chains and no ligands or water molecules. Chain A contains residues 28-724. Chain B contains residues 24-724. Chain C contains residues 27-724. Chain D contains residues 25-724. Chain E contains residues 27-724. Chain F contains residues 24-725. Some residues are not modeled due to insufficient cryo-EM density. In chain A, the unmodeled residues are 35-42, 51-57, 122-144, 189-195, 317-320, 361-425, and 620-630. In chain B, the unmodeled residues are 36-41, 123-144, 190-203, and 362-425. In chain C, the unmodeled residues are 36-42, 51-55, 122-139, 186-204, 363-425, and 619-631. In chain D, the unmodeled residues are 37-39, 51-52, 122-144, 190-196, 362-426, and 620-630. In chain E, the unmodeled residues are 36-40, 51-52, 122-144, 189-201, and 361-426. In chain F, the unmodeled residues are 37-39, 51-52, 122-144, 190-196, and 363-425.

The final model of EvVS with five cyclases (PDB 9E2J, EMD-47454) contains 6 polypeptide chains and no ligands or water molecules. Chain A contains residues 27-725. Chain B contains residues 27-725. Chain C contains residues 28-724. Chain D contains residues 27-724. Chain E contains residues 28-724. Chain F contains residues 426-724. Some residues are not modeled due to insufficient cryo-EM density. In chain A, the unmodeled residues are 124-142, 362-426, and 620-630. In chain B, the unmodeled residues are 36-41, 52-53, 57-61, 83-88, 125-146, 189-201, 277-289, 358-425, and 620-630. In chain C, the unmodeled residues are 36-41, 51-55, 125-144, 189-194, 362-425, and 619-630. In chain D, the unmodeled residues are 38-42, 125-143, 190-196, 318-320, 365-425, and 620-627. In chain E, the unmodeled residues are 37-40, 125-146, 190-208, 240-244, 266-270, 275-284, 310-319, 359-425, and 619-628. In chain F, the unmodeled residues are 620-628.

The final model of EvVS with one cyclase designated Map 1 (PDB 9E2K, EMD-47455) contains 6 polypeptide chains, one ligand chain, and no water molecules. Chain A contains residues 426-725. Chain B contains residues 26-698. Chain C contains residues 425-709. Chain D contains residues 425-701. Chain E contains residues 426-709. Chain F contains residues 425-710. Chain G contains the ligands: three magnesium ions (MG; 1-3) and a pyrophosphate ion (POP; 4). Some residues are not modeled due to insufficient cryo-EM density. In chain A, the unmodeled residues are 620-630. In chain B, the unmodeled residues are 123-144, 361-424, 452-459, and 620-630. In chain C, the unmodeled residues are 620-630. In chain D, the unmodeled residues are 452-457 and 620-630. In chain E, the unmodeled residues are 620-630. In chain F, the unmodeled residues are 620-630.

The final model of EvVS with one cyclase designated Map 2 (PDB 9E2L, EMD-47456) contains 6 polypeptide chains, one ligand chain, and no water molecules. Chain A contains residues 426-723. Chain B contains residues 26-698. Chain C contains residues 426-709. Chain D contains residues 427-701. Chain E contains residues 427-709. Chain F contains residues 427-709. Chain G contains the ligands: three magnesium ions (MG; 1-3) and a pyrophosphate ion (POP; 4). Some residues are not modeled due to insufficient cryo-EM density. In chain A, the unmodeled residues are 620-630. In chain B, the unmodeled residues are 123-145, 361-425, 452-459, and 620-630. In chain C, the unmodeled residues are 620-630. In chain D, the unmodeled residues are 452-457 and 620-630. In chain E, the unmodeled residues are 620-630. In chain F, the unmodeled residues are 620-630.

The final model of EvVS with one cyclase designated Map 3 (PDB 9E2M, EMD-47457) contains 6 polypeptide chains, one ligand chain, and no water molecules. Chain A contains residues 426-723. Chain B contains residues 26-698. Chain C contains residues 426-709. Chain D contains residues 427-701. Chain E contains residues 427-709. Chain F contains residues 27-709. Chain G contains the ligands: three magnesium ions (MG; 1-3) and a pyrophosphate ion (POP; 4). Some residues are not modeled due to insufficient cryo-EM density. In chain A, the unmodeled residues are 620-630. In chain B, the unmodeled residues are 124-148, 361-425, 452-459, and 620-630. In chain C, the unmodeled residues are 620-630. In chain D, the unmodeled residues are 452-457 and 620-630. In chain E, the unmodeled residues are 620-630. In chain F, the unmodeled residues are 620-630.

## Acknowledgments

We thank Kollin Schultz and Ronen Marmorstein for guidance and advice with cryo-EM studies, and we thank the Beckman Center for Cryo-EM (RRID: SCR_022375) at the University of Pennsylvania for instrumentation access in the initial stages of this work. We also thank the Laboratory for BioMolecular Structure (LBMS) at Brookhaven National Laboratory for the use of cryo-EM data collection facilities; the LBMS is supported by the DOE Office of Biological and Environmental Research (KP1607011). Finally, we thank the NIH for grant GM56838 in support of this research.

## Competing Interests

The authors declare no competing interests.

## Data Availability

Atomic coordinates have been deposited in the Protein Data Bank (PDB, www.rcsb.org) and cryo-EM maps have been deposited in the Electron Microscopy Data Bank (EMD) with accession codes as follows: EvVS prenyltransferase hexamer, PDB 9E2H, EMD-47452; EvVS hexamer with six cyclase domains bound, PDB 9E2I, EMD-47453; EvVS hexamer with five cyclase domains bound, PDB 9E2J, EMD-47454; EvVS hexamer with one cyclase domain bound: Map 1, PDB 9E2K, EMD-47455; Map 2, PDB 9E2L, EMD-47456; Map 3, PDB 9E2M, EMD-47457.

## Notes

### Competing Interest Statement

The authors have declared no competing interest.

