## Supporting Information for "Structure of Bifunctional Variediene Synthase Yields Unique Insight on Biosynthetic Diterpene Assembly and Cyclization"

**EvVS DNA Sequence.** Full sequence of the EvVS expression construct. The $His_6$-tag adds twenty residues to the N-terminus (underlined).

ATGGGCAGCAGCCATCATCATCATCATCACAGCAGCGGCCTGGTGCCGCGCGGCAGCCATATGTCACAA
AGTTCTGATTTTATACTAAATTCCACCCTGAGTAGCGTTGTAGAGCGCTCAACCCCGGATATCGCGGGTT
TTTGCAGCGGTTATGAGCTCCGTCGCCACCACCATGAACATTTGGCGAATGAGGGCTCCTTGCGTTGCC
GTACTGACTGGGAACAGTTCATTGGTCCAATTGAACGTTGGGGTAGCTGCAATCCGTGGGAAGGTCATT
TTGGTGCTGTGGTGTTGCCGTTCTGCAAGCCGGAGCGCTTGGCGGTCATTTGCTACATTTTCGAATATGC
TTTTCTGTATGATAATGTGGTGGAAAGCGCGGCGAAGTCTACCCTGAACCTGAACACCGACAACATCGC
GCTGGACGAGACCGAGTATCGTACCGTGCGCTCCATTTTAGGAACTAAGCAGATTCAGAGCAAAATGTT
GCTGGAGCTGCTGTCTATCGACGCACCACGTGCAGAGGTGGTTATCAACTCTTGGAAAGAAATGATCTC
CACCACCGCAAAGAAAGATAAGACACGGGCATTCAACAACCTGGAAGAATACGTGGATTATAGAATCAT
TGATACCGGTGCTCCGTTTGTCGACATGCTGATGCGTTTCGGCATGGGCATTATGCTGACCCAAGAAGAA
CAAAAACGCATCGAGCCGATTGTCAAACCGTGTTATGCAGCGTTGGGCCTCGCGAACGATTACTTCTCC
TTCGACATCGAGTGGGAAGAATTTCAGGCGGAGAGCGACAAGACGACCATGACCAATGCGGTTTGGTT
ATTTATGCAGTGGGAAAACCTTAACGCGGAGCAGGCCAAGCGCCGTGTCCAAGAGGTTACCAAACAAT
ACGAGCAACAATATCTGCGTAACATCGCGGATTTTGCAGCGGGCGAAGGTAAAGAAAACATCAAGCTG
CAAACGTACCTGAAGGCGCAGGGTTACCAGGTTCCAGGTAATGTTGCATGGAGCTTGCGTTGCCCGCGT
TACCATCCGTGGCTGTGTAAGGAGGCCGCGAGCCTGCTGCACCAGGATACCATTCAGGAACTGGAGGC
CGGGCGTAAACCGCAGGCGCTGGAAGAGTACCGCAGCCGTTCGCACTCCGAGTCCGACTTGTCTGATG
CTAGCCCGACGTTTTGGAGCGGAAGCTGTCGTTCGTCCGCGCGTTCTTCGGTTAGCAGCGCCTTCGGCC
CCCCGGACAAGGATATCAGCATTACCCCGGCGATTCTGGGCGATGAGCACTTGCTGGGTCCGGCTGAGT
ACATCTCTTCTCTGCCAAGCAAAGGCGTCAGAGAAGCGTTTATCGATGGTCTAAACGTATGGCTGGTCC
TGCCGGACCATCGTGTTAATCAGTTGAAGTCCATCGCCCAAACCCTCCACAACGCTTCCCTGATGCTCG
ACGATATCGAGGATCACAGCCCGCTGCGTCGGGGCCGTCCGAGCACCCACATGATTTTCGGTACGGAAC
AAACGATTAACAGCGCTAATTTCCTGCTGATCGACGTGATGGAAAAAGTTCGTCAGCTGGACGATCCGC
GCTGCATGGACATCTATTTGGAGGAGATGAGAAACCTGTTCATTGGCCAGTCTTTTGATCTGTATTGGAC
CCGTAACGGCGAGTGTCCGAGCGAAGAGCAATACCTGGACATGATCCGCCAGAAGACGGGCGGTCTGT
TTCGTCTGCTAACTCGCATGATGGTGCAAATTGCGCCGGTTCAACAAAAAGGCCTGGAAACCCAGCTAG
CCTCTCTGAGCGACGTGCTTGGCGAGTTCTTCCAGGTGCGCGATGATTACAAAAACCTGACCGAAGAGT
ACACCGGTCAAAAGGGCTTCTGCGAAGACCTGGACGAATGCAAGTTTTCCTATCCGCTTATCCATGCCC
TGACCAGCCAGCCGAAAAATGTTCAGTTGCGTGGTATTTTGCAGCAGTCTCGTAGCGCAGGTGGTTTGG
ACGTTCCGCTGAAGGAAACGGTGCTGAGCCATCTGCGCCAAGCTGGTTCCATTGAGTACACCGAGGCG
AAAATGGGTGAACTCATGGAAAAAATCACCGACAGCGTTGTGAGCCTGGAGGGCGAGACTGGCTCTCC
GAATTGGGTTGTTCGTCTGCTGATCCACCGTTTGAAGGTGTAA

**EvVS$_{CY}$ DNA Sequence.** Full sequence of the EvVS$_{CY}$ expression construct. The His$_6$-tag adds twenty residues to the N-terminus (underlined).

ATGGGCAGCAGCCATCATCATCATCATCACAGCAGCGGCCTGGTGCCGCGCGGCAGCCATATGTCACAA
AGTTCTGATTTTATACTAAATTCCACCCTGAGTAGCGTTGTAGAGCGCTCAACCCCGGATATCGCGGGTT
TTTGCAGCGGTTATGAGCTCCGTCGCCACCACCATGAACATTTGGCGAATGAGGGCTCCTTGCGTTGCC
GTACTGACTGGGAACAGTTCATTGGTCCAATTGAACGTTGGGGTAGCTGCAATCCGTGGGAAGGTCATT
TTGGTGCTGTGGTGTTGCCGTTCTGCAAGCCGGAGCGCTTGGCGGTCATTTGCTACATTTTCGAATATGC
TTTTCTGTATGATAATGTGGTGGAAAGCGCGGCGAAGTCTACCCTGAACCTGAACACCGACAACATCGC
GCTGGACGAGACCGAGTATCGTACCGTGCGCTCCATTTTAGGAACTAAGCAGATTCAGAGCAAAATGTT
GCTGGAGCTGCTGTCTATCGACGCACCACGTGCAGAGGTGGTTATCAACTCTTGGAAAGAAATGATCTC
CACCACCGCAAAGAAAGATAAGACACGGGCATTCAACAACCTGGAAGAATACGTGGATTATAGAATCAT
TGATACCGGTGCTCCGTTTGTCGACATGCTGATGCGTTTCGGCATGGGCATTATGCTGACCCAAGAAGAA
CAAAAACGCATCGAGCCGATTGTCAAACCGTGTTATGCAGCGTTGGGCCTCGCGAACGATTACTTCTCC
TTCGACATCGAGTGGGAAGAATTTCAGGCGGAGAGCGACAAGACGACCATGACCAATGCGGTTTGGTT
ATTTATGCAGTGGGAAAACCTTAACGCGGAGCAGGCCAAGCGCCGTGTCCAAGAGGTTACCAAACAAT
ACGAGCAACAATATCTGCGTAACATCGCGGATTTTGCAGCGGGCGAAGGTAAAGAAAACATCAAGCTG
CAAACGTACCTGAAGGCGCAGGGTTACCAGGTTCCAGGTAATGTTGCATGGAGCTTGCGTTGCCCGCGT
TACCATCCGTGGCTGTGTAAGGAGGCCGCGAGCCTGCTGCACCAGGATTAA

**Bifunctional terpene synthase sequences used for multiple sequence alignments and AlphaFold predictions.** Enzyme acronyms are defined in Table S2; linker segments are underlined.

>EvVS (UniProt: A0A0P0ZD79)
MSQSSDFILNSTLSSVVERSTPDIAGFCSGYELRRHHHEHLANEGSLRCRTDWEQFIGPIERWGSCNPWEGH
FGAVVLPFCKPERLAVICYIFEYAFLYDNVVESAAKSTLNLNTDNIALDETEYRTVRSILGTKQIQSKMLLELL
SIDAPRAEVVINSWKEMISTTAKKDKTRAFNNLEEYVDYRIIDTGAPFVDMLMRFGMGIMLTQEEQKRIEPI
VKPCYAALGLANDYFSFDIEWEEFQAESDKTTMTNAVWLFMQWENLNAEQAKRRVQEVTKQYEQQYLR
NIADFAAGEGKENIKLQTYLKAQGYQVPGNVAWSLRCPRYHPWLCKEAASLLHQDTIQELEAGRKPQALE
EYRSRSHSESDLSDASPTFWSGSCRSSARSSVSSAFGPPDKDISITPAILGDEHLLGPAEYISSLPSKGVREAFI
DGLNVWLVLPDHRVNQLKSIAQTLHNASLMLDDIEDHSPLRRGRPSTHMIFGTEQTINSANFLLIDVMEKVR
QLDDPRCMDIYLEEMRNLFIGQSFDLYWTRNGECPSEEQYLDMIRQKTGGLFRLLTRMMVQIAPVQQKGL
ETQLASLSDVLGEFFQVRDDYKNLTEEYTGQKGFCEDLDECKFSYPLIHALTSQPKNVQLRGILQQSRSAGG
LDVPLKETVLSHLRQAGSIEYTEAKMGELMEKITDSVVSLEGETGSPNWVVRLLIHRLKV

>MpMS (UniProt: K2SUY0)
MCNTKCYNTLAKMTVITEPAMEYMYSVPLDESEYDKCGFCQDPRYRPRRHKDQHLARAGSAKAKELCEA
LIGVYPRPTCESAVGHSIALVMPECMPGRVEAMGEFMESIFYMDNIAESGSQQDTGNLGTEWANDMETGPT
TSVNSNTGAKQVMAKLALQLLSIDPVCAGNVMKAWKEWAAGFAKPRRFDSIEQYIDYRLVDSGAIVAVHL
MNFGMGLDISVEELREVSDIVNHAGKALSYQNDFFSFNYEHDMFVKLPDSIGIANAVFVLAETEGLSLAEA
KERVKELAKEHEDAVLRLKDEVESKVSYKLRICLEGLVDMVVGNLVWSASCDRYSSYRREKHQMELPIRIQ
GPPTPPQEPVYEKATLPNGKQLDAPTESSGKDLSDGVATLSGDEPVLGDEIVSAPIKYLESLPSKGFREAIIDG
MNGWLNLPARSVSIIKDVVKHIHTASLLCDDIEDSSPLRRGQPSAHIIFGVSQTVNSTSYLWTLAIDRLSELSS
PKSLRIFIDEVRKMQIGQSFDLHWTAALQCPSEEEYLSMIDMKTGGLFHLLIRLMIAESPRKVDMDFSGLVS
MTGRYFQIRDDLSNLTSEEYENQKGYCEDLDEGKYSLPLIHALKHTKNKVQLESLLIQRKTQGGMTLEMKR
LAIQIMKEAGSLEHTRKVVLELQDAVHRELAKLEEAFGQENYVIQLALERLRIKA

>CoSS (UniProt: A0A8F4SNK4)
MASEMWKYSSPIDPEVVKATGCFTTLPVRINNRDDLANGASSRVLKDWAEHTGNQSIDRNRVSFSPVGSFC
SLIYCETIPERLDSISYLTDLFFLIDDATEEVENDTLAQEQWAGFSGAMTDSLAEAAPQRDHDLDMMKKKKL
VARVMLDFMRLDPVLGLDLVKSCKAGWTPLTAGVEWETMEDYLRFRRLSAGLDIYWTKTVFGLGEKLTD
DEEKLIRPLTWAAEKAAMLNNDYWSWDIEYLQAGGNIEKLTNAVAVLMRKEGLTAEEGKQRIKNLIIGYEE
EYSRLRDDFYNAHPSARLYLRKRVELAGSMAAGVSFWSANSPRYHIPTQQTETTTESGEAAQPAAGTWHA
EVRDTSRPGSDSSVSSTSESDDSIDASASSGTTTTTTTTTTALSSSQTSLSSSLSEAEEEEEAHKPPPCFDAPAK
LGRAAIDAPIDYVSGMPSKGVRTSLIDAMNQWCRVPSAQLGAVKRVVDVLHNSSLILDDIQDDSPMRRGKT
ATHLVFGAAQAINSATFLHVRAVREVHATGSAALMAVLLEELEDLHVGQSWDLYWKYNLRWPTEDEYFSM
IDLKTGGLFRMLVRMMRVLAPEPTGGETKGGEFACDALVSMVSRFFQVRDDYLNLNSREYGSQKGWCED
LDEGKFSYLVIHCLETSPRFRDRVMGLFRQRAGTASASSGPTPMPSVAKVQIIEYLYEAGSFDACWRLLVRLE
DDIEGEIRRLEEATGEENPQMHLLLKLLSVKNDKPNKGPVVVPAGL

>CsSS (NBCI: MW685620)
MLEANELYPYSVAVDRDEVVQSGALTSLPVRIHRYNHLADAGALCLTNDWRCTMKDGQDRKSNGSPCVV
GNWGSFIWPESRPERLGLLCYLLDAGCFHDDACEEMPIAAAHQEHLDLDAAMDVEDRRELSSDSRSLRTK
QLISNAVLECIKVDRVGAMRMLEAYRKKWLRIMETYNTEEINTVEDYFLARANNGGMGAYYAMLEFSLGI
LVTDEEYEMMAEPIAHVERCMLLTNDYWSWPRERKQAEYQEAGKVFNIVWFLKKIELCTEEEAVSKVRD
MVHAEERNWTAAKTRLYSQFGNLRQDLVKFLENLHTALAGNDYWSSQCYRHNDWEHIPDLPGEDAPKLH
ELATLGRRL[illegible]ADSARFTECKPSVGTVSGDETQSLPGRSTSGEESASGYMYSISSA
SAPPSPSKEHQSSYPSIPYQSSAHVAGDLDSPVLREPIKYIRNMPSKNLRTQLIDCFNIWLNASGPAISVIKEVI
DCLHHSSLILDDIEDGSHLRRGFPATHVVYGTCQAVNSATFLYVQAVESVHAAARNNPEMMDVFLKHLRQL
FNGQSWDLYWTYHRQCPTEEQYLDMVDQKTGAMLQLLVGLMQTAQPQHPGKVGGVVHSEVLFRFTQLF
GRFFQVRDDYMNLTSTDYARQKGFAEDLDEQKFSYMIVHMYQRYPEAKDKVEGVFRAMQQGGISQVAAD
TSKRYILSILDETGSTAATKALLLKWHDEITEEIGALERHFGVDNALLRLLVETLRV

>ZbSS (UniProt: A0A0F4GLU2)
MAEFAIPVPDDVVKQSGTLSRFPTAVHREHARCLAAANKIRDDFAAQVDWDLDAKTTGHYPTLGAVHVVA
FTMPECLPERLALMTRFTDFTIMNDDYYDAVDRDQATSFNAELQRSLGRDCHSNTVQGNASVAIKTKQFQA
SILVEMMVMDRDLAMDVMDTYSDGLETATFPPSDICTIEEYLPVRLVNCGLDVFQEMSCFGLGVHLTKAEK
EKLSDIANTALYTAALINDCHSWPKELKHHLETPGSDVPFNAVCILMRQFNCSDVKAIERLRAIYVEIQERHL
SLVRNLEQSEGSIPETHRKYIMAAQYAASGSEFWSLYAPRYPSKEDLEQPEYVLVDNVLHRRSMSDKDLPTS
DKDLARADSAMHIETIKTAGSSGMSHMNEAYSSTPATEMVAWDAGSEIIHTEIDSNGSKELAPNGAQTRVQK
PSEDAVRAPYDYIRALPSKRIRETFIDALDSWLAVPAGSSASIKSIIGMLHQSSLMLDDIEDDSTLRRGKPTAH
TLFGIAQTINSANWVFACAFEELRSLRGVDAATIFVEEVKNLHCGQALDLHWKHHTYIPSVDEYLNMVDH
KTGGLFRLCVRLMQGESSTSCHHIDAERFITLLGRYFQIRDDYQNLVSDEYTNQKGFCEDLDEGKISLPLIYC
LAGSDPTQIMIRGILQHKRAGEMPLSMKKLILEKMRSGGALNATISLLKDLQDNILEELKSLESAFGSGNPM
LELVLRRLWI

>PfVS (UniProt: A0A8F4PPI6)
MAGTRSSRPGATSFIQHSIPLPRSAYEGVEYFCRFRPRIHRDAILADAGSWQCQVDFFGSSATARADSIRNKN
HTSYAVGCINPVVGNFTALCACEAIPDRLALTTYMVEYAYIHDDVIEYAENKDEDRDNVRRRQLQAKMAVE
LMDIDKVKGKECLRLWKEMSDVFVQIRELKFTKLDDYLTFRVIDAGCPWTMSLLCFSMDFTLNDDEVEKT
AAITSAAYDGWVLVNDYFSWEKEWKNHQANGGSGVIANAIFLFMRWYSVDAVEGRRMLRKEILAREEKY
CKAKEEFLVSGNVTDKTSQWLELLDHVTAGNFAWSMTTARYQLGGKDAYPALRAANTDNWETSTTDSLS
NPISHNADKIARKINLIFKEQKFLDARGLVNHTEDYPPIVLTAQVSQPDETPEFIHSQVTQARSFTQYEKMILQ
PQNYLESMPSKGVRNSVIDGLEMWYQVPERSLATIRKIVNLLHSSSLMLDDIEDNSPLRRGLPATHTVFGISQ
TINSANLLMFKALKAAESLSPAAVRIFIERIIEGHIGQGMELYWTFHTEIPTEEEYFVMVDGKTGGLFILLAEL
MRSEATRHKDLDTSLLMKLVGRFFQARDDYQNLESAQYTQQKGFAEDIGEGKLSLPLIHALGSKTPQRGRL
MSILQQRKSTVDLPFHIRKLALDDIKATGGLKYAKKMAMSLQDSVNETLTQYEDKVGAKNWILRLVQKRL
ELEV

>NfSS (UniProt: A1DN30)
MEVWEHSRPIADDTIKKTPSFTTLPIRINKQNDVADAATRRALRDWDYYLHDGLAERALISISELGNLGAFA
YPEVPPERLAIVTYLTDLGILHDDGYEAMDMDQARTEHREFGALFDPHEQLPSRRGTRAAKLKKLVSQILLE
AIRIDRDMGMYMFDMYNKGWLSVAGGEGKVPQFKSVEEYQAYRRDDFGIRAFWPMVEFGMAMRLSDED
KKLIEPVMEPIDKAIIWTNDYWSFDREYHESITNGSRLTNVVEVVRQIENKSIDEAKAAVRQLLVNLEQQYL
ERKRAIYAQNPSIPSHLRKWIEVVGITVAGTHFWASCSPRHHAWRNNSRNGLKPANHVAAPTLITPSNNLNSS
KGSEEQMQDSDNGTRTQMCPANDHEVMQLNAKLSLGKQDGGHAMRAALALLSRAAEQCESLFDGMEHE
RARLLQSGEEKARLSWEGRSKGSQELEHSWYKPAKTALQAPIHYICSMPSKGVRSRMIEAFNYWLEVDETS
LTKIRRLVDLLHNASLILDDIEDHSPKRRGRPATHTIFGHSQAINSANFMFVQAVQVARQFRNPNAVDILLEEL
ENLYLGQSWDLDWKYKLRCPSPSEYLNMVDNKTGGLFRLLLRLMQAERKGTTEVDLDGLTVLFGRFFQIR
DDYMNLRSGLYTEQKGFCEDLDEGKFSYPIVVCVANHADFRDLIDGVFRQRPTAITSGMQPLAPEIKRYVVE
YLNTSGTFQHCREFLMQLESLIESEIDRIEKVTNEANPMLRLLLEKLSVKEN

>PbSS (UniProt: A0A2Z6AQX7)
MDFLSGAFHYSDSVNPSKYSPRPSDYFGTLPFRTSRFEREAADVTADYLRKWQKAVKADNPERKDLVFHGS
TTTLGHFVSWAYPECIPDRVDLCTQICDFGFYWDDVTDSVNVQENAEITQDLALALLSELTLGQRLEPKLEI
NKIVVQMLWGVLDKDRKSGLEMIKFWKGHLDGQAESAHNNMSFEEYTKHRLSEVGARWAVEVGCWSLG
INLSREKKDSVAHFVNKGLLAAALMNDYYSFNKEFDEHQRAGSMDRLQNGLGILMREYGYTETEARSILR
EEIRKGERAIMDGYIAWRESADSSSESHELNRYIVMIILMIGGITFWSSHASRYHRDDLITTAGDRAMIVGKF
QCSMRLLDGYPPPNRWKSATSSNDISGRKRKSWSDSNGVDTHGACYTNGSSNRAKRNGTEAGHKANGHD
SMDIYTAPFLKAPSEVCEAPYEYINSLQGKNMRNKFMDALNHWLCVPAPSMQIIKNIVQMLHNSSLMLDDI
EDESPLRRGQPVAHTFYGISQTINSANFVYVKSVKETSRLKNPICMEIFTDELSNLHTGQSLDLYWRYHGRCP
SINEYIMMVDNKTGGLFRLMLRLMEAESPAASSASLVKLLTLTGRYYQIRDDYLNLTSVEYTSKKGFCEDLD
EGKFSLPLLHLLNHTRHPDRITAPLFNRASGARSLAREVKVHIIQAMDEAGTFEYAQGVLKYLHEEIMRTLD
EVEADLGRNTEARILLLGLGL

>PvPS (UniProt: A0A2Z6AQX6)
MAATKKSTATAAHQIIPSQPTSMSADKFLFSCLQVFVLFFEWARSFLLSTQSRLSAVPAENARLEPSKPISEEE
ESVIEKPASSITPRYSNLVDPSTYNDLGLCSALPLRVHKFAHLADKGALRAQEDWKRLVGPIRNFTGCLSPRF
NGIAVAVPECIPERLEIVTYANEFAFLHDDILDNVGKEEGDHENNEMAAGFGSVLNPADNVKMSASGKSQM
QAKLILELLAINEPQAMVLLKCWEGLVKGESGSQHFNFQRLDEYLPHRVINLGQTFWFGIITFAMGLTISPDE
AEKANNITDPAYATLALANDYFSWEKEYIEFKQNPTSDDMANAIWIIMKEHSVDLEEAKKICQDKIRESCEE
YVRRHRQFEREATGKVSTDLLRYLAALEFSISGNVVWSQYTHRYNFHKPAAKENEDTDDEGAKSDDSKTT
LNDSTDSTVVDVKTPATSGLLSSANDVLMSRTAKSLVGPILDVQLPELPDKVVLSPSQYVKSLPSKKVRHHA
IDALNIWFNVPEAELEVIKEAIDLLHNSSLMLDDIEDDSPLRRGFPSTHVVYGISQTINSANYLYVMALEMTQ
RLNSPACLNVFIDELKRLHIGQSLDLYWTANVQCPSLEEYLKMVDYKTGGLFQMVAKLMALKSPMAGRVP
DLSNMTTLFGRYFQIRDDYQNLMSEEYTNQKGWCEDLDEGKFSLPLIHSLTTKPNVRLQAMLHQRLINGK
MTFEMKKLALDHLAETKSLEYTKEMLGYMYTQLQKEVDFLERQTGSENFLLRLLLKRLQV

>EvQS (UniProt: A0A1Y1C7Q5)
MASEVIVISDHARKEAGTVSVFPVLIHTDYARVIEDVRKVEDQFNSEMKTSIDTKTTADFPELGLAHVTAFTI
PYCRPDRLSIMTRLTEITFFNDDYYDDAGVEKILDYNNHLRECFGGRAEDELTKASAVTKSKQLQASVLVE
MHYIDSELARDMMLTYNRILEVTSLGKNAGLKSLDEYLPFRIGNSGIEVYQDMSCFGMGVKLTKEEKEKLD
PIVIAAHNSTTLINDYHSWPKEVRKYFNEVQATGKADLPVNAVCIFMQTEGLSEQASRQRVREEIIAQQKSH
LAMIQDLVEQEGPLPEKYYMYFKAAQYTASGSEYWAAITSRYPTKTELNQPEVIIVDGELKYESSEIQQTPK
QIATTFNGIPESAKSIIESKVNGTSAHIPDIRHSPADGAQTHHLISDGQFREVINGHANVHTNGKANGTSQGED
LEVYEVTTGNFQRAPEDTVLAPYQYIASLPSKNIRNKFIDALNLWLGVPPLALSSIKRIVEYLHHSSLMLDDI
EDNSTLRRGKPCTHMLYGNAQTINAANYAFVSAFAEVQNLQSPSAITIFIREVQNMHRGQSLDLSWKYHTH
CPTVDEYMMMVDNKTGAMFRLCVQLMQAESSVPCQKITQSDFITQLGRYFQIRDDYQNLVSSEYTTQKGF
CEDLDEGKISLPLIYTIMDSSPEASVVKGIFHHRLREGGLPLHLKEYILSQMEEKGALSATHSLLQKMQKELI
EGLHRVEETFGSKNALVELMLRRLWV

>CgDS (UniProt: P9WEV7)
MASTMMNYQDCGPMRYKSSVPVPASLYENTAYPSKFRPRISKHVDVADKACWEACDDFENATGLKLKADS
VGCINPIGGNVNALWFPEAIPERLHIISYLSELLFRHDDLTDDAVTPEQFDEVHGPLARFLGSESKQSDHTTK
HNAMNTMQARVAIEALEQNEQLGKLVIEKWKGIVSVRGQDAFMEHKTLDSYMHVRHYDAGAYSVWSQIL
FCCDISLTDEELTGLEPLTWLAFTQMILWHDYCSWDKEAATYLEREEGGSNMSAVQVYMAMYGLDQYAAK
EFLLSEITRIEDEYCERKASYMIEFPPAPHITHYIGLIEMCMAGNTLWHLSSRRYNPAAPLPRREDIGKVNGGP
LDASEVSKPVECSESDLGILTPVSSRLSTKRLRPFWNNQRTEYTTMTPAETSSDDKKKKAKASHETREDLLT
VSPCAWPAEPDEKDILAAYLYTAARPASGARDKLMDALDKWYRVPPDALATIRTIIRIMHNASLMLDDVQD
NSPVRRGSPSAHVIFGTAQTTNSASYLMIKCVDLARRLGDDSLSCLLSELSQLHLGQSHDLAWTFHCKAPSI
PEYYSHLEQKTGGLFRMASRMMRASATQNKHLDACKLMSLLGRLYQLRDDYQDITSESLSTYDDLDEGSF
TLPLIHALHREEEQGEVQLHSILQSARAARSASASSNNDGKLSVETKLLIREMLEEAGSLEHTRVVIRGLYDE
TRAVLTAMENEAGSGGKNWMLHLITFQLKV

>AcSS (UniProt: A0A0U5GLI1)
MGTMGSEKSPLPYTRSDPVPIDAETECKYISRFQPRISKHAAVSQDACIECHIDLFGMDKIGKVAGGMNAHT
ADFTALCAPEALPERIALCSYFIEYAFVHDDVSVDAVQGAETCREGTKTLSKALGPERYEDLQAIMDGGLKR
KQSRAKIWSALREIDQDYLGRCQSTFKQWYETGQQMRDQTFASLENYLAVRALDCGANWVVRMMGWAS
GVELTQEEEIETGPVTYLAFVVLAVTNDIWSWEKEKTVTRDSGESLPLINAVQMVIQMQQVDEDTAKHRVL
DVIRQNEKQYCFLRDEHLKRPNTSHSVRKWFQILELSMAGNALWSIHAPRYHLNVQNPYISPSVVPSVFKEL
QIMQSRSKGGKAEKTQTQKTTEMKNPQTVLERLDDLVLWRPYEYITSLPSKGIRQHLIDALQTWFNVPRSS
LAAISGVASLLHEASLMLDDIQDGSPLRRGQPAVHEMFGVGQTINSACYCINNALRLVQEISPSAALIFSEQM
GHLYIGQAHDISWAGQRAIPTEEEYFEMVDGKTGALFVLLFRLMQSEASENRNLDMCPFMNKLGRCFQIRD
DYQNLASQEYTSQRGFCQDLDEGKPSFPFIRACHELQDSTALTEWFKMPRNGAGASVEVKRYILSQIRGSGS
FEYTKELLSHLLYDLEDMVRDMESVTGQKNWILRNILVQMRVKEERAVQKKETTVGEVMRVWGQYQETA
WTSSLS

>FoFS (UniProt: P9WER6)
MDQLSYQSRLIPPEEAQQTGCFTSLPIRIHPRNDIADAATAKFIADWAKHVGDGREKRTHFCPSRVGNWNSL
LYPEGLPERLGSVSYLLDLGLIHDDVNEELSVQDAMAAHERLRPALDPQDNRKWDPESPQMKFKMLLSEC
VIECIKTDRELGTAMLKSFRVLWLDIAENATSDAPQTMDDYWDVRMTNGGMSVFWPMVLYATNLRLSEEQ
HTLVQPIIAAAEEALCWANDYFSYEREVWELETGKAKRIVNIVEMVSRTKGLSSAEAKAEVKRMILGAEAK
YCRLRDDLLSSNPEMSMDLKRWIEYIGLSISGNHYWLSACSRQNTWKTNCSIDGKINGLTNGSVNDTNNRS
VDGVVNGTVDTGIEEPSTGNKDTSLKALKLLFNSTPNESHPVCRYPNDKLSDYAMVAPMTHISSLPSKGTRS
ELISALNVWLKVPPVVLGHISSAIDMLHNASLILDDIQDNSPLRRGVPAAHVVFGTAQSINSATFMFVKATEA
VRSTLSPAALEALLRGLQTLFMGQSWDLYWKHNLQCPAEGDYIRMVDHKTGGMFVMLVQLMAAESPYY
GASVIEDLERLMRLLGRFYQIRDDYMNFSAYSAQKGFAEDLDEGKFSFPVVCGFERDPELRGQILAIFRQRP
TSGAGEATQLSRKVKEHLIRCIAASGGFDETLKCLRSLENELDTEIAELEKKLGQVNPLLRLCLATLSMEGCE
KICW

>TpcA (UniProt: M2U578)
MEQLSYQSKLICSDESRHTGCFTTLPIRIHPRDDLADAASRRFVQDWAREMRDGREQSTHFSFSPVGNWSSL
IYPEAIPERLGVLAYLSDLGLIHDDGGEGLSIEDAQAEHGELCAALDPSDISSAAPGSRAMKTKKLVSQCML
ECISLDRELGLKMLAAFRDVWLAISERNSDKEAQTMEEYLKYRSDNGGMLVFWPMLQFSLGMSISEAEEA
LVQPIIDAATEGLLLANDYFSWEREYRELQSGQSKRIVSAVDLFIRTKGLSIDDAKEEVKRKIIAAERDFCQRR
DDLYTNHPNIPLKLKRWIDCAGLAVSGNHYWCSACPRQNAWKDMSSQSLNGAKRKTSHGATIGMHEAPFK
KRKDSSFFGSQPSDDEPSLSEVSSYPFYKPSGLALEAPSKYVSDMPSKGVRSTLIEALNTWLHVPSERLDSIM
SVINTLHNASLILDDLEDNSPLRRGYPATHILFGHSQSINTANFMFVRAVQEVAQNLSPNALVALLEELKGLY
LGQSWDLYWKHNLACPSEAEYVNMIDHKTGGMFRMLLRIMQAESDVTPQPDFHRLTLLFGRFFQIRDDYM
NFQDYTAQKGLCEDLDEGKFSYPVVYCLENHPEYRGYFLSMFRQRPTIATVNACPLSGESKQYLTACLKKS
GAFNKTIACLTDMERDLEFEINRLEQQTGETNPMLRLCLAKLSVKGIGRIGEVSPSTSK

>FgMS (GenBank: KY462789)
MDFTYRYSFEPTDYDTDGLCDGVPVRMHKGADLDEVAIFKAQYDWEKHVGPKLPFRGALGPRHNFICLTL
PECLPERLEIVSYANEFAFLHDDITDVESAETVAAENDEFLDALQQGVREGDIQSRESGKRHLQAWIFKSMV
AIDRDRAVAAMNAWATFINTGAGCAHDTNFKSLDEYLHYRATDVGYMFWHALIIFGCAITIPEHEIELCHQL
ALPAIMSVTLTNDIWSYGKEAEAAEKSGKPGDFVNALVVLMREHNCSIEEAERLCRARNKIEVAKCLQVTK
ETRERKDVSQDLKDYLYHMLFGVSGNAIWSTQCRRYDMTAPYNERQQARLKQTKGELTSTYDPVQAAKE
AMMESTRPEIHRLPTPDSPRKESFAVRPLVNGSGQYNGNNHINGVSNEVDVRPSIERHASTKRATSADDIDW
TAHKKVDSGADHKKTLSDIMLQELPPMEDDVVMEPYRYLCSLPSKGVRNKTIDALNFWLKVPIENANTIK
AITESLHGSSLMLDDIEDHSQLRRGKPSAHAVFGEAQTINSATFQYIQSVSLISQLRSPKALNIFVDEIRQLFIG
QAYELQWTSNMICPPLEEYLRMVDGKTGGLFRLLTRLMAAESTTEVDVDFSRLCQLFGRYFQIRDDYANLK
LADYTEQKGFCEDLDEGKFSLPLIIAFNENNKAPKAVAQLRGLMMQRCVNGGLTFEQKVLALNLIEEAGGIS
GTEKVLHSLYGEMEAELERLAGVFGAENHQLELILEMLRID

>AcOS (UniProt: A0A1V1FVQ6)
MEYKYSTIVDKSKWDPEGLTEGIPLRRHEAGDLEEVGSFRVQEDWRRLVGPLENPYRGSLGPEISFITYTVPE
CLPERLEAISYSLDYGFMHDDEIDLNIASAELSDVGGALKQGGATGKIDEGKSSSGKRKMAAQLLREMMAL
DPERAMALAKSWAQGVQHSARRVEEKDWKSLDEYIPFRCMDLGYMHWHGLVTFGCAITVPEEEEEERRR
LLEPAVIACMMTNDLFSYEKEKNDNNPQNAVTVIMKINKCGEEEAKEVCKKRIRVECAKYAQIVKETLART
DISLDLKKYIEIMQYTVSGNWAWSTQCPRYHFPGRWNELQKLRAEHGIAKYPARYSLKERTNGVNGVNGV
NGINGTNGINGTNGVNGKRNRDEDGDENDARINGNGFKKPALTSQGKDSFVLDDVVALSLNLHLPDLGDG
VVLQPYRYLTSLPSKGFRDMAIDALNTWLRVPSTSTSTIKDLIKKLHSASLMLDDIEDNSPLRRAKPSTHIIYG
NAQTINSATYQYTEATSLAANLSNPLSLRIFLDEIQQLYIGQSYDLYWTHNALCPSITEYLRMVDQKTGGLFR
MLTRLMVAESPGSNKILDRALFPLSHLIGRFFQIRDDYQNLSSAEYSRQKGFVEDLDEGKYSFTLIHCIQTVE
ANEALASEMMALRAFLIKRRVDGGLSNEAKMEVLGIMKKTKSLEYTLGVLRALQEELEREVGRLEGKFGE
ENLPLRLMVDMLKV

>PaPS (UniProt: P9WEV6)
MEYRYSYVIDPSSYDNQGLCNGIPLRVHRNADIEEYATISLRNDWRKHVGPLPLTSYGGNLGPKYNFTAVTL
PECRPDRLEIVSYIMEFAFLHDDLVDTAQVDEALALNDTWRDGITEGLDTTSAKGKKSGEGLILRNILKEVT
AIDPVRAAELMKFWKRDLDVSRDRKHFRDFDDYMEYRIVDCASYFLIALSTFAMALTIPAEDKDEVFTLLT
RPVWAAAALTNDVQSWEKEDKLFQKDNATDMTNGVWMLMKQYSIGVEEAKRRILGKAREHVAEFVKTL
SQIHNRLDLSLDSRLFVEAMQYMISGNLMWGISTPRYHSDQSLDEMMVARMKYGWPNHREVTKLTSDLEN
RGTKRTHQDDTEGVQSVKRFNGASTKNGINGTNGINGLNGINGSNGVKIKRHKNKEYSGALTKDSDLVLN
MDLNGLSSAIICAPADYIGSLPSKGIRDNVADALSIWLDVPAKELNQIKRAINLLHNASLMLDDVQDGSVLR
RAQPTTHTVFGPAQTINSAGHQIIQAMNEIRKLGSDDCLDIFSEELEKLYVGQSHDLYWVYNDSCSPTIEDYF
KMVDYKTGGLFNMLARLMTAKSSSSSSPDLTALVGLLGRYFQIRDDYMNLTSADYTVEKGFCEDLDEGKFS
ITLLHALSAAPEPEALLLRNLMSGRRNDGKLSVVQKNLALSIIEGARSLEYTAAVLQKLYKAIVRELESTERQ
FGENKPFRFLLSLLKV

>PrDS (UniProt: W6QAE7)
MGETIADVYAESIDPEIYANNPAYSSLFTPYIHKQTIIADHVSVQCHIDLNGIDAVGSKFGNLNAHAGNFTSLC
APNCLPERLALVAYTVEYAFLHDDETDNAADQEALLLENKMLHQAINQSSMTSVSNRVSAKAQRKSEVQA
KIAAEYLRLDPVFGEFFLKAWQTFTASVQDVRSLEFPSLDDYLEFRIVDAAADWTLYNFRWGSGITLTPEEE
KIADPMSYVAYAELCLVNDLFSWDKEYDAHVKSNGEVPLVNAVHIVAVTQGLTHCAAKAVVQAEIRAHEER
FCYLKEQYKATASPSDSILSWLKLLEHSMAGNWVWSLCVPRYFKVERNPYKDHLEKFGSEAVRVLTPEEHL
RDSKQEINGTKEIELQEPKSNNTAESDVLAKYTSGYPTIDEPVLNPYTYINSLPSKNVRQTMIAALNSWYKV
PVKSLLIIEGAVNFLHNSSLLLDDIQDGSVLRRGRPVAHQIFGVGQTINTATYLMNEALYLVQMLSPSAVLVY
TDEMRNLQLGQGRDLHWSYHTHVPTPAQYISMVDGKTGGLFRLISRLMRSEATVNRDLDISQFATLLGRHF
QIRDDYQNLQSDDYTKNKGFCDDLDEGKLSFPIILSMQSPGFSNTALSSVFKGSQKGETLSPEMKQYILEEIT
ARGAFSQTKAVLRKLHIELLRLLMETEQKAGGIENWALRLLIMKLDLGDEKKKEAHKSDSAWKVNQRRA
WKGSQKNGRPIDKACFLRAMEEASQK

>PaFS (UniProt: A2PZA5)
MEFKYSEVVEPSTYYTEGLCEGIDVRKSKFTTLEDRGAIRAHEDWNKHIGPCGEYRGTLGPRFSFISVAVPE
CIPERLEVISYANEFAFLHDDVTDHVGHDTGEVENDEMMTVFLEAAHTGAIDTSNKVDIRRAGKKRIQSQL
FLEMLAIDPECAKTTMKSWARFVEVGSSRQHETRFVELAKYIPYRIMDVGEMFWFGLVTFGLGLHIPDHEL
ELCRELMANAWIAVGLQNDIWSWPKERDAATLHGKDHVVNAIWVLMQEHQTDVDGAMQICRKLIVEYVA
KYLEVIEATKNDESISLDLRKYLDAMLYSISGNVVWSLECPRYNPDVSFNKTQLEWMRQGLPSLESCPVLA
RSPEIDSDESAVSPTADESDSTEDSLGSGSRQDSSLSTGLSLSPVHSNEGKDLQRVDTDHIFFEKAVLEAPYDY
IASMPSKGVRDQFIDALNDWLRVPDVKVGKIKDAVRVLHNSSLLLDDFQDNSPLRRGKPSTHNIFGSAQTV
NTATYSIIKAIGQIMEFSAGESVQEVMNSIMILFQGQAMDLFWTYNGHVPSEEEYYRMIDQKTGQLFSIATSL
LLNAADNEIPRTKIQSCLHRLTRLLGRCFQIRDDYQNLVSADYTKQKGFCEDLDEGKWSLALIHMIHKQRSH
MALLNVLSTGRKHGGMTLEQKQFVLDIIEEEKSLDYTRSVMMDLHVQLRAEIGRIEILLDSPNPAMRLLLEL
LRV

>EvAS (UniProt: A0A169T193)
MEFKYSTLIDPEMYETEGLCDGIPVRYHNNPELEEIDCLRCHEHWRENVGPLGVYKGGLADQWNGISIAIPE
ALPDRLGVVSYASEFAFVHDDVIDIAQHGNEQNDDLRVGFEQMIDAGAIKYSTSGKRALQSYIAKRMLSIDR
ERAIISLRAWLEFIEKTGRQEERRFNNEKEFLKYRIYDVGMLFWYGLLTFAQKITIPENELTTCHELAIPAYRH
MALLNDLVSWEKERASSIALGKDYCINFIFVAMEESGISEDEAKERCREEIKLATVDYLRVFDEAKDRIDLSH
DTMLYLESLLYSMSGNVVWGLQSPRYYTDAKFSQRQLDWIKNGLPLEVRLEDRVFGLSPSEDRVTHQAVIE
NGLPESGLGKNGNSSNGVDVNKALLSAVLHEHLKGHAVFKMSDHEVKVKASNGRSLDTKVLQAPYEYITG
LPSKRLREQAIDAMNVWFRVPAEKLDLIKSITTILHNASLMLDDVEDGSELRRGNPSTHTIFGLSQTINSANY
QLVRALERVQKLEDSESLLVFTEELRNLYIGQSMDLYWTGNLICPTMNEYFHMVECKTGGLFRLFTRLMSL
HSTSAVKVDPTTLSTRLGIYFQTRDDYKNLVSTEYTKQKGYCEDLEEGKFSLPLIHLIQAMPDNHVLRNILTQ
WRVTRKVTLAQKQVVLGLMEKSGSLKFTRETLASLYSGLEKSFTELEEKFGTENFQLKLILQFLRTE

>TndC (UniProt: A0A8K1AY78)
MEYRYSTVVDPSTYETHGLCDGIPLRCHESPELEEIETLRCQEDWRRWVGPLGFYKGGLGPRWNFMAITVP
ECLPERLGVLGYANELAFLHDDVTDVADYGDLHNDDLKAAFEQAASTGHIEGSASGKRAIQAHIANEMMS
IHKQHAITTLEAWAKFAELGSGRQHTTHFKTEDEYIKYRMIDIGTMFWYGMVTFGMGISIPEHELEMCHRL
ADTAYLNLGLTNDLYSWQKEYETAVAMDRDYVANIIGVIMEERNISEAEAKEVCREKIKKTIVDFRKIVDDT
KARDDVSLDTKRYLEGLLYSLSGNLVWSIDCPRYHPWSSYNERQLDWMKNGIPKSPPKVNGNATANGNGV
HHAPKESLANGTLNGHDRIHAPAVNGNGASHTSSIKGSTGGNGVTHSPVSNGSAVVNSALSMEIDTDLVNV
FARKEYKSINGFKMHEGDNHPSNGQTKLNGNVTSWKVPGDVQTKVIQAPYDYISSLPSKGVRDHAIDALN
VWCRVPAAKLDLIKLITNMLHNTSLMLDDLEDGSHLRRGRSSTHTIFGAGQTVNAANYHVIRALEEVQKFG
DAESIVIFIEELKSLYVGQSLDLYWTNNAICPSVDEYFQMVENKTGGLFRLFGRLMSLHSSHPVKADLTGFL
NQFGRYFQTRDDYQNLTSPEYTKQKGFCEDLDEGKFSLPLIHLMHSAPSNLVVRNIWTQRLVNNKASPAHK
QTILELMKENGSLQFTMDALDVLHAKVEKSISDLEARFGVENFQLRLILEMLRKA

>EvSS (UniProt: A0A0P0ZEM1)
MEYKFSTVVDPGTYETHGLCEGYEVRYHKNAELEDIGCLRCQEHWRQSVGPLGAFKGTLGNPFNLLSLVIP
ECLPDRLSIVGFANELAFIHDDVTDIVQYGDAHNNDFKEAFNSMATTGSMENAASGKRALQAYIAREMVRI
DKERAIPTIKAWAKFVDYGGRQETTRFTSEKEYTEYRIQDIGLWFWYGLLSFAMALDVPEHEREMCHEVCR
TAYVQIMLVHDLASWEKEKLNAAALGKDVITNIIFVLMEEHGISEEEAKERCRETAKTLAADYLKIVEEYKA
RDDISLDSRKYIESWLYTISGNTVWSFICPRYNSSGSFSDHQLELMKNGVPKDPASGSTNGTSNGTSNGTSH
VAVNGNGHVTNDDLSANGIKTDGELLSAITMEHLKNRNSFKLGDHDQEVKSLHGHGQALDPRVLQAPYEY
ITALPSKGLREQAIDALNVWFRVPTAKLEIIKSITTILHNASLMLDDVEDGSELRRGKPATHNIFGLGQTINSA
NYQLVRALQELQKLGDARSLLVFTEELHNLYVGQSMDLYWTSNLVCPSMHEYFQMIEHKTGGLFRLFGRL
MAVHSTNPVQVDLTDFTNHLGRYFQTRDDYQNLVSAEYTKQKGFCEDFEEGKFSLPMIHLMQTMPDNLVL
RNVWTQRRVNGTATHGQKQTILNLMKEAGTLKFTQDSLGVLYSDVEKSVAELESKFGIENFQLRLIMELLK
TG

### Table S1. Cryo-EM data collection, reconstruction, and refinement statistics

| | EvVS Hexameric Core | EvVS with Six Cyclase Domains | EvVS with Five Cyclase Domains | EvVS with One Cyclase Domain (Map 1) | EvVS with One Cyclase Domain (Map 2) | EvVS with One Cyclase Domain (Map 3) |
|---|---|---|---|---|---|---|
| **Data Collection** | | | | | | |
| Magnification | 81,000 | 81,000 | 81,000 | 81,000 | 81,000 | 81,000 |
| Voltage (kV) | 300 | 300 | 300 | 300 | 300 | 300 |
| Exposure ($e^-/Å^2$) | 43 | 43 | 43 | 43 | 43 | 43 |
| Defocus range (μM) | -1.0 to -3.0 | -1.0 to -3.0 | -1.0 to -3.0 | -1.0 to -3.0 | -1.0 to -3.0 | -1.0 to -3.0 |
| Pixel size (Å/pix) | 0.54 | 0.54 | 0.54 | 0.54 | 0.54 | 0.54 |
| Symmetry imposed | *C*3 | *C*3 | *C*1 | *C*1 | *C*1 | *C*1 |
| Initial particles (no.) | 421,245 | 421,245 | 421,245 | 421,245 | 421,245 | 421,245 |
| Final particles (no.) | 491,079 | 60,587 | 51,215 | 298,050 | 294,158 | 289,027 |
| Map resolution (FSC=0.143) (Å) | 2.77 | 3.18 | 3.59 | 3.00 | 2.98 | 3.08 |
| **Model** | | | | | | |
| Chains | 6 | 6 | 6 | 7 | 7 | 7 |
| Atoms | 13,074 | 28,392 | 25,560 | 15,523 | 15,439 | 15,415 |
| Residues | 1,646 | 3,522 | 3,174 | 1,939 | 1,928 | 1,926 |
| Water | 0 | 0 | 0 | 0 | 0 | 0 |
| Ligands | 0 | 0 | 0 | 4 | 4 | 4 |
| **Root-mean-squared deviations[a]** | | | | | | |
| Bond lengths (Å) | 0.004 (0) | 0.005 (0) | 0.004 (0) | 0.005 (0) | 0.004 (0) | 0.005 (0) |
| Bond angles (°) | 0.7 (3) | 1.0 (3) | 0.8 (19) | 1.0 (1) | 1.0 (3) | 1.0 (0) |
| **Validation** | | | | | | |
| MolProbity score | 2.21 | 2.93 | 2.94 | 2.11 | 2.13 | 2.37 |
| Clash score | 8.51 | 19.43 | 21.12 | 10.12 | 9.95 | 9.41 |
| Poor rotamers (%) | 4.7 | 8.03 | 6.74 | 2.32 | 2.33 | 5.25 |
| **Ramachandran plot (%, MolProbity)** | | | | | | |
| Favored | 96.36 | 92.80 | 91.92 | 95.65 | 95.25 | 95.41 |
| Allowed | 2.96 | 6.33 | 7.27 | 3.72 | 3.90 | 3.70 |
| Outliers | 0.68 | 0.87 | 0.81 | 0.63 | 0.84 | 0.90 |

| | | | | | | |
|---|---|---|---|---|---|---|
| **Peptide plane (%)** | | | | | | |
| Cis proline/ general | 0.0/0.0 | 0.0/0.0 | 0.0/0.0 | 0.0/0.0 | 0.0/0.0 | 0.0/0.0 |
| Twisted proline/ general | 0.0/0.0 | 0.0/0.0 | 0.0/0.0 | 0.0/0.0 | 0.0/0.0 | 0.0/0.0 |
| **B-factors (Å$^2$)** | | | | | | |
| (min/max /mean) | 30/138 /57 | 13/218 /104 | 30/206 /106 | 22/168 /60 | 25/131 /62 | 26/141 /60 |
| **Model vs. Data** | | | | | | |
| CC (mask) | 0.80 | 0.77 | 0.73 | 0.80 | 0.80 | 0.80 |
| CC (box) | 0.79 | 0.76 | 0.74 | 0.80 | 0.79 | 0.80 |
| CC (peaks) | 0.79 | 0.76 | 0.72 | 0.79 | 0.79 | 0.79 |
| CC (volume) | 0.80 | 0.78 | 0.74 | 0.80 | 0.80 | 0.80 |
| Mean CC for ligands | - | - | - | 0.66 | 0.68 | 0.71 |
| **Accession codes** | | | | | | |
| PDB | 9E2H | 9E2I | 9E2J | 9E2K | 9E2L | 9E2M |
| EMDB | EMD-47452 | EMD-47453 | EMD-47454 | EMD-47455 | EMD-47456 | EMD-47457 |

[a]Numbers in parentheses indicate the number of outliers ≥4σ in each refined structure.

**Table S2. Attributes of bifunctional terpene synthases used in this study for multiple sequence alignments and AlphaFold predictions**

| Enzyme | ipTM Score | pTM Score | Source Organism | Product | Cx | Linker length | Ref. |
|---|---|---|---|---|---|---|---|
| EvVS | 0.69 ± 0.01 | 0.70 ± 0.00 | *Emericella variecolor* | Variediene | 20 | 66 | 1 |
| MpMS | 0.68 ± 0.01 | 0.71 ± 0.01 | *Macrophomina phaseolina* | Macrophomene | 30 | 61 | 2 |
| CoSS | 0.68 ± 0.01 | 0.70 ± 0.00 | *Colletotrichum orbiculare* | Sesterorbiculene | 25 | 108 | 3 |
| CsSS | 0.67 ± 0.01 | 0.69 ± 0.01 | *Cytospora schulzeri* | Schultriene | 25 | 92 | 4 |
| ZbSS | 0.67 ± 0.01 | 0.69 ± 0.01 | *Zymoseptoria brevis* | Sesterevisene | 25 | 86 | 3 |
| PfVS | 0.66 ± 0.02 | 0.67 ± 0.02 | *Pestalotiopsis fici* | Variculatriene A | 25 | 43 | 3 |
| NfSS | 0.63 ± 0.01 | 0.65 ± 0.01 | *Neosartorya fischeri* | Sesterfisherol | 25 | 128 | 5 |
| PbSS | 0.63 ± 0.03 | 0.65 ± 0.03 | *Penicillium brasilianum* | Sesterbrasiliatriene | 25 | 107 | 6 |
| PvPS | 0.62 ± 0.02 | 0.63 ± 0.01 | *Penicillium verruculosum* | Preasperterpenoid A | 25 | 79 | 6 |
| EvQS | 0.61 ± 0.06 | 0.63 ± 0.04 | *Emericella variecolor* | Quiannulatene | 25 | 123 | 7 |
| CgDS | 0.61 ± 0.04 | 0.63 ± 0.04 | *Colletotrichum gloeosporioides* | Dolastadiene | 20 | 111 | 8 |
| AcSS | 0.59 ± 0.01 | 0.62 ± 0.01 | *Aspergillus calidoustus* | Spiroviolene | 20 | 60 | 3 |
| FoFS | 0.59 ± 0.04 | 0.60 ± 0.04 | *Fusarium oxysporum* | Fusoxypene | 25 | 71 | 9 |
| TpcA | 0.58 ± 0.05 | 0.60 ± 0.05 | *Cochliobolus heterostrophus* | Preterpestacin I | 25 | 68 | 10 |
| FgMS | 0.51 ± 0.08 | 0.52 ± 0.07 | *Fusarium graminearum* | Mangicdiene | 25 | 119 | 11 |
| AcOS | 0.47 ± 0.02 | 0.49 ± 0.02 | *Emericella variecolor* | Ophiobolin F | 25 | 112 | 12 |
| PaPS | 0.47 ± 0.02 | 0.49 ± 0.02 | *Phomopsis amygdali* | Phomopsene | 20 | 114 | 13 |
| PrDS | 0.46 ± 0.03 | 0.48 ± 0.04 | *Penicillium roqueforti* | Deoxyconidiogenol | 20 | 55 | 14 |
| PaFS | 0.43 ± 0.01 | 0.45 ± 0.01 | *Phomopsis amygdali* | Fusicoccadiene | 20 | 69 | 15 |
| EvAS | 0.43 ± 0.01 | 0.46 ± 0.01 | *Emericella variecolor* | Astellifadiene | 25 | 107 | 16 |
| TndC | 0.43 ± 0.01 | 0.45 ± 0.01 | *Aspergillus flavipes* | Talarodiene | 20 | 151 | 17 |
| EvSS | 0.43 ± 0.01 | 0.45 ± 0.01 | *Emericella variecolor* | Stellatatriene | 25 | 103 | 18 |

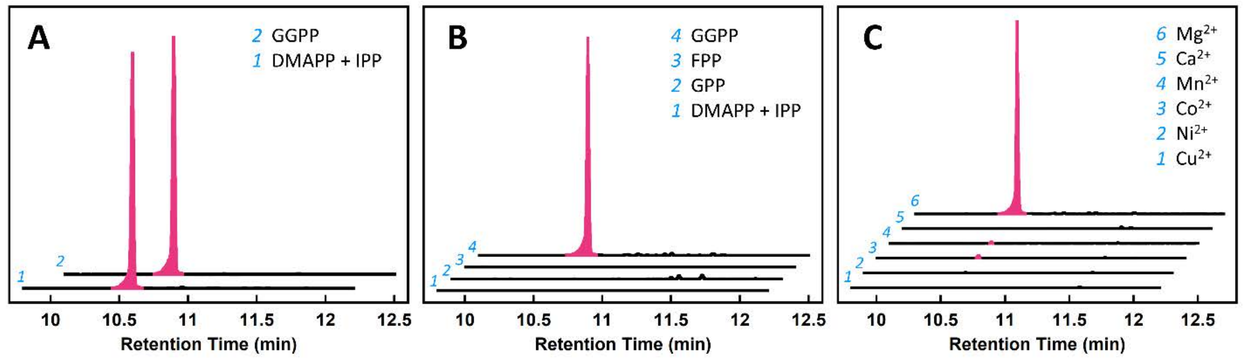


**Figure S1. GC-MS characterization of the reactions catalyzed by EvVS.** (**A**) The same single product, variediene, is generated when either GGPP or DMAPP and IPP are provided to the full-length enzyme. Therefore, both the prenyltransferase and cyclase domains are catalytically active. (**B**) The individual cyclase domain construct, $EvVS_{CY}$, forms no products from DMAPP and IPP, geranyl diphosphate (GPP, $C_{10}$), or farnesyl diphosphate (FPP, $C_{15}$). (**C**) $EvVS_{CY}$ exhibits maximal catalytic activity with $Mg^{2+}$; trace product is observed when $Mn^{2+}$, $Co^{2+}$, and $Ni^{2+}$ are utilized. No product is observed when $Cu^{2+}$ or $Ca^{2+}$ are utilized. These results are similar to those reported upon the initial discovery and characterization of full-length EvVS.[1]

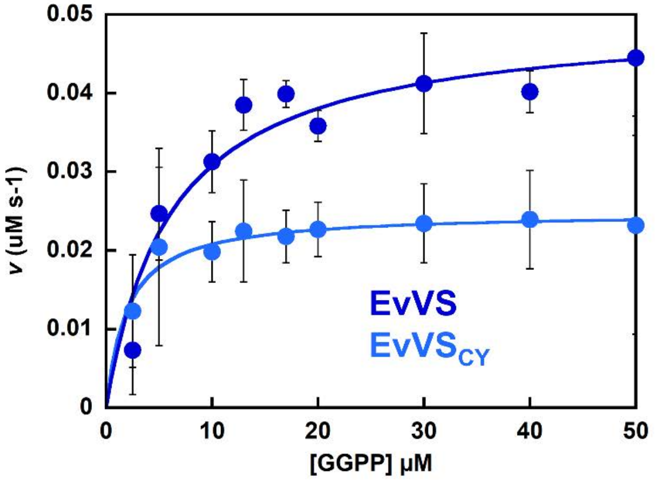


| Catalytic Parameter | EvVS | $EvVS_{CY}$ |
|---|---|---|
| $V_{max}$ (µM $s^{-1}$) | 0.050 | 0.025 |
| $k_{cat}$ ($s^{-1}$) | 0.1 | 0.05 |
| $K_M$ (µM) | 6.15 | 1.94 |
| $k_{cat}/K_M$ ($M^{-1}s^{-1}$) | 16000 | 26000 |
| $R^2$ | 0.89 | 0.89 |

**Figure S2. Steady-state kinetics of GGPP cyclization by EvVS and $EvVS_{CY}$.** The EnzChek™ pyrophosphate detection kit was used to measure the steady-state kinetic parameters of EvVS (dark blue) and $EvVS_{CY}$ (light blue). Parameters derived from fitting to the Michaelis-Menten equation are listed at right. Each data point represents the average of two independent measurements.

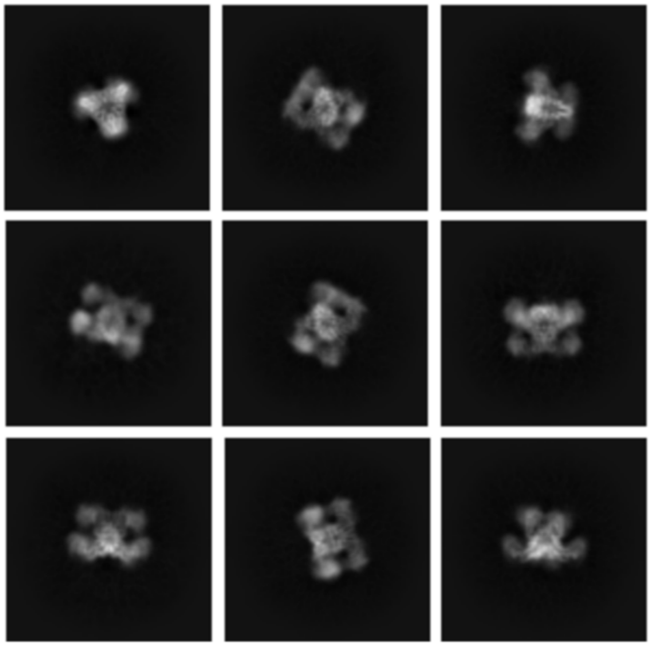

**Figure S3. 2D classes of EvVS determined by cryo-EM.** The top-left class shows the best top-down view, while other classes reveal side-on orientations with clear densities corresponding to cyclase domains above and below the prenyltransferase hexamer.

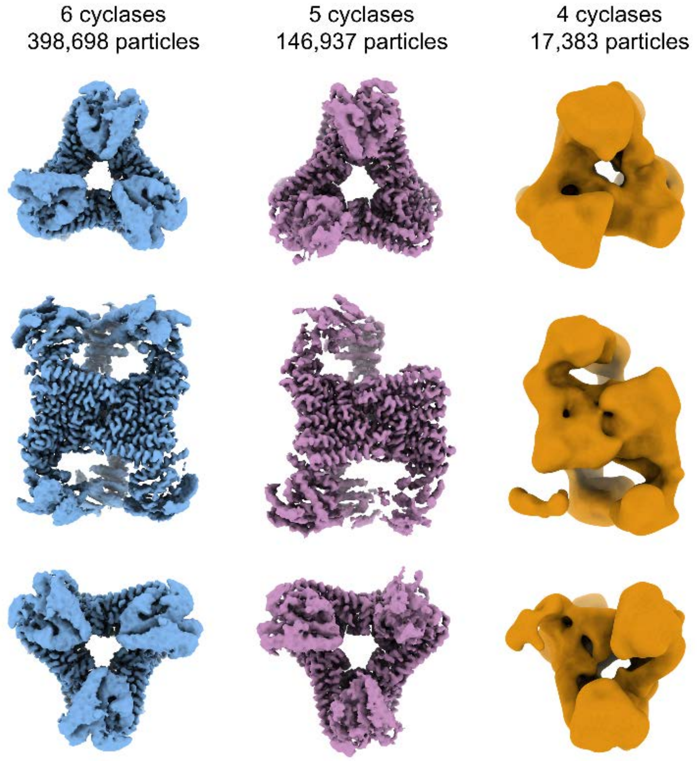


**Figure S4. Initial sorting of EvVS particles.** Top, side, and bottom views are shown. The majority of particles (63%) reveal densities corresponding to all six cyclase domains bound to the hexameric core (blue), although some particles (23%) are missing density corresponding to one cyclase domain (purple) and others still (3%) are missing densities corresponding to two cyclase domains (orange). All high-quality particles contain either five or six cyclase domains associated with the central prenyltransferase hexamer. Particles with densities for only four ordered cyclase domains (orange) reveal weak, uninterpretable density in a fifth site, suggestive of conformational heterogeneity. These particles have poor quality overall and align to only 9.9 Å resolution.

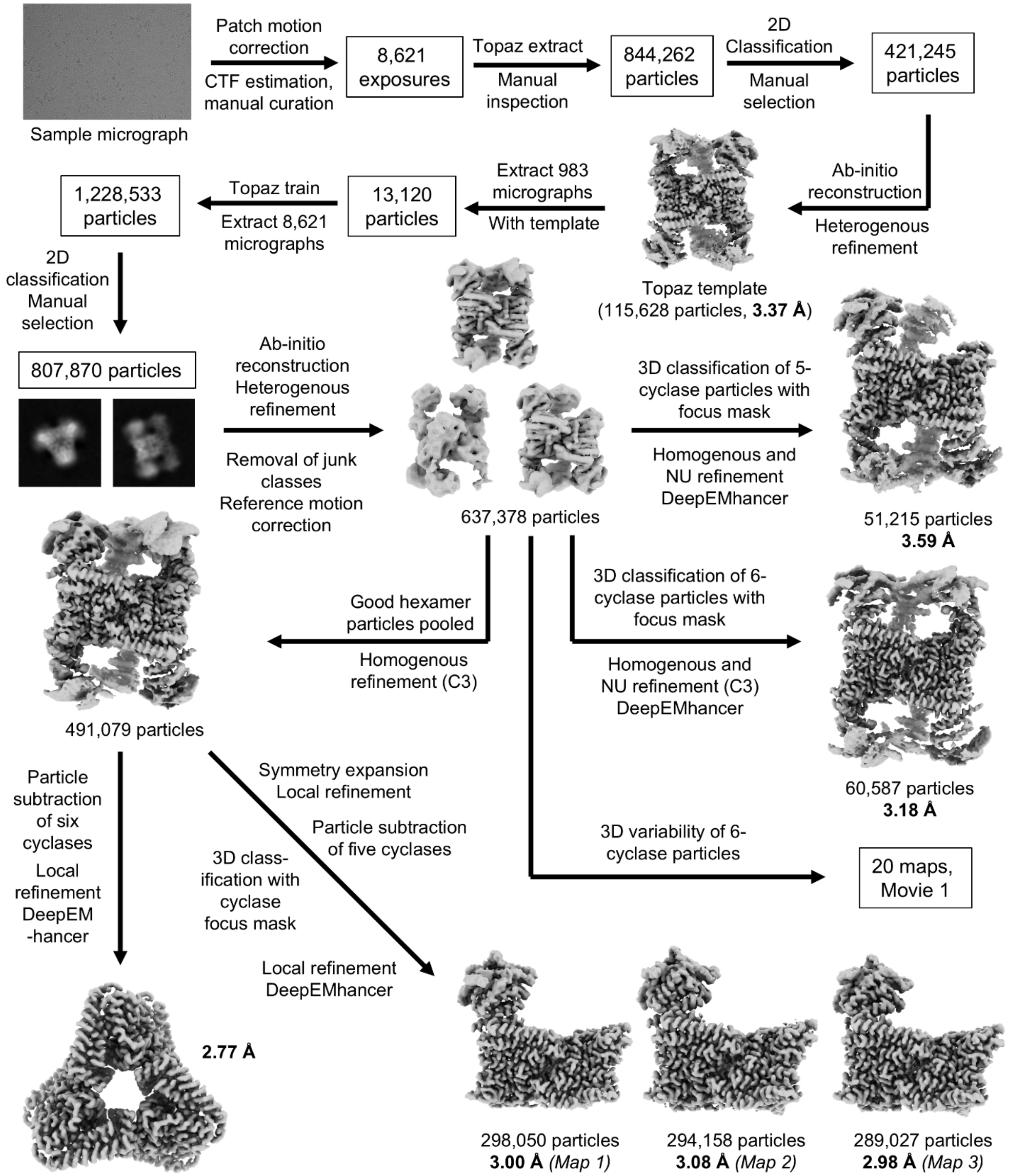


**Figure S5. Cryo-EM workflow.** A total of 8,621 exposures (sample micrograph shown) were taken from a single grid. All data processing and analysis steps were performed using cryoSPARC. Topaz was used to improve particle picking. Map sharpening was achieved using DeepEMhancer.

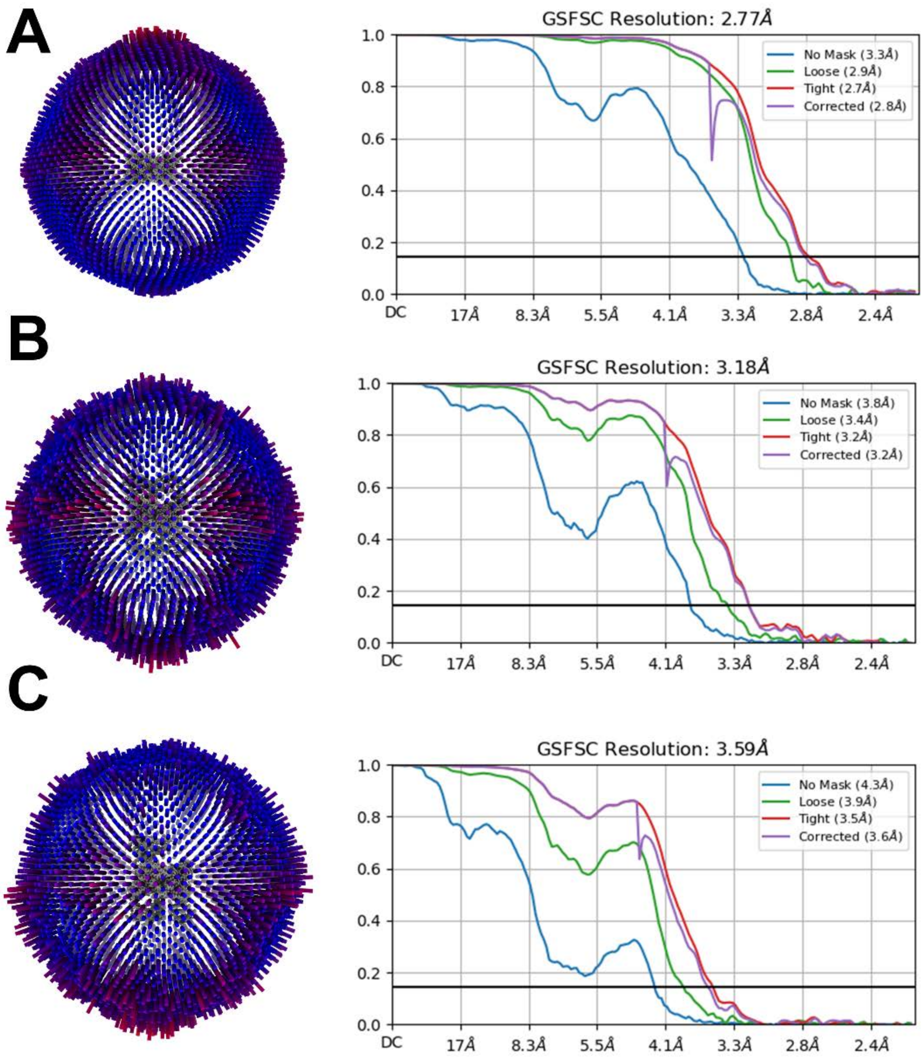


**Figure S6. GSFSC and data coverage plots for Cryo-EM structures.** (A) Angular distribution plot and Gold-Standard Fourier Shell Correlation (GSFSC) plot for the structure of the hexameric prenyltransferase core of EvVS. The resolution estimate at GSFSC = 0.143 is 2.77 Å. (B) Angular distribution and GSFSC plot for the map of EvVS with six cyclases. The resolution estimate at GSFSC = 0.143 is 3.18 Å. (C) Angular distribution and GSFSC plot for the map of EvVS with five cyclases. The resolution estimate at GSFSC = 0.143 is 3.59 Å.

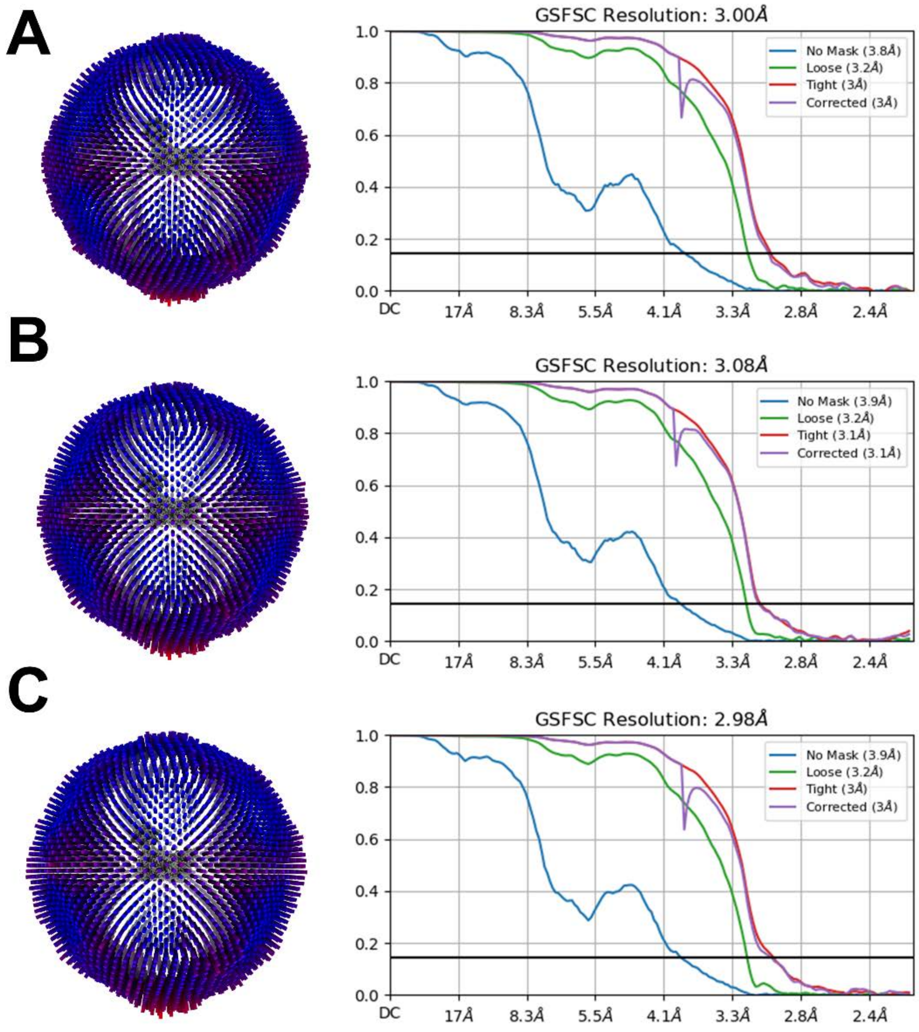


**Figure S7. GSFSC and data coverage plots for Cryo-EM structures.** (A) Angular distribution plot and Gold-Standard Fourier Shell Correlation (GSFSC) plot for the structure of the EvVS core and one high-resolution cyclase (Map 1). The resolution estimate at GSFSC = 0.143 is 3.00 Å. (B) Angular distribution and GSFSC plot for the structure of the EvVS core and one high-resolution cyclase (Map 2). The resolution estimate at GSFSC = 0.143 is 3.08 Å. (C) Angular distribution and GSFSC plot for the structure of the EvVS core and one high-resolution cyclase (Map 3). The resolution estimate at GSFSC = 0.143 is 2.98 Å.

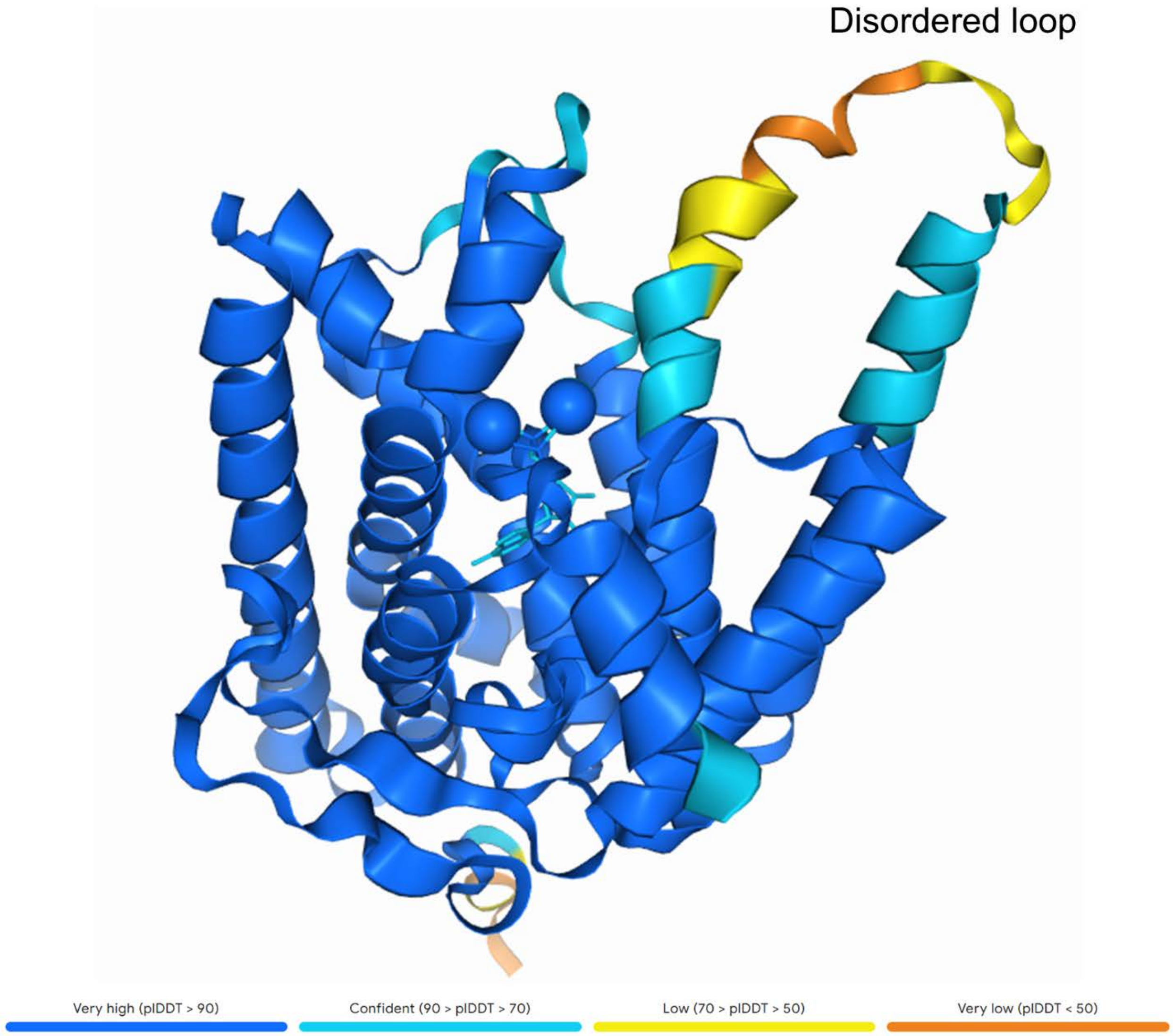


**Figure S8. AlphaFold-predicted structure of the EvVS cyclase domain colored by prediction confidence.** AlphaFold correctly identifies a loop connecting two helices adjacent to the active site as a segment of low fold-stability. This loop is not modeled in the cryo-EM structure of the cyclase domain due to disorder (identified in **Figure 7A**). Magnesium ions and a bound ligand, adenosine diphosphate (ADP), are visible in the active site. ADP was the only ligand in the AlphaFold library containing a diphosphate group so it was used as a stand-in for substrate GGPP in the structure prediction.

```
EvVS   PDHRVNQLKSIAQTLHNASLMLDDIEDHSPLRRGRPSTHMIFGTEQTINSANFLLIDVME
MpMS   PARSVSIIKDVVKHIHTASLLCDDIEDSSPLRRGQPSAHIIFGVSQTVNSTSYLWTLAID
CoSS   PSAQLGAVKRVVDVLHNSSLILDDIQDDSPMRRGKTATHLVFGAAQAINSATFLHVRAVR
CsSS   SGPAISVIKEVIDCLHHSSLILDDIEDGSHLRRGFPATHVVYGTCQAVNSATFLYVQAVE
ZbSS   PAGSSASIKSIIGMLHQSSLMLDDIEDDSTLRRGKPTAHTLFGIAQTINSANWVFACAFE
PfVS   PERSLATIRKIVNLLHSSSLMLDDIEDNSPLRRGLPATHTVFGISQTINSANLLMFKALK
NfSS   DETSLTKIRRLVDLLHNASLILDDIEDHSPKRRGRPATHTIFGHSQAINSANFMFVQAVQ
PbSS   PAPSMQIIKNIVQMLHNSSLMLDDIEDESPLRRGQPVAHTFYGISQTINSANFVYVKSVK
PvPS   PEAELEVIKEAIDLLHNSSLMLDDIEDDSPLRRGFPSTHVVYGISQTINSANYLYVMALE
EvQS   PPLALSSIKRIVEYLHHSSLMLDDIEDNSTLRRGKPCTHMLYGNAQTINAANYAFVSAFA
CgDS   PPDALATIRTIIRIMHNASLMLDDVQDNSPVRRGSPSAHVIFGTAQTTNSASYLMIKCVD
AcSS   PRSSLAAISGVASLLHEASLMLDDIQDGSPLRRGQPAVHEMFGVGQTINSACYCINNALR
FoFS   PPVVLGHISSAIDMLHNASLILDDIQDNSPLRRGVPAAHVVFGTAQSINSATFMFVKATE
TpcA   PSERLDSIMSVINTLHNASLILDDLEDNSPLRRGYPATHILFGHSQSINTANFMFVRAVQ
FgMS   PIENANTIKAITESLHGSSLMLDDIEDHSQLRRGKPSAHAVFGEAQTINSATFQYIQSVS
AcOS   PSTSTSTIKDLIKKLHSASLMLDDIEDNSPLRRAKPSTHIIYGNAQTINSATYQYTEATS
PaPS   PAKELNQIKRAINLLHNASLMLDDVQDGSVLRRAQPTTHTVFGPAQTINSAGHQIIQAMN
PrDS   PVKSLLIIEGAVNFLHNSSLLLDDIQDGSVLRRGRPVAHQIFGVGQTINTATYLMNEALY
PaFS   PDVKVGKIKDAVRVLHNSSLLLDDFQDNSPLRRGKPSTHNIFGSAQTVNTATYSIIKAIG
EvAS   PAEKLDLIKSITTILHNASLMLDDVEDGSELRRGNPSTHTIFGLSQTINSANYQLVRALE
TnDC   PAAKLDLIKLITNMLHNTSLMLDDLEDGSHLRRGRSSTHTIFGAGQTVNAANYHVIRALE
EvSS   PTAKLEIIKSITTILHNASLMLDDVEDGSELRRGKPATHNIFGLGQTINSANYQLVRALQ

EvVS   SHLRQAGSIEYTEAKMGELMEKITDSVVSLEGETG-S-PNWVVRLLIHRLKV---------
MpMS   QIMKEAGSLEHTRKVVLELQDAVHRELAKLEEAFG-Q-ENYVIQLALERLRIKA-------
CoSS   EYLYEAGSFDACWRLLVRLEDDIEGEIRRLEEATG-E-ENPQMHLLLKLLSVKNDKPNKGP
CsSS   SILDETGSTAATKALLLKWHDEITEEIGALERHFG-V-DNALLRLLVETLRV---------
ZbSS   EKMRSGGALNATISLLKDLQDNILEELKSLESAFG-S-GNPMLELVLRRLWI---------
PfVS   DDIKATGGLKYAKKMAMSLQDSVNETLTQYEDKVG-A-KNWILRLVQKRLELEV-------
NfSS   EYLNTSGTFQHCREFLMQLESLIESEIDRIEKVTN-E-ANPMLRLLLEKLSVKEN------
PbSS   QAMDEAGTFEYAQGVLKYLHEEIMRTLDEVEADLG---RNTEARILLLGLGL---------
PvPS   DHLAETKSLEYTKEMLGYMYTQLQKEVDFLERQTG-S-ENFLLRLLLKRLQV---------
EvQS   SQMEEKGALSATHSLLQKMQKELIEGLHRVEETFG-S-KNALVELMLRRLWV---------
CgDS   EMLEEAGSLEHTRVVIRGLYDETRAVLTAMENEAGSGGKNWMLHLITFQLKV---------
AcSS   SQIRGSGSFEYTKELLSHLLYDLEDMVRDMESVTG-Q-KNWILRNILVQMRVKEERAVQKK
FoFS   RCIAASGGFDETLKCLRSLENELDTEIAELEKKLG-Q-VNPLLRLCLATLSMEGCE-KICW
TpcA   ACLKKSGAFNKTIACLTDMERDLEFEINRLEQQTG-E-TNPMLRLCLAKLSVKGIG-RIGE
FgMS   NLIEEAGGISGTEKVLHSLYGEMEAELERLAGVFG-A-ENHQLELILEMLRID--------
AcOS   GIMKKTKSLEYTLGVLRALQEELEREVGRLEGKFG-E-ENLPLRLMVDMLKV---------
PaPS   SIIEGARSLEYTAAVLQKLYKAIVRELESTERQFG---ENKPFRFLLSLLKV---------
PrDS   EEITARGAFSQTKAVLRKLHIELLRLLMETEQKAG-GIENWALRLLIMKLDLGDEK---KK
PaFS   IIEEEKSLDYTRSVMMDLHVQLRAEIGRIEILLD-S-PNPAMRLLLELLRV----------
EvAS   LMEKSGSLKFTRETLASLYSGLEKSFTELEEKFG-T-ENFQLKLILQFLRTE---------
TnDC   LMKENGSLQFTMDALDVLHAKVEKSISDLEARFG-V-ENFQLRLILEMLRKA---------
EvSS   LMKEAGTLKFTQDSLGVLYSDVEKSVAELESKFG-I-ENFQLRLIMELLKTG---------
```

**Figure S9. Multiple Sequence Alignment of Prenyltransferases.** Alignment of 22 class I bifunctional terpene synthases. Above shows a region of highest identity among the sequences, with the conserved metal binding DDXXD motif of the prenyltransferase domain highlighted in blue (residues 447-506 of EvVS at top. Below shows the alignment at the C-terminal helix of EvVS (residues 671-725 for EvVS) with residues that form the cyclase domain interaction highlighted in red. These residues are generally not conserved among the sequences. Proteins are listed by ipTM score in descending order (see **Table S2**). Alignments were generated using Kalign, available on the European Bioinformatics Institute website, https://www.ebi.ac.uk/jdispatcher/msa/kalign.

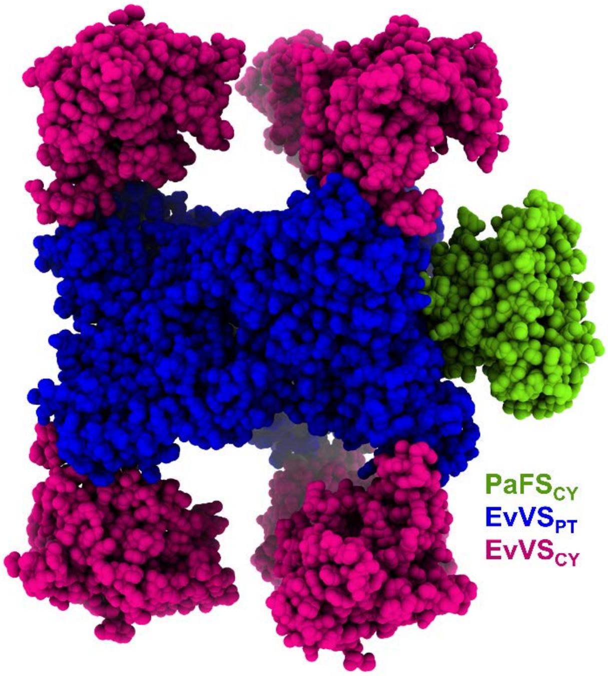


**Figure S10. Model of $PaFS_{CY}$ docked at the side of the EvVS prenyltransferase hexamer.** $PaFS_{CY}$ (green) is capable of docking at the side of the complete EvVS assembly (prenyltransferase domains = blue, cyclase domains = magenta). Such a binding interaction might support GGPP channeling as indicated by substrate competition experiments.

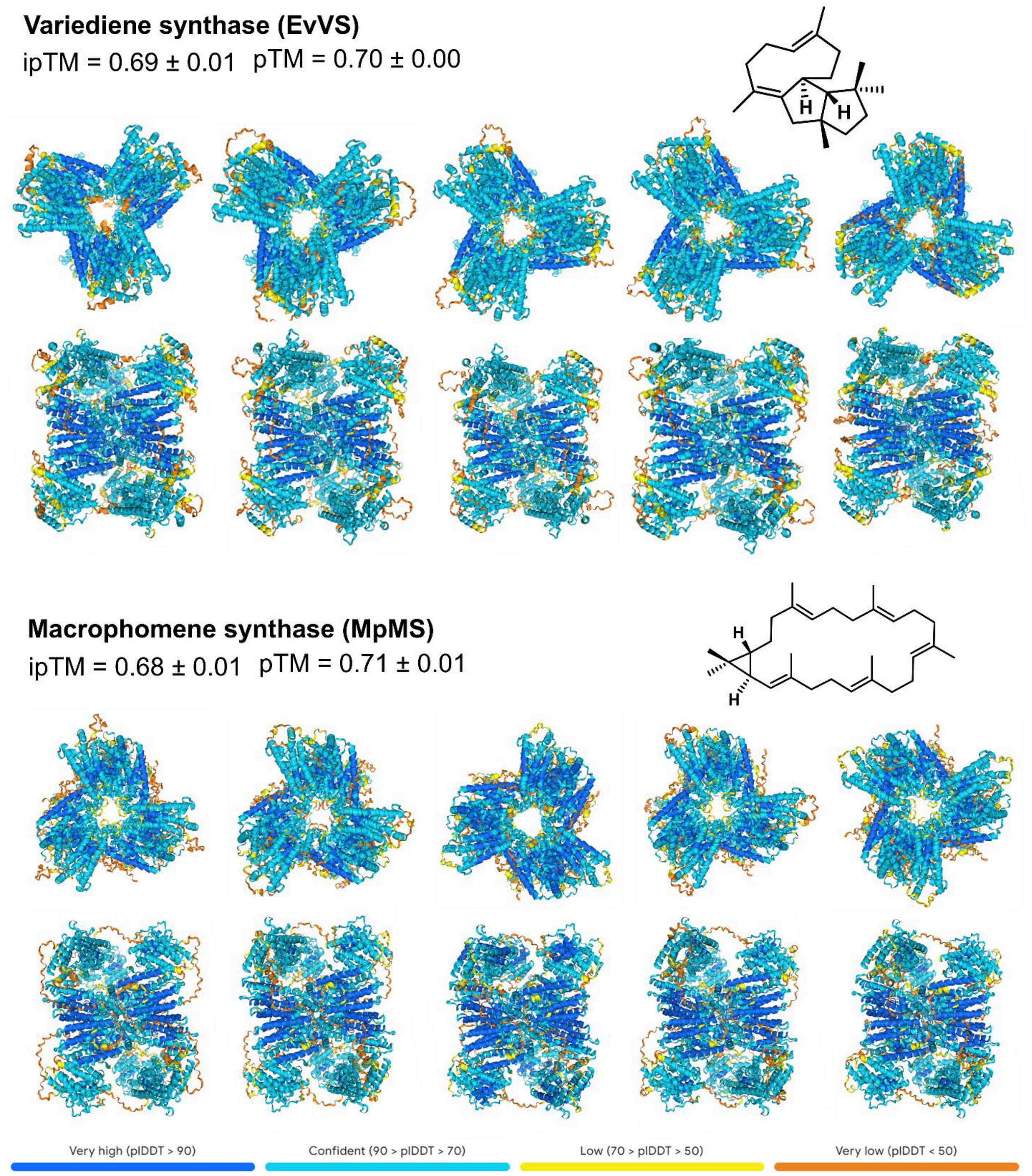


**Figure S11. AlphaFold predictions for EvVS and MpMS.** Five predictions generated by AlphaFold for the hexameric forms of EvVS and MpMS and their average ipTM and pTM scores. The structures are colored according to prediction confidence (legend at bottom). Top and side views are shown for each predicted structure. The structures of their respective products are shown at right.

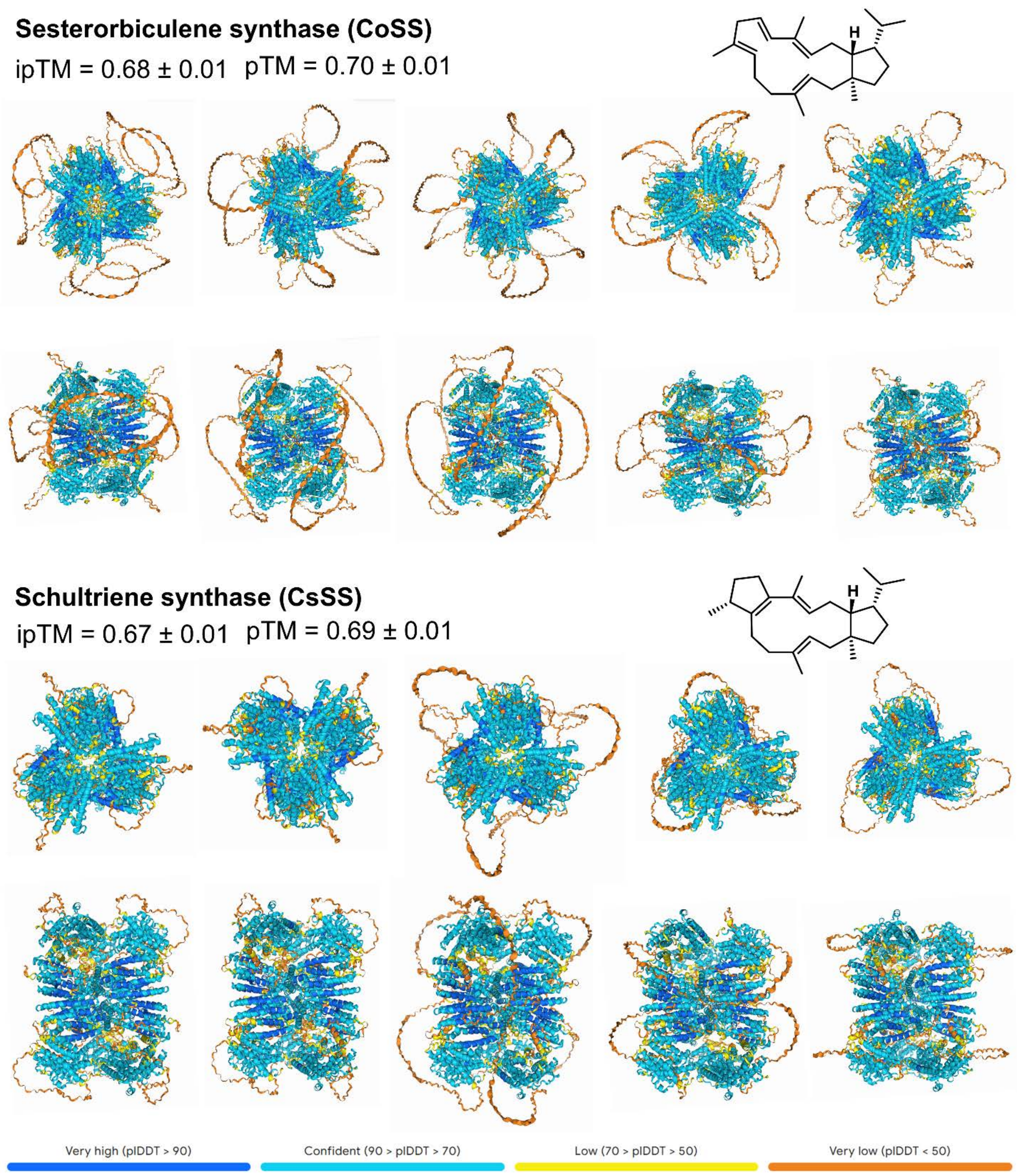


**Figure S12. AlphaFold predictions for CoSS and CsSS.** Five predictions generated by AlphaFold for the hexameric forms of CoSS and CsSS and their average ipTM and pTM scores. The structures are colored according to prediction confidence (legend at bottom). Top and side views are shown for each predicted structure. The structures of their respective products are shown at right.

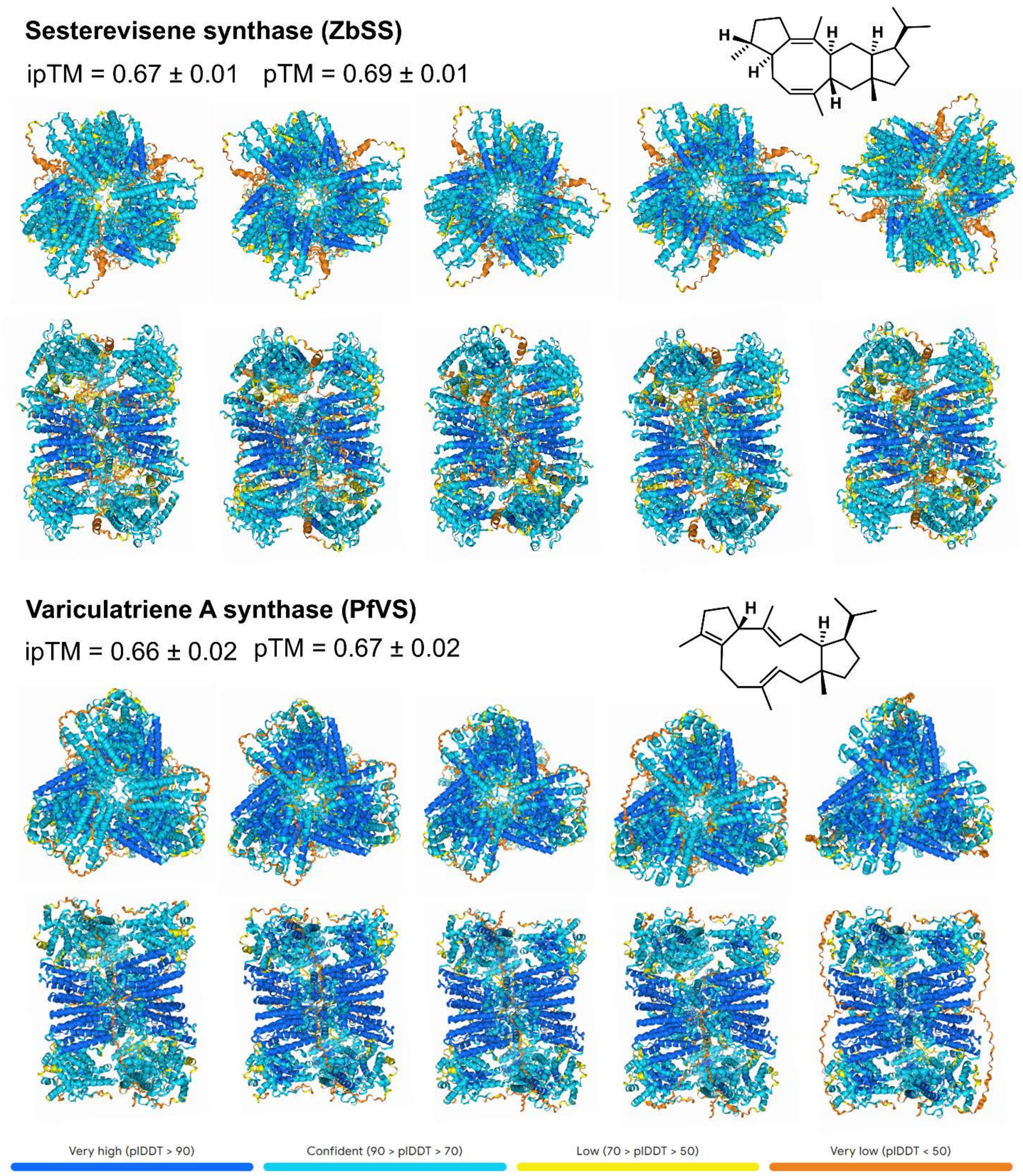


**Figure S13. AlphaFold predictions for ZbSS and PfVS.** Five predictions generated by AlphaFold for the hexameric forms of ZbSS and PfVS and their average ipTM and pTM scores. The structures are colored according to prediction confidence (legend at bottom). Top and side views are shown for each predicted structure. The structures of their respective products are shown at right.

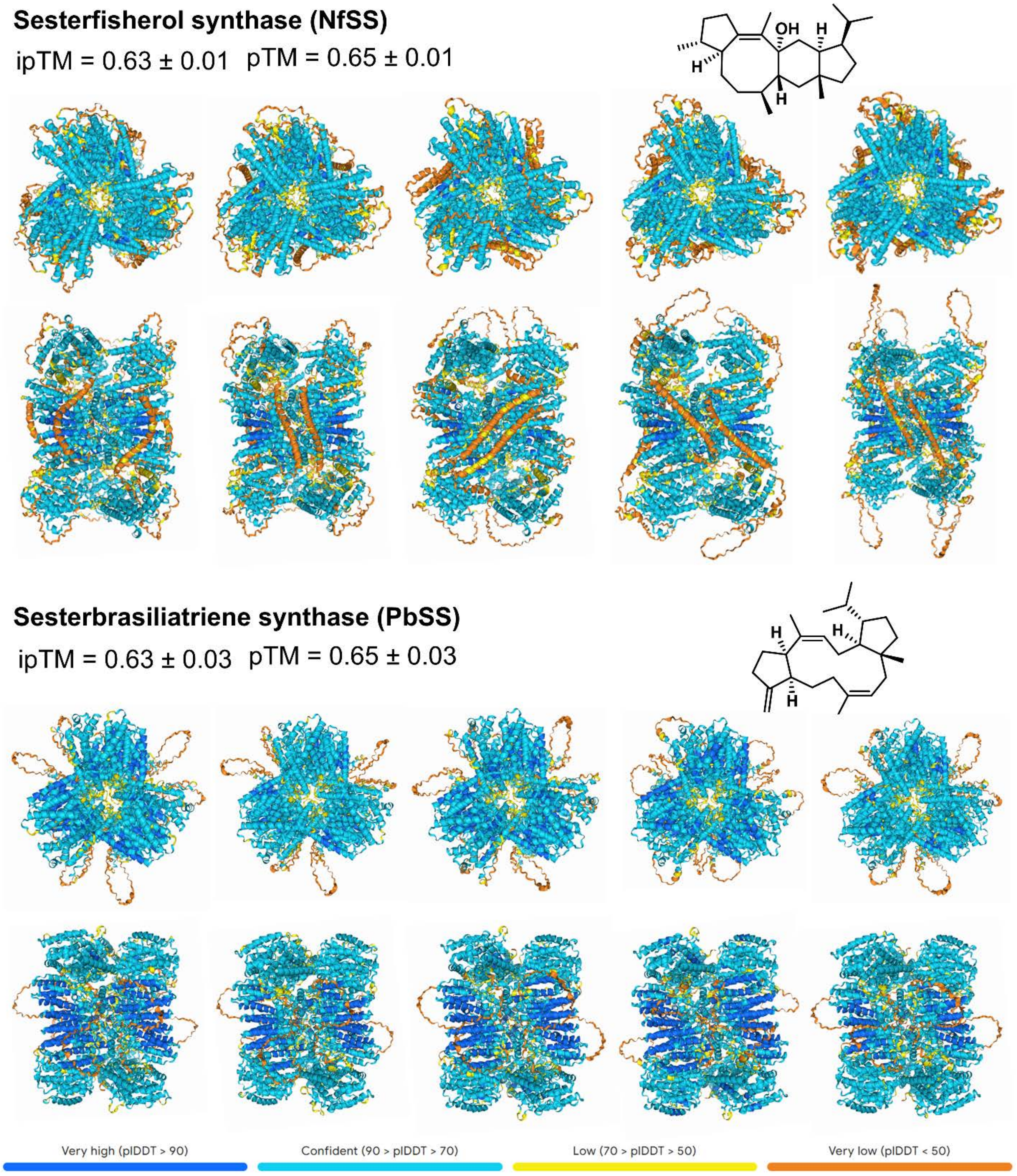


**Figure S14. AlphaFold predictions for NfSS and PbSS.** Five predictions generated by AlphaFold for the hexameric forms of NfSS and PbSS and their average ipTM and pTM scores. The structures are colored according to prediction confidence (legend at bottom). Top and side views are shown for each predicted structure. The structures of their respective products are shown at right.

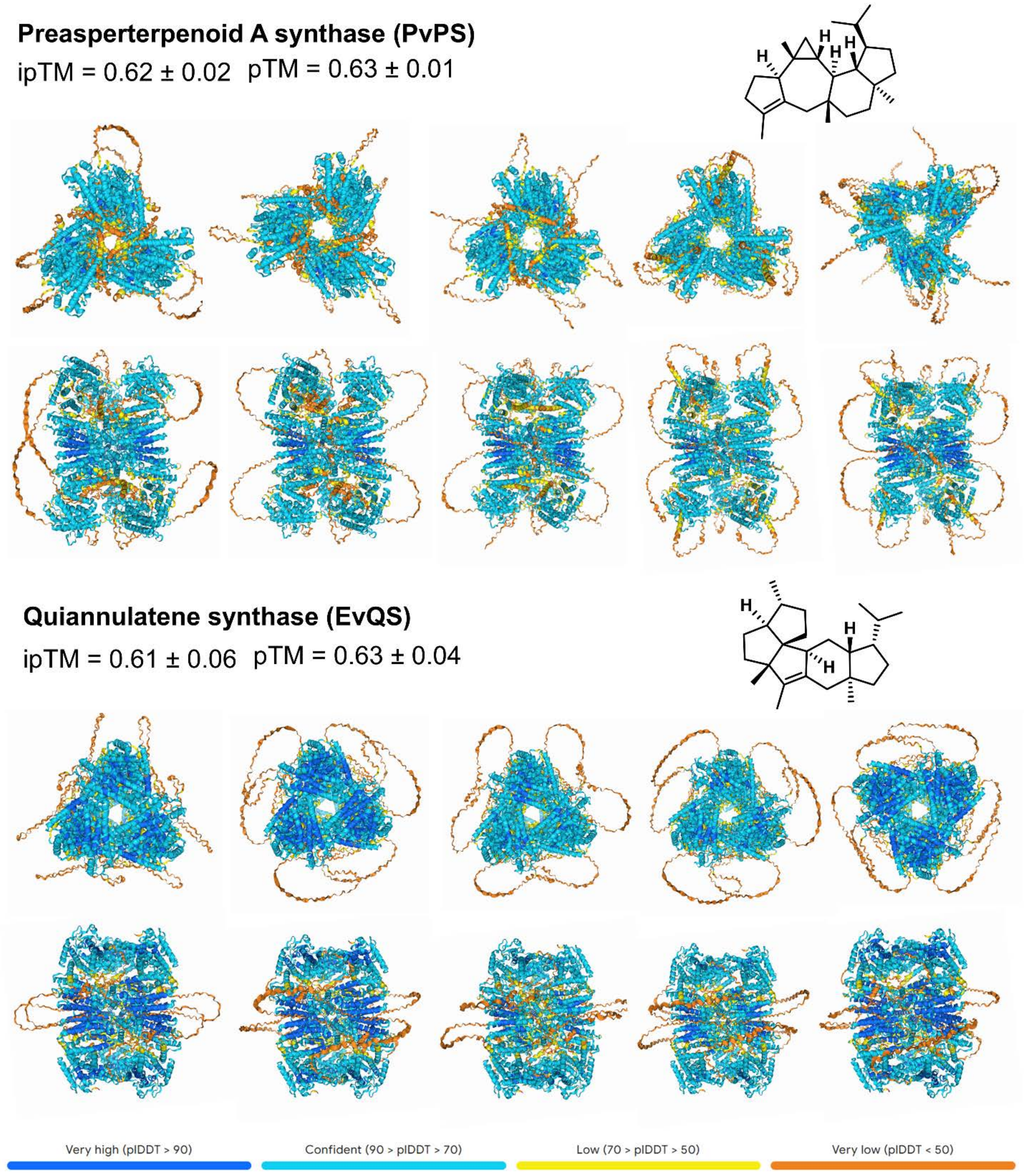


**Figure S15. AlphaFold predictions for PvPS and EvQS.** Five predictions generated by AlphaFold for the hexameric forms of PvPS and EvQS and their average ipTM and pTM scores. The structures are colored according to prediction confidence (legend at bottom). Top and side views are shown for each predicted structure. The structures of their respective products are shown at right.

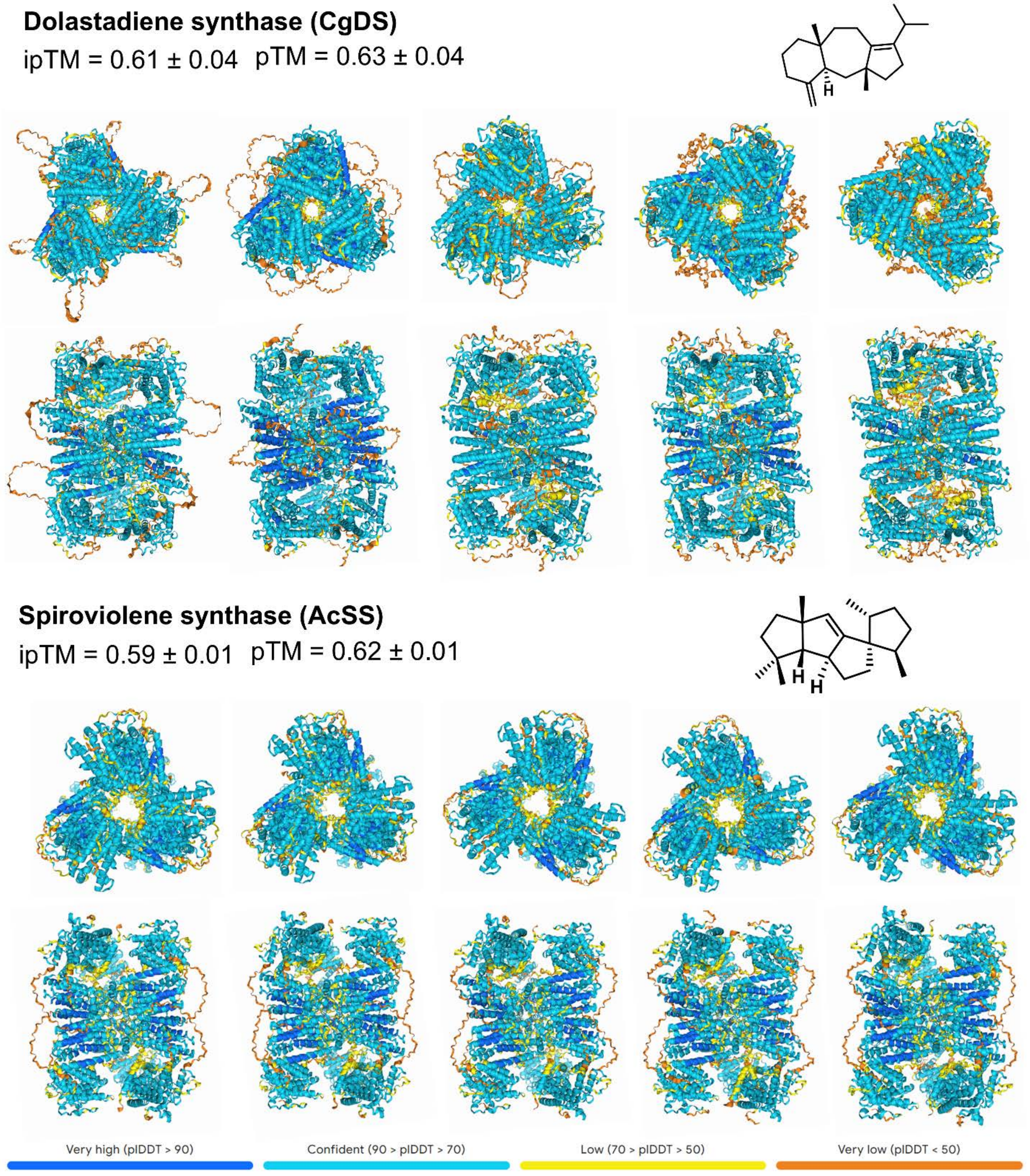


**Figure S16. AlphaFold predictions for CgDS and AcSS.** Five predictions generated by AlphaFold for the hexameric forms of CgDS and AcSS and their average ipTM and pTM scores. The structures are colored according to prediction confidence (legend at bottom). Top and side views are shown for each predicted structure. The structures of their respective products are shown at right.

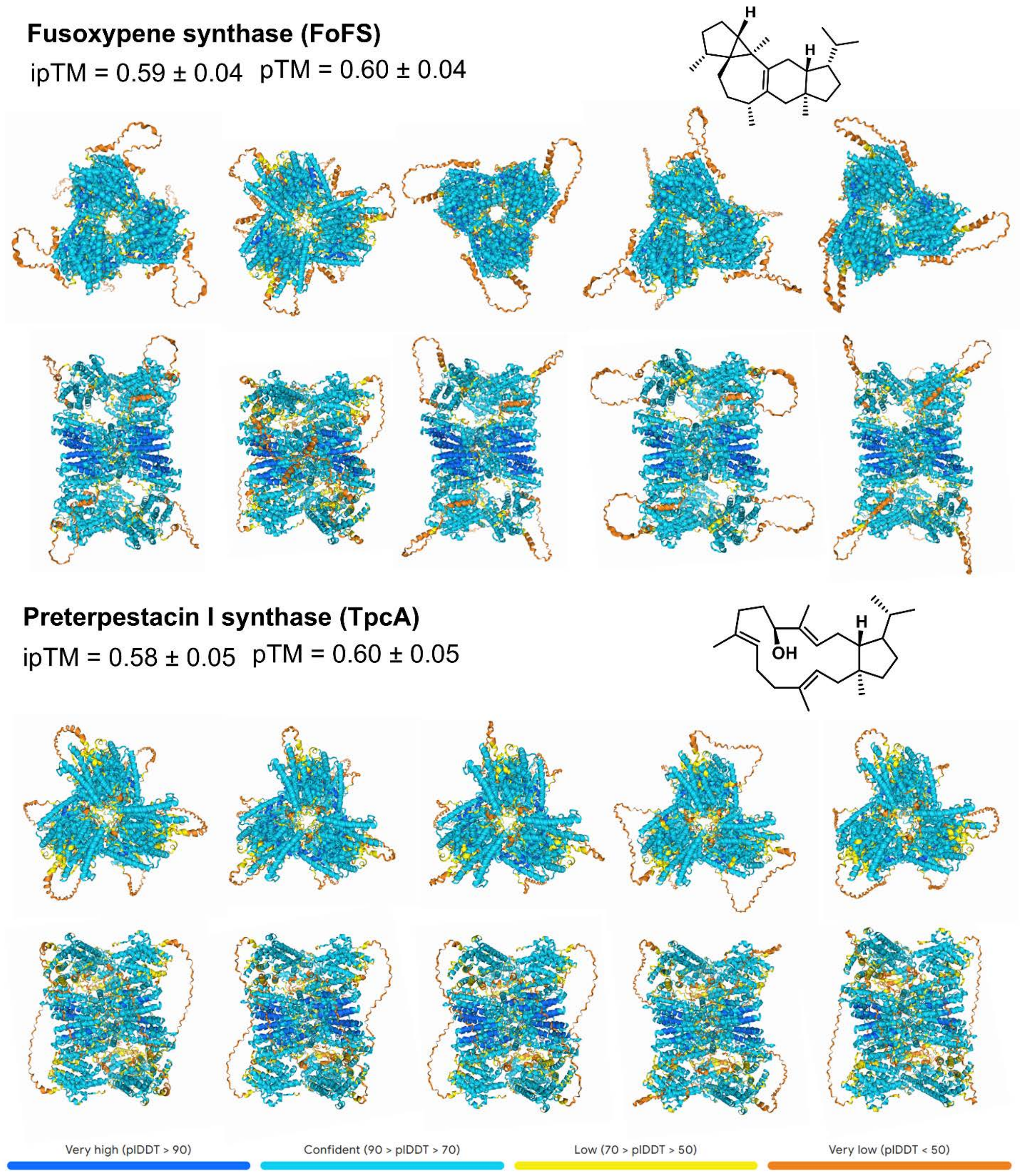


**Figure S17. AlphaFold predictions for FoFS and TpcA.** Five predictions generated by AlphaFold for the hexameric forms of FoFS and TpcA and their average ipTM and pTM scores. The structures are colored according to prediction confidence (legend at bottom). Top and side views are shown for each predicted structure. The structures of their respective products are shown at right.

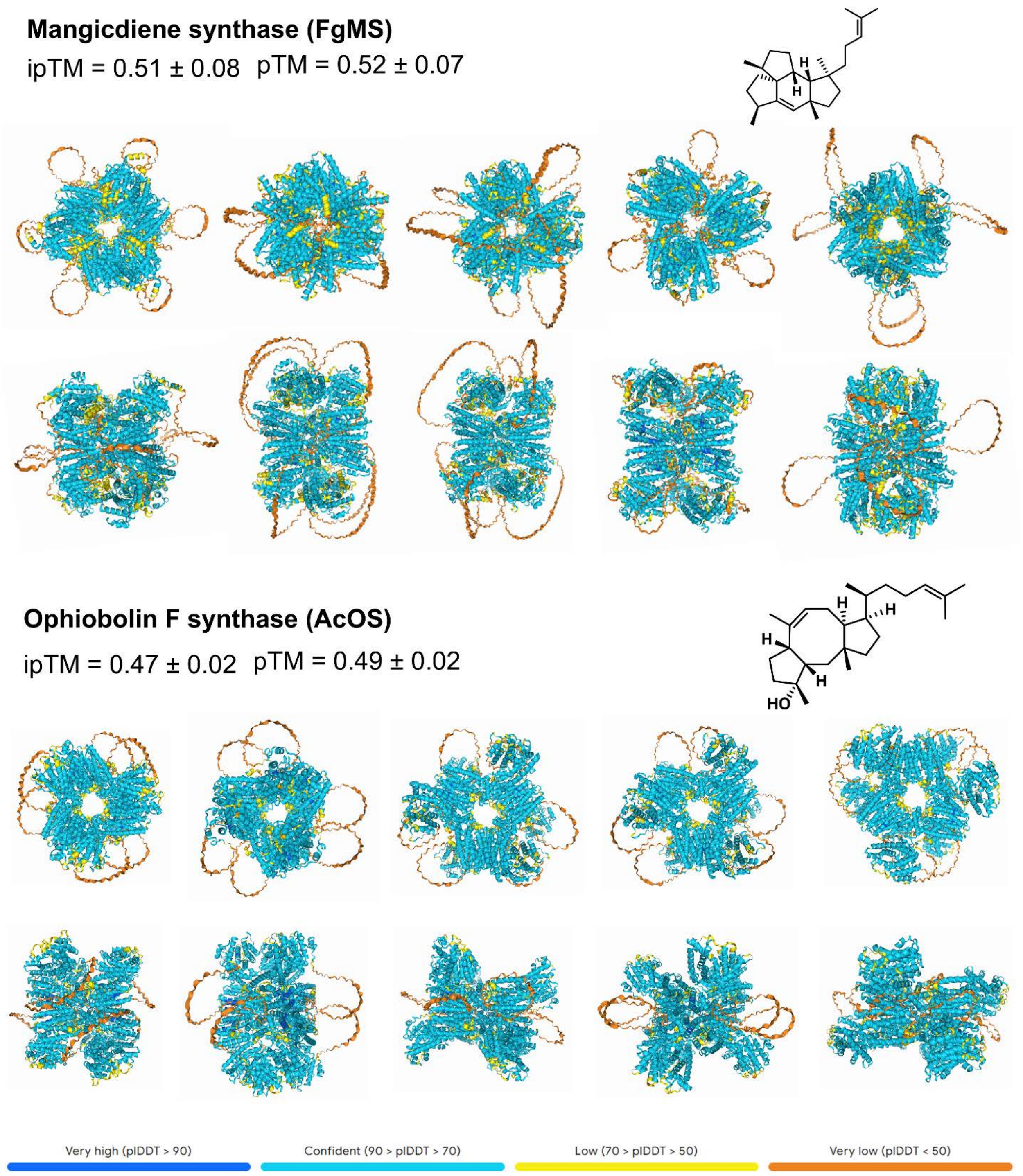


**Figure S18. AlphaFold predictions for FgMS and AcOS.** Five predictions generated by AlphaFold for the hexameric forms of FgMS and AcOS and their average ipTM and pTM scores. The structures are colored according to prediction confidence (legend at bottom). Top and side views are shown for each predicted structure. The structures of their respective products are shown at right.

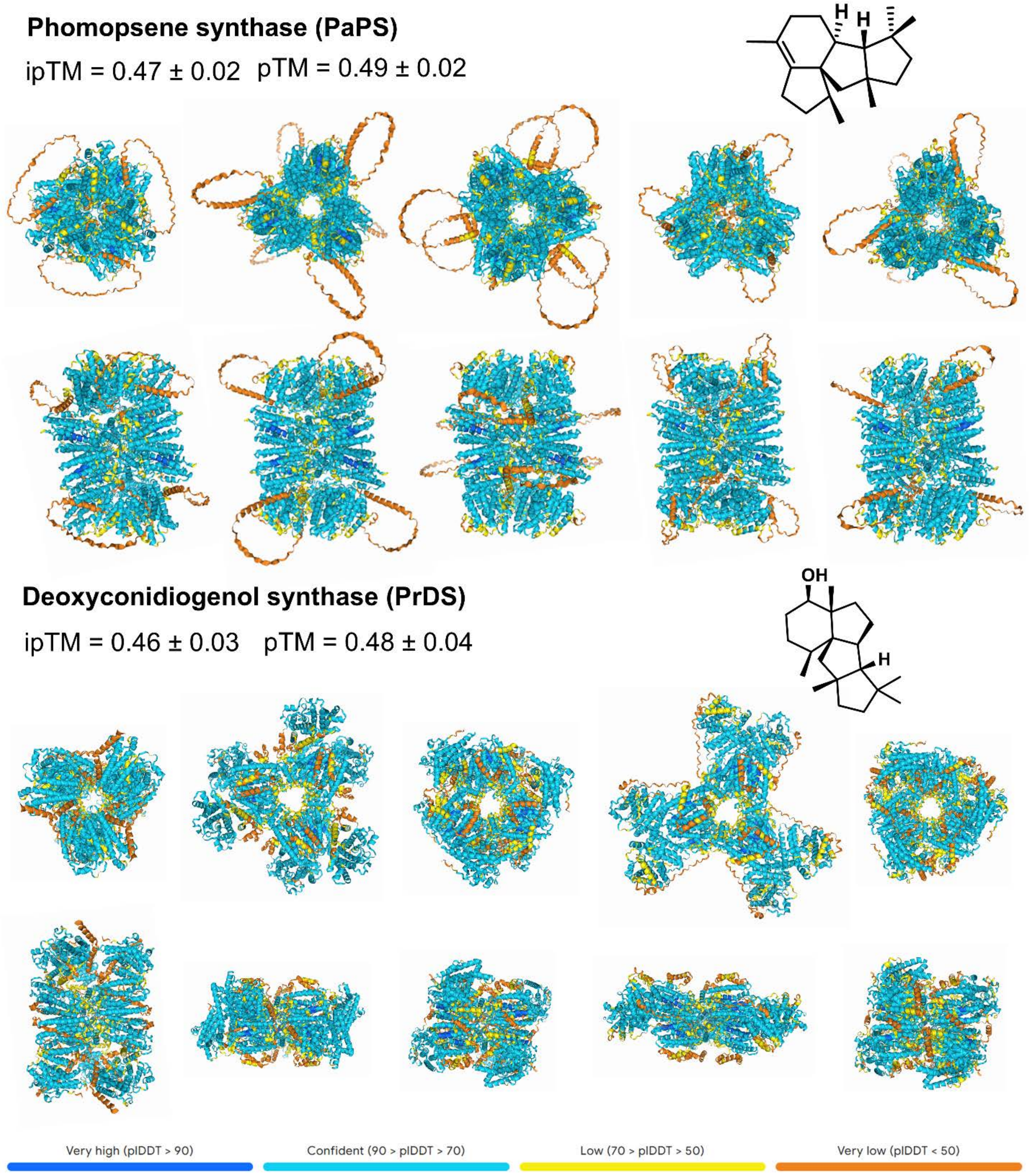


**Figure S19. AlphaFold predictions for PaPS and PrDS.** Five predictions generated by AlphaFold for the hexameric forms of PaPS and PrDS and their average ipTM and pTM scores. The structures are colored according to prediction confidence (legend at bottom). Top and side views are shown for each predicted structure. The structures of their respective products are shown at right.

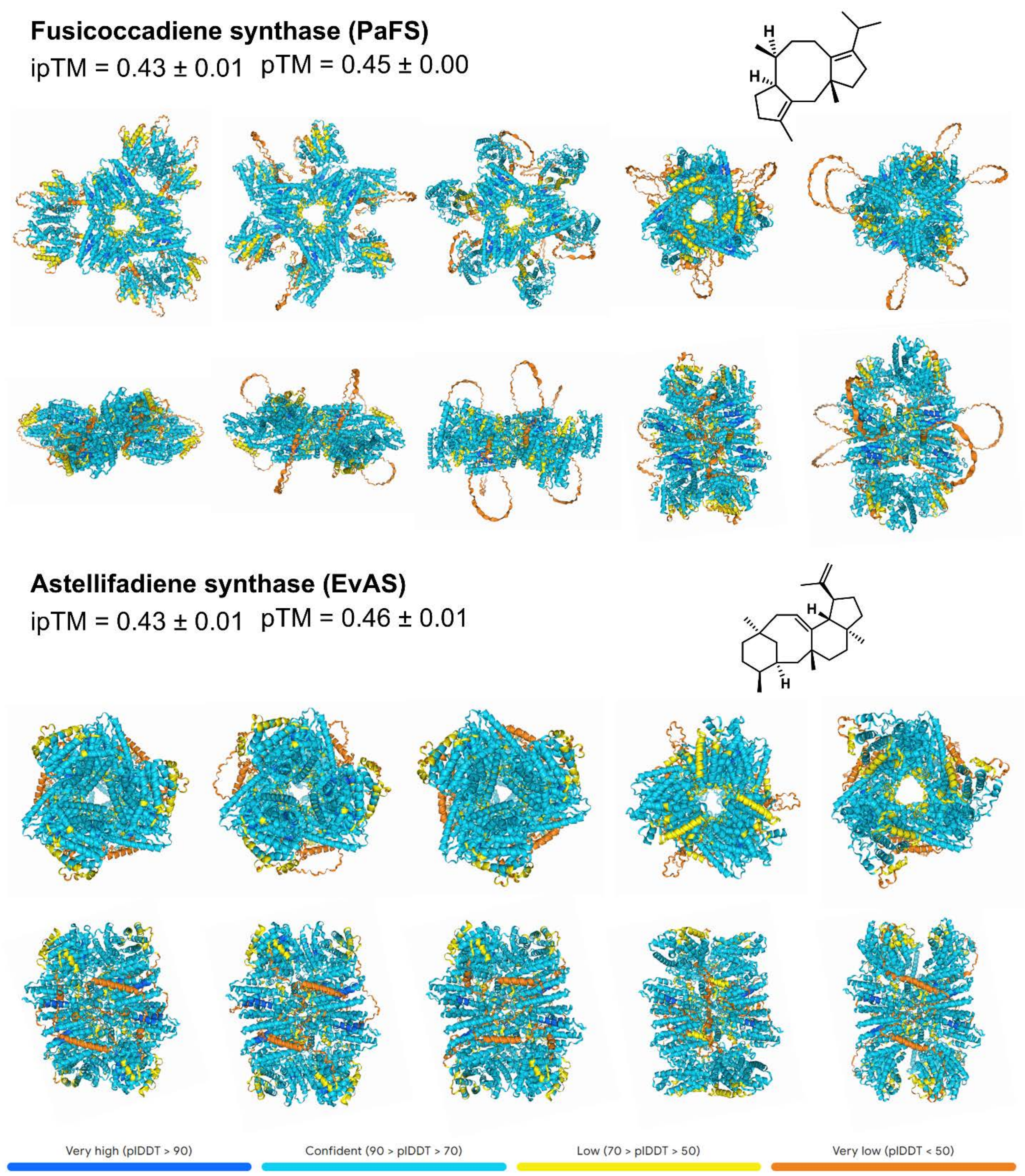


**Figure S20. AlphaFold predictions for PaFS and EvAS.** Five predictions generated by AlphaFold for the hexameric forms of PaFS and EvAS and their average ipTM and pTM scores. The structures are colored according to prediction confidence (legend at bottom). Top and side views are shown for each predicted structure. The structures of their respective products are shown at right.

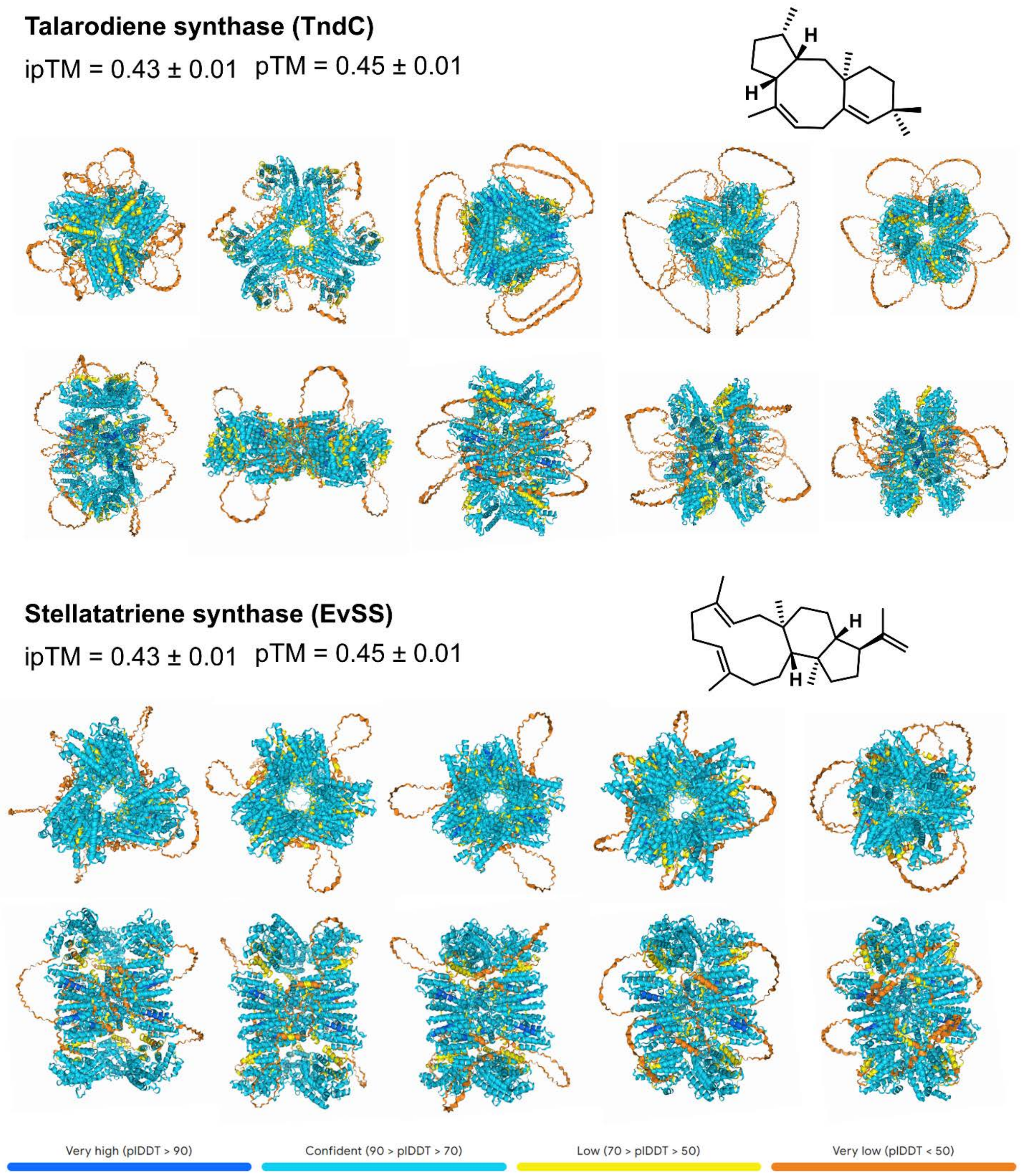


**Figure S21. AlphaFold predictions for TndC and EvSS.** Five predictions generated by AlphaFold for the hexameric forms of TndC and EvSS and their average ipTM and pTM scores. The structures are colored according to prediction confidence (legend at bottom). Top and side views are shown for each predicted structure. The structures of their respective products are shown at right.

### Full Length

| | EvVS | MpMS | CoSS | CsSS | ZbSS | PfVS | NfSS | PbSS | PvPS | EvQS | CgDS | AcSS | FoFS | TpcA | FgMS | AcOS | PaPS | PrDS | PaFS | EvAS | TnDC | EvSS |
|---|---|---|---|---|---|---|---|---|---|---|---|---|---|---|---|---|---|---|---|---|---|---|
| **EvVS** | 100.0 | 40.0 | 31.3 | 28.7 | 29.4 | 31.7 | 30.6 | 30.0 | 37.1 | 30.3 | 28.1 | 30.7 | 27.5 | 30.8 | 33.9 | 34.4 | 33.2 | 31.2 | 34.5 | 34.2 | 34.6 | 33.4 |
| **MpMS** | 40.0 | 100.0 | 29.4 | 27.0 | 30.8 | 31.8 | 27.9 | 30.2 | 37.0 | 28.4 | 26.4 | 28.8 | 28.5 | 30.2 | 34.8 | 36.0 | 32.0 | 29.6 | 32.8 | 32.3 | 32.5 | 32.9 |
| **CoSS** | 31.3 | 29.4 | 100.0 | 33.0 | 29.3 | 29.7 | 37.6 | 28.5 | 31.7 | 27.6 | 27.2 | 27.2 | 35.1 | 39.4 | 26.7 | 30.9 | 29.4 | 28.1 | 31.2 | 27.6 | 28.5 | 29.1 |
| **CsSS** | 28.7 | 27.0 | 33.0 | 100.0 | 29.2 | 26.5 | 34.7 | 26.4 | 31.0 | 29.1 | 22.2 | 25.9 | 36.2 | 37.6 | 27.0 | 29.1 | 27.1 | 26.5 | 26.7 | 26.7 | 28.0 | 26.4 |
| **ZbSS** | 29.4 | 30.8 | 29.3 | 29.2 | 100.0 | 30.0 | 32.3 | 31.3 | 32.9 | 47.9 | 26.2 | 29.8 | 28.4 | 31.9 | 30.9 | 29.8 | 29.7 | 29.4 | 29.1 | 28.3 | 30.2 | 30.1 |
| **PfVS** | 31.7 | 31.8 | 29.7 | 26.5 | 30.0 | 100.0 | 27.5 | 27.3 | 36.4 | 29.8 | 31.6 | 35.9 | 28.7 | 29.9 | 30.6 | 30.0 | 29.5 | 37.1 | 28.6 | 30.3 | 30.1 | 30.8 |
| **NfSS** | 30.6 | 27.9 | 37.6 | 34.7 | 32.3 | 27.5 | 100.0 | 28.8 | 29.3 | 28.3 | 26.3 | 29.7 | 37.7 | 45.3 | 28.2 | 30.1 | 26.9 | 29.4 | 28.6 | 26.9 | 27.8 | 28.7 |
| **PbSS** | 30.0 | 30.2 | 28.5 | 26.4 | 31.3 | 27.3 | 28.8 | 100.0 | 33.1 | 28.7 | 26.1 | 28.2 | 30.4 | 28.5 | 30.2 | 30.6 | 31.3 | 29.5 | 28.0 | 30.0 | 30.2 | 30.2 |
| **PvPS** | 37.1 | 37.0 | 31.7 | 31.0 | 32.9 | 36.4 | 29.3 | 33.1 | 100.0 | 31.3 | 28.6 | 30.5 | 31.8 | 33.2 | 40.3 | 40.1 | 41.2 | 32.8 | 40.6 | 40.6 | 43.3 | 41.1 |
| **EvQS** | 30.3 | 28.4 | 27.6 | 29.1 | 47.9 | 29.8 | 28.3 | 28.7 | 31.3 | 100.0 | 24.5 | 28.7 | 28.5 | 30.6 | 29.2 | 28.5 | 28.2 | 29.9 | 28.7 | 29.7 | 29.2 | 29.6 |
| **CgDS** | 28.1 | 26.4 | 27.2 | 22.2 | 26.2 | 31.6 | 26.3 | 26.1 | 28.6 | 24.5 | 100.0 | 31.6 | 26.7 | 27.0 | 27.0 | 27.7 | 26.9 | 30.9 | 28.5 | 27.3 | 27.3 | 26.8 |
| **AcSS** | 30.7 | 28.8 | 27.2 | 25.9 | 29.8 | 35.9 | 29.7 | 28.2 | 30.5 | 28.7 | 31.6 | 100.0 | 28.0 | 28.6 | 29.7 | 30.6 | 30.2 | 40.1 | 28.3 | 30.3 | 29.8 | 30.1 |
| **FoFS** | 27.5 | 28.5 | 35.1 | 36.2 | 28.4 | 28.7 | 37.7 | 30.4 | 31.8 | 28.5 | 26.7 | 28.0 | 100.0 | 54.6 | 30.1 | 32.4 | 29.9 | 29.5 | 27.3 | 28.6 | 27.7 | 28.8 |
| **TpcA** | 30.8 | 30.2 | 39.4 | 37.6 | 31.9 | 29.9 | 45.3 | 28.5 | 33.2 | 30.6 | 27.0 | 28.6 | 54.6 | 100.0 | 29.5 | 32.5 | 27.8 | 29.6 | 28.7 | 30.4 | 32.6 | 30.3 |
| **FgMS** | 33.9 | 34.8 | 26.7 | 27.0 | 30.9 | 30.6 | 28.2 | 30.2 | 40.3 | 29.2 | 27.0 | 29.7 | 30.1 | 29.5 | 100.0 | 44.9 | 38.6 | 31.4 | 41.4 | 40.4 | 40.8 | 40.1 |
| **AcOS** | 34.4 | 36.0 | 30.9 | 29.1 | 29.8 | 30.0 | 30.1 | 30.6 | 40.1 | 28.5 | 27.7 | 30.6 | 32.4 | 32.5 | 44.9 | 100.0 | 41.0 | 30.5 | 41.6 | 38.2 | 42.9 | 41.9 |
| **PaPS** | 33.2 | 32.0 | 29.4 | 27.1 | 29.7 | 29.5 | 26.9 | 31.3 | 41.2 | 28.2 | 26.9 | 30.2 | 29.9 | 27.8 | 38.6 | 41.0 | 100.0 | 30.2 | 37.2 | 38.6 | 39.9 | 39.3 |
| **PrDS** | 31.2 | 29.6 | 28.1 | 26.5 | 29.4 | 37.1 | 29.4 | 29.5 | 32.8 | 29.9 | 30.9 | 40.1 | 29.5 | 29.6 | 31.4 | 30.5 | 30.2 | 100.0 | 32.1 | 30.8 | 32.6 | 31.7 |
| **PaFS** | 34.5 | 32.8 | 31.2 | 26.7 | 29.1 | 28.6 | 28.6 | 28.0 | 40.6 | 28.7 | 28.5 | 28.3 | 27.3 | 28.7 | 41.4 | 41.6 | 37.2 | 32.1 | 100.0 | 40.4 | 42.9 | 42.0 |
| **EvAS** | 34.2 | 32.3 | 27.6 | 26.7 | 28.3 | 30.3 | 26.9 | 30.0 | 40.6 | 29.7 | 27.3 | 30.3 | 28.6 | 30.4 | 40.4 | 38.2 | 38.6 | 30.8 | 40.4 | 100.0 | 55.2 | 59.0 |
| **TnDC** | 34.6 | 32.5 | 28.5 | 28.0 | 30.2 | 30.1 | 27.8 | 30.2 | 43.3 | 29.2 | 27.3 | 29.8 | 27.7 | 32.6 | 40.8 | 42.9 | 39.9 | 32.6 | 42.9 | 55.2 | 100.0 | 58.3 |
| **EvSS** | 33.4 | 32.9 | 29.1 | 26.4 | 30.1 | 30.8 | 28.7 | 30.2 | 41.1 | 29.6 | 26.8 | 30.1 | 28.8 | 30.3 | 40.1 | 41.9 | 39.3 | 31.7 | 42.0 | 59.0 | 58.3 | 100.0 |

### Prenyltransferase Domain

| | EvVS | MpMS | CoSS | CsSS | ZbSS | PfVS | NfSS | PbSS | PvPS | EvQS | CgDS | AcSS | FoFS | TpcA | FgMS | AcOS | PaPS | PrDS | PaFS | EvAS | TnDC | EvSS |
|---|---|---|---|---|---|---|---|---|---|---|---|---|---|---|---|---|---|---|---|---|---|---|
| **EvVS** | 100.0 | 45.9 | 40.5 | 41.6 | 43.1 | 41.6 | 43.5 | 43.0 | 47.6 | 40.9 | 37.3 | 41.1 | 39.8 | 42.3 | 45.9 | 47.8 | 45.9 | 40.1 | 42.0 | 47.3 | 45.1 | 46.8 |
| **MpMS** | 45.9 | 100.0 | 40.7 | 38.0 | 45.2 | 43.9 | 39.8 | 44.2 | 49.7 | 42.0 | 37.9 | 39.1 | 37.0 | 38.9 | 50.3 | 50.5 | 43.5 | 39.3 | 42.0 | 44.6 | 42.9 | 43.2 |
| **CoSS** | 40.5 | 40.7 | 100.0 | 44.6 | 41.4 | 38.2 | 50.8 | 40.1 | 43.4 | 40.5 | 35.0 | 36.0 | 44.5 | 49.4 | 39.7 | 39.6 | 43.2 | 34.9 | 39.6 | 40.2 | 43.9 | 42.6 |
| **CsSS** | 41.6 | 38.0 | 44.6 | 100.0 | 37.8 | 34.8 | 45.5 | 38.8 | 44.0 | 43.2 | 29.2 | 37.2 | 43.1 | 41.6 | 40.9 | 41.2 | 40.1 | 37.9 | 37.6 | 38.4 | 40.1 | 38.2 |
| **ZbSS** | 43.1 | 45.2 | 41.4 | 37.8 | 100.0 | 44.8 | 43.8 | 46.3 | 46.9 | 60.6 | 37.8 | 41.6 | 42.0 | 44.7 | 48.6 | 44.4 | 44.8 | 43.7 | 40.5 | 45.9 | 47.5 | 46.4 |
| **PfVS** | 41.6 | 43.9 | 38.2 | 34.8 | 44.8 | 100.0 | 40.3 | 39.5 | 48.3 | 42.6 | 38.8 | 42.6 | 39.8 | 37.7 | 46.1 | 43.9 | 41.0 | 44.6 | 37.5 | 41.7 | 40.6 | 40.5 |
| **NfSS** | 43.5 | 39.8 | 50.8 | 45.5 | 43.8 | 40.3 | 100.0 | 45.4 | 43.2 | 41.0 | 33.8 | 40.8 | 47.5 | 59.1 | 44.5 | 43.5 | 42.6 | 42.2 | 40.8 | 42.0 | 43.7 | 43.0 |
| **PbSS** | 43.0 | 44.2 | 40.1 | 38.8 | 46.3 | 39.5 | 45.4 | 100.0 | 47.8 | 46.4 | 37.9 | 37.7 | 41.8 | 39.3 | 45.6 | 44.6 | 45.1 | 43.3 | 38.1 | 42.9 | 44.6 | 43.2 |
| **PvPS** | 47.6 | 49.7 | 43.4 | 44.0 | 46.9 | 48.3 | 43.2 | 47.8 | 100.0 | 47.1 | 39.0 | 42.5 | 42.7 | 45.4 | 50.7 | 51.7 | 49.3 | 43.2 | 43.6 | 51.0 | 50.9 | 50.5 |
| **EvQS** | 40.9 | 42.0 | 40.5 | 43.2 | 60.6 | 42.6 | 41.0 | 46.4 | 47.1 | 100.0 | 35.3 | 42.9 | 41.5 | 42.1 | 47.1 | 46.0 | 43.2 | 45.6 | 41.4 | 44.8 | 48.0 | 44.3 |
| **CgDS** | 37.3 | 37.9 | 35.0 | 29.2 | 37.8 | 38.8 | 33.8 | 37.9 | 39.0 | 35.3 | 100.0 | 35.9 | 34.7 | 34.3 | 37.9 | 41.4 | 41.6 | 38.3 | 37.2 | 37.6 | 37.5 | 38.1 |
| **AcSS** | 41.1 | 39.1 | 36.0 | 37.2 | 41.6 | 42.6 | 40.8 | 37.7 | 42.5 | 42.9 | 35.9 | 100.0 | 38.6 | 39.3 | 40.9 | 43.5 | 43.7 | 48.3 | 39.9 | 43.0 | 43.5 | 43.5 |
| **FoFS** | 39.8 | 37.0 | 44.5 | 43.1 | 42.0 | 39.8 | 47.5 | 41.8 | 42.7 | 41.5 | 34.7 | 38.6 | 100.0 | 56.7 | 43.0 | 44.3 | 42.5 | 39.9 | 38.5 | 39.5 | 38.4 | 39.5 |
| **TpcA** | 42.3 | 38.9 | 49.4 | 41.6 | 44.7 | 37.7 | 59.1 | 39.3 | 45.4 | 42.1 | 34.3 | 39.3 | 56.7 | 100.0 | 41.9 | 44.4 | 42.1 | 39.9 | 39.3 | 42.1 | 43.5 | 43.5 |
| **FgMS** | 45.9 | 50.3 | 39.7 | 40.9 | 48.6 | 46.1 | 44.5 | 45.6 | 50.7 | 47.1 | 37.9 | 40.9 | 43.0 | 41.9 | 100.0 | 54.2 | 48.3 | 44.5 | 45.4 | 49.3 | 47.6 | 49.3 |
| **AcOS** | 47.8 | 50.5 | 39.6 | 41.2 | 44.4 | 43.9 | 43.5 | 44.6 | 51.7 | 46.0 | 41.4 | 43.5 | 44.3 | 44.4 | 54.2 | 100.0 | 49.7 | 43.3 | 47.7 | 48.8 | 47.1 | 49.2 |
| **PaPS** | 45.9 | 43.5 | 43.2 | 40.1 | 44.8 | 41.0 | 42.6 | 45.1 | 49.3 | 43.2 | 41.6 | 43.7 | 42.5 | 42.1 | 48.3 | 49.7 | 100.0 | 41.6 | 44.1 | 46.9 | 48.5 | 47.8 |
| **PrDS** | 40.1 | 39.3 | 34.9 | 37.9 | 43.7 | 44.6 | 42.2 | 43.3 | 43.2 | 45.6 | 38.3 | 48.3 | 39.9 | 39.9 | 44.5 | 43.3 | 41.6 | 100.0 | 42.3 | 42.2 | 43.2 | 44.6 |
| **PaFS** | 42.0 | 42.0 | 39.6 | 37.6 | 40.5 | 37.5 | 40.8 | 38.1 | 43.6 | 41.4 | 37.2 | 39.9 | 38.5 | 39.3 | 45.4 | 47.7 | 44.1 | 42.3 | 100.0 | 43.7 | 44.3 | 44.3 |
| **EvAS** | 47.3 | 44.6 | 40.2 | 38.4 | 45.9 | 41.7 | 42.0 | 42.9 | 51.0 | 44.8 | 37.6 | 43.0 | 39.5 | 42.1 | 49.3 | 48.8 | 46.9 | 42.2 | 43.7 | 100.0 | 64.2 | 72.6 |
| **TnDC** | 45.1 | 42.9 | 43.9 | 40.1 | 47.5 | 40.6 | 43.7 | 44.6 | 50.9 | 48.0 | 37.5 | 43.5 | 38.4 | 43.5 | 47.6 | 47.1 | 48.5 | 43.2 | 44.3 | 64.2 | 100.0 | 67.7 |
| **EvSS** | 46.8 | 43.2 | 42.6 | 38.2 | 46.4 | 40.5 | 43.0 | 43.2 | 50.5 | 44.3 | 38.1 | 43.5 | 39.5 | 43.5 | 49.3 | 49.2 | 47.8 | 44.6 | 44.3 | 72.6 | 67.7 | 100.0 |

**Figure S22. Multiple sequence alignment matrix of full length and prenyltransferase domains.** The sequences of 22 known class I bifunctional terpene synthases were aligned using Kalign and a matrix of their identities was generated. Cells are colored according to value; the more intense the blue color, the higher the sequence identity. The enzymes are listed in descending order by their average calculated ipTM scores (see **Table S2**). The full-length sequences generally have low identity to each other, and enzymes with higher ipTM scores do not have higher sequence identity than those enzymes without a confidently-predicted inter-domain interaction. Most of the sequence identity is driven by the prenyltransferase domains (see **Figure S23**).

### Cyclase Domain

| | EvVS | MpMS | CoSS | CsSS | ZbSS | PfVS | NfSS | PbSS | PvPS | EvQS | CgDS | AcSS | FoFS | TpcA | FgMS | AcOS | PaPS | PrDS | PaFS | EvAS | TnDC | EvSS |
|---|---|---|---|---|---|---|---|---|---|---|---|---|---|---|---|---|---|---|---|---|---|---|
| EvVS | 100.0 | 36.7 | 24.4 | 22.1 | 21.0 | 26.8 | 26.1 | 21.7 | 31.9 | 25.7 | 22.1 | 23.1 | 19.1 | 22.0 | 25.8 | 25.9 | 27.2 | 25.2 | 29.1 | 26.5 | 29.0 | 26.1 |
| MpMS | 36.7 | 100.0 | 22.9 | 20.5 | 21.8 | 24.5 | 21.4 | 21.0 | 28.8 | 20.5 | 18.6 | 21.0 | 22.6 | 23.9 | 25.2 | 25.8 | 26.0 | 23.7 | 27.5 | 24.0 | 26.9 | 27.2 |
| CoSS | 24.4 | 22.9 | 100.0 | 28.2 | 23.2 | 25.7 | 32.6 | 23.0 | 22.0 | 22.3 | 23.5 | 21.0 | 31.5 | 35.2 | 21.0 | 27.3 | 22.2 | 24.1 | 27.1 | 20.0 | 19.9 | 21.3 |
| CsSS | 22.1 | 20.5 | 28.2 | 100.0 | 23.4 | 23.0 | 29.5 | 20.8 | 22.4 | 21.8 | 18.6 | 19.4 | 33.1 | 35.4 | 20.3 | 23.9 | 20.7 | 19.4 | 21.0 | 22.2 | 24.4 | 22.1 |
| ZbSS | 21.0 | 21.8 | 23.2 | 23.4 | 100.0 | 21.5 | 27.0 | 22.9 | 24.8 | 42.2 | 20.1 | 21.5 | 20.1 | 23.5 | 21.3 | 21.6 | 21.5 | 20.2 | 22.9 | 18.0 | 21.9 | 21.2 |
| PfVS | 26.8 | 24.5 | 25.7 | 23.0 | 21.5 | 100.0 | 19.2 | 21.5 | 29.9 | 24.0 | 30.4 | 31.9 | 21.2 | 24.2 | 21.9 | 22.7 | 23.8 | 34.1 | 22.2 | 25.7 | 26.0 | 27.7 |
| NfSS | 26.1 | 21.4 | 32.6 | 29.5 | 27.0 | 19.2 | 100.0 | 21.7 | 21.4 | 23.2 | 24.3 | 21.8 | 33.5 | 37.0 | 20.7 | 22.6 | 19.4 | 20.9 | 23.2 | 17.3 | 20.1 | 21.8 |
| PbSS | 21.7 | 21.0 | 23.0 | 20.8 | 22.9 | 21.5 | 21.7 | 100.0 | 24.5 | 17.7 | 19.3 | 19.2 | 23.3 | 21.3 | 21.3 | 22.3 | 22.6 | 19.6 | 23.3 | 21.6 | 21.2 | 21.6 |
| PvPS | 31.9 | 28.8 | 22.0 | 22.4 | 24.8 | 29.9 | 21.4 | 24.5 | 100.0 | 21.9 | 24.8 | 22.3 | 22.0 | 24.8 | 35.2 | 34.0 | 36.1 | 28.5 | 42.7 | 35.4 | 41.4 | 36.0 |
| EvQS | 25.7 | 20.5 | 22.3 | 21.8 | 42.2 | 24.0 | 23.2 | 17.7 | 21.9 | 100.0 | 21.9 | 19.9 | 21.2 | 24.1 | 20.7 | 18.5 | 20.5 | 21.7 | 23.2 | 22.2 | 21.2 | 22.9 |
| CgDS | 22.1 | 18.6 | 23.5 | 18.6 | 20.1 | 30.4 | 24.3 | 19.3 | 24.8 | 21.9 | 100.0 | 28.5 | 21.9 | 22.9 | 21.0 | 20.5 | 19.5 | 28.0 | 22.3 | 20.6 | 22.2 | 19.9 |
| AcSS | 23.1 | 21.0 | 21.0 | 19.4 | 21.5 | 31.9 | 21.8 | 19.2 | 22.3 | 19.9 | 28.5 | 100.0 | 21.0 | 21.3 | 21.2 | 20.7 | 20.0 | 35.0 | 21.1 | 21.8 | 21.1 | 20.1 |
| FoFS | 19.1 | 22.6 | 31.5 | 33.1 | 20.1 | 21.2 | 33.5 | 23.3 | 22.0 | 21.2 | 21.9 | 21.0 | 100.0 | 55.9 | 21.9 | 24.5 | 22.0 | 21.3 | 20.0 | 22.5 | 21.1 | 21.4 |
| TpcA | 22.0 | 23.9 | 35.2 | 35.4 | 23.5 | 24.2 | 37.0 | 21.3 | 24.8 | 24.1 | 22.9 | 21.3 | 55.9 | 100.0 | 21.9 | 24.6 | 17.8 | 22.5 | 20.5 | 23.5 | 25.0 | 20.2 |
| FgMS | 25.8 | 25.2 | 21.0 | 20.3 | 21.3 | 21.9 | 20.7 | 21.3 | 35.2 | 20.7 | 21.0 | 21.2 | 21.9 | 21.9 | 100.0 | 43.8 | 36.1 | 22.9 | 43.3 | 39.6 | 43.0 | 40.5 |
| AcOS | 25.9 | 25.8 | 27.3 | 23.9 | 21.6 | 22.7 | 22.6 | 22.3 | 34.0 | 18.5 | 20.5 | 20.7 | 24.5 | 24.6 | 43.8 | 100.0 | 36.5 | 23.5 | 41.3 | 35.6 | 43.0 | 40.1 |
| PaPS | 27.2 | 26.0 | 22.2 | 20.7 | 21.5 | 23.8 | 19.4 | 22.6 | 36.1 | 20.5 | 19.5 | 20.0 | 22.0 | 17.8 | 36.1 | 36.5 | 100.0 | 23.2 | 35.9 | 35.0 | 36.5 | 35.9 |
| PrDS | 25.2 | 23.7 | 24.1 | 19.4 | 20.2 | 34.1 | 20.9 | 19.6 | 28.5 | 21.7 | 28.0 | 35.0 | 21.3 | 22.5 | 22.9 | 23.5 | 23.2 | 100.0 | 24.1 | 23.4 | 26.2 | 23.3 |
| PaFS | 29.1 | 27.5 | 27.1 | 21.0 | 22.9 | 22.2 | 23.2 | 23.3 | 42.7 | 23.2 | 22.3 | 21.1 | 20.0 | 20.5 | 43.3 | 41.3 | 35.9 | 24.1 | 100.0 | 43.5 | 46.5 | 44.6 |
| EvAS | 26.5 | 24.0 | 20.0 | 22.2 | 18.0 | 25.7 | 17.3 | 21.6 | 35.4 | 22.2 | 20.6 | 21.8 | 22.5 | 23.5 | 39.6 | 35.6 | 35.0 | 23.4 | 43.5 | 100.0 | 54.6 | 54.0 |
| TnDC | 29.0 | 26.9 | 19.9 | 24.4 | 21.9 | 26.0 | 20.1 | 21.2 | 41.4 | 21.2 | 22.2 | 21.1 | 21.1 | 25.0 | 43.0 | 43.0 | 36.5 | 26.2 | 46.5 | 54.6 | 100.0 | 56.9 |
| EvSS | 26.1 | 27.2 | 21.3 | 22.1 | 21.2 | 27.7 | 21.8 | 21.6 | 36.0 | 22.9 | 19.9 | 20.1 | 21.4 | 20.2 | 40.5 | 40.1 | 35.9 | 23.3 | 44.6 | 54.0 | 56.9 | 100.0 |

### Linker Region

| | EvVS | MpMS | CoSS | CsSS | ZbSS | PfVS | NfSS | PbSS | PvPS | EvQS | CgDS | AcSS | FoFS | TpcA | FgMS | AcOS | PaPS | PrDS | PaFS | EvAS | TnDC | EvSS |
|---|---|---|---|---|---|---|---|---|---|---|---|---|---|---|---|---|---|---|---|---|---|---|
| EvVS | 100.0 | 2.0 | 27.7 | 4.8 | 10.5 | 8.3 | 3.2 | 13.9 | 11.3 | 4.8 | 14.3 | 4.2 | 4.1 | 7.8 | 12.3 | 7.7 | 3.1 | 4.8 | 21.4 | 10.8 | 6.2 | 3.1 |
| MpMS | 2.0 | 100.0 | 4.9 | 6.6 | 17.0 | 10.8 | 3.3 | 4.9 | 11.5 | 6.7 | 7.1 | 10.2 | 10.4 | 7.3 | 11.5 | 8.2 | 11.5 | 12.9 | 6.9 | 24.6 | 3.3 | 8.2 |
| CoSS | 27.7 | 4.9 | 100.0 | 13.2 | 9.0 | 14.0 | 9.4 | 9.4 | 12.7 | 10.6 | 6.7 | 6.7 | 11.4 | 14.7 | 8.5 | 5.9 | 3.8 | 5.5 | 13.0 | 8.5 | 10.2 | 5.9 |
| CsSS | 4.8 | 6.6 | 13.2 | 100.0 | 7.8 | 4.7 | 12.0 | 8.7 | 18.0 | 2.2 | 8.1 | 1.7 | 12.7 | 13.4 | 7.6 | 8.8 | 9.8 | 16.7 | 13.2 | 9.8 | 12.0 | 6.6 |
| ZbSS | 10.5 | 17.0 | 9.0 | 7.8 | 100.0 | 26.2 | 9.1 | 9.1 | 6.3 | 9.5 | 9.1 | 14.6 | 4.4 | 11.3 | 9.2 | 12.7 | 7.9 | 9.4 | 17.7 | 9.2 | 3.5 | 9.3 |
| PfVS | 8.3 | 10.8 | 14.0 | 4.7 | 26.2 | 100.0 | 9.3 | 9.3 | 2.3 | 11.6 | 2.5 | 34.4 | 3.0 | 13.2 | 4.7 | 11.6 | 9.3 | 0.0 | 14.6 | 7.0 | 18.6 | 7.0 |
| NfSS | 3.2 | 3.3 | 9.4 | 12.0 | 9.1 | 9.3 | 100.0 | 7.6 | 6.3 | 5.0 | 7.6 | 1.7 | 8.5 | 5.9 | 6.8 | 9.1 | 10.7 | 20.8 | 5.8 | 7.6 | 8.8 | 6.9 |
| PbSS | 13.9 | 4.9 | 9.4 | 8.7 | 9.1 | 9.3 | 7.6 | 100.0 | 5.1 | 8.7 | 10.1 | 6.7 | 5.6 | 1.5 | 5.7 | 9.0 | 16.2 | 1.9 | 8.7 | 11.3 | 8.4 | 5.8 |
| PvPS | 11.3 | 11.5 | 12.7 | 18.0 | 6.3 | 2.3 | 6.3 | 5.1 | 100.0 | 3.9 | 6.9 | 6.7 | 15.0 | 11.8 | 7.6 | 3.8 | 7.7 | 11.9 | 10.1 | 10.1 | 10.1 | 8.9 |
| EvQS | 4.8 | 6.7 | 10.6 | 2.2 | 9.5 | 11.6 | 5.0 | 8.7 | 3.9 | 100.0 | 23.5 | 3.3 | 5.6 | 7.4 | 8.6 | 10.0 | 14.3 | 11.5 | 6.0 | 10.5 | 7.4 | 8.9 |
| CgDS | 14.3 | 7.1 | 6.7 | 8.1 | 9.1 | 2.5 | 7.6 | 10.1 | 6.9 | 23.5 | 100.0 | 5.4 | 8.3 | 3.2 | 7.1 | 13.0 | 13.8 | 11.5 | 9.5 | 12.4 | 10.1 | 12.8 |
| AcSS | 4.2 | 10.2 | 6.7 | 1.7 | 14.6 | 34.4 | 1.7 | 6.7 | 6.7 | 3.3 | 5.4 | 100.0 | 8.8 | 5.0 | 6.7 | 11.7 | 8.5 | 8.0 | 13.5 | 5.0 | 15.0 | 10.0 |
| FoFS | 4.1 | 10.4 | 11.4 | 12.7 | 4.4 | 3.0 | 8.5 | 5.6 | 15.0 | 5.6 | 8.3 | 8.8 | 100.0 | 6.7 | 8.5 | 10.0 | 19.7 | 11.5 | 10.0 | 8.5 | 9.9 | 7.0 |
| TpcA | 7.8 | 7.3 | 14.7 | 13.4 | 11.3 | 13.2 | 5.9 | 1.5 | 11.8 | 7.4 | 3.2 | 5.0 | 6.7 | 100.0 | 5.9 | 5.9 | 4.5 | 6.5 | 8.6 | 5.9 | 4.4 | 4.4 |
| FgMS | 12.3 | 11.5 | 8.5 | 7.6 | 9.2 | 4.7 | 6.8 | 5.7 | 7.6 | 8.6 | 7.1 | 6.7 | 8.5 | 5.9 | 100.0 | 12.5 | 8.8 | 5.6 | 8.7 | 9.4 | 10.3 | 7.8 |
| AcOS | 7.7 | 8.2 | 5.9 | 8.8 | 12.7 | 11.6 | 9.1 | 9.0 | 3.8 | 10.0 | 13.0 | 11.7 | 10.0 | 5.9 | 12.5 | 100.0 | 16.7 | 4.1 | 4.4 | 10.9 | 13.6 | 15.3 |
| PaPS | 3.1 | 11.5 | 3.8 | 9.8 | 7.9 | 9.3 | 10.7 | 16.2 | 7.7 | 14.3 | 13.8 | 8.5 | 19.7 | 4.5 | 8.8 | 16.7 | 100.0 | 3.7 | 4.4 | 15.1 | 7.9 | 13.7 |
| PrDS | 4.8 | 12.9 | 5.5 | 16.7 | 9.4 | 0.0 | 20.8 | 1.9 | 11.9 | 11.5 | 11.5 | 8.0 | 11.5 | 6.5 | 5.6 | 4.1 | 3.7 | 100.0 | 2.6 | 9.3 | 3.6 | 3.8 |
| PaFS | 21.4 | 6.9 | 13.0 | 13.2 | 17.7 | 14.6 | 5.8 | 8.7 | 10.1 | 6.0 | 9.5 | 13.5 | 10.0 | 8.6 | 8.7 | 4.4 | 4.4 | 2.6 | 100.0 | 5.8 | 11.6 | 10.1 |
| EvAS | 10.8 | 24.6 | 8.5 | 9.8 | 9.2 | 7.0 | 7.6 | 11.3 | 10.1 | 10.5 | 12.4 | 5.0 | 8.5 | 5.9 | 9.4 | 10.9 | 15.1 | 9.3 | 5.8 | 100.0 | 11.2 | 17.5 |
| TnDC | 6.2 | 3.3 | 10.2 | 12.0 | 3.5 | 18.6 | 8.8 | 8.4 | 10.1 | 7.4 | 10.1 | 15.0 | 9.9 | 4.4 | 10.3 | 13.6 | 7.9 | 3.6 | 11.6 | 11.2 | 100.0 | 10.7 |
| EvSS | 3.1 | 8.2 | 5.9 | 6.6 | 9.3 | 7.0 | 6.9 | 5.8 | 8.9 | 8.9 | 12.8 | 10.0 | 7.0 | 4.4 | 7.8 | 15.3 | 13.7 | 3.8 | 10.1 | 17.5 | 10.7 | 100.0 |

**Figure S23. Multiple sequence alignment matrix of the cyclase domain and linker region.** Cells are colored according to value; the more intense the blue color, the higher the sequence identity. The cyclase domains have lower identity than the prenyltransferase domains (see above). The linker regions have very little sequence identity.
