## Supplementary figures and images for "Structure of Bifunctional Variediene Synthase Yields Unique Insight on Biosynthetic Diterpene Assembly and Cyclization"

### Movie 1

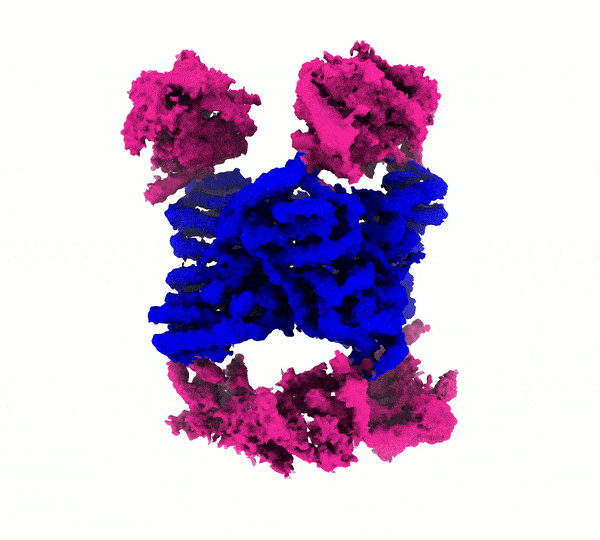

### Movie 2

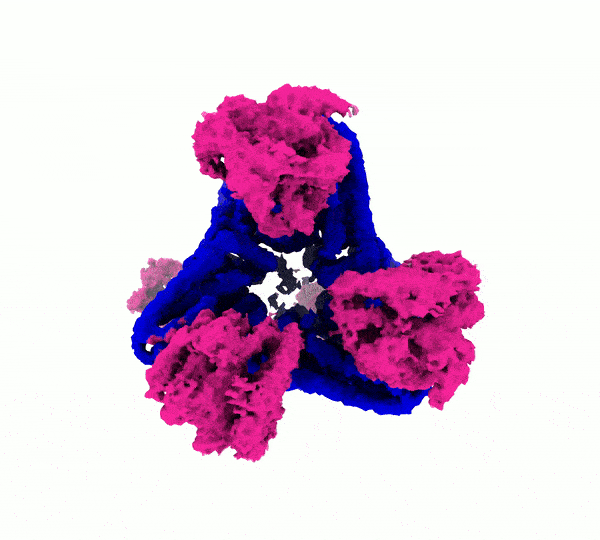
